# The cytokine structural homologs IL-1β and IL-1Ra have distinct dynamical character that potentially influence their roles in allosteric regulation of the IL-1 receptor

**DOI:** 10.64898/2026.09.10.750673

**Authors:** Glorisé Torres-Montalvo, Taylor R. Cole, Anthony C. Bishop, Rosana Lopes, A. Joshua Wand

## Abstract

Interleukin-1β (IL-1β) and the interleukin-1 receptor antagonist (IL-1Ra) share the β-trefoil fold, engage the same receptor (IL-1R), but produce opposite biological outcomes. IL-1β recruits the accessory protein IL-1RAcP to initiate inflammatory signaling while IL-1Ra occupies the receptor without supporting co-receptor recruitment. Seeking the origin of this divergence in properties not apparent from static structures, we used solution NMR relaxation to characterize fast internal motion in both proteins. Rigorous treatment of macromolecular tumbling shows both to be nearly maximally triaxial, which admits two near-degenerate axially symmetric solutions. This ambiguity was propagated throughout and found to have negligible impact. Complete replicate data sets from independently prepared IL-1β samples establish the experimental precision directly. Backbone amide and methyl-bearing side chain order parameters are indistinguishable between the two proteins, the latter averaging 0.484 ± 0.030 and 0.486 ± 0.033, indicating essentially equivalent residual conformational entropy. Their spatial distributions are uncorrelated when projected onto the shared fold. Importantly, the difference between the proteins is concentrated at the receptor interface, where methyl probes lying on the IL-1β footprint are considerably more mobile than those of IL-1Ra. It is therefore the agonist, not the antagonist, that presents an anomalously mobile binding surface. IL-1Ra also buries some 16% less surface while ostensibly binding IL-1R more tightly, the signature of an interface predisposed toward productive contact. Conformational entropy can thus distinguish protein function where structure and average flexibility are conserved, and the distinction resides not in how much entropy a protein has but in where it is kept.

---

Interleukin-1 (IL-1) signaling is a central inflammatory pathway that connects cellular injury, infection, and tissue stress to innate immune activation (1). As one of the most extensively studied members of the IL-1 cytokine family — which now comprises 11 ligands and 10 receptors — IL-1β has evolved to serve as a principal mediator of innate immunity, with evidence that even low doses of recombinant IL-1β can protect against lethal bacterial infection, for example (1). IL-1β, a pro-inflammatory agonist, binds IL-1R with low nanomolar affinity, recruiting the IL-1 receptor accessory protein (IL-1RAcP, also termed IL-1R2) to form an active ternary complex (2). This assembly gathers the intracellular Toll-IL-1-Receptor (TIR) domains of the receptor chains, initiating downstream inflammatory cascades, most notably NF-κB activation, that drive cytokine production (1, 3). Interleukin-1 receptor antagonist (IL-1Ra) binds IL-1R with even higher affinity (4) but fails to recruit IL-1RAcP thus rendering a structural mimic that imposes a non-signaling state (4–6). This distinction — agonism versus competitive antagonism at the same receptor — illustrates how subtle differences in receptor complex assembly generate fundamentally opposing biological outcomes.

The balance between IL-1β and IL-1Ra is physiologically critical. Insufficient IL-1β activity can compromise host defense, while excessive or persistent signaling contributes to autoinflammatory and chronic inflammatory disease, including type 2 diabetes, cardiovascular disease, and rheumatoid arthritis (7). Indeed, reduced IL-1Ra expression in pancreatic β-cells tilts the IL-1/IL-1Ra balance toward a pro-inflammatory state, impairing insulin secretion (7). The antagonistic function, combined with target specificity and minimal toxicity, has made IL-1Ra an attractive therapeutic target. The FDA-approved protein drug Anakinra, a non-glycosylated recombinant form of human IL-1Ra, has demonstrated clinical efficacy in treating rheumatoid arthritis in both juvenile and adult patients (6, 8).

IL-1β and IL-1Ra share the conserved β-trefoil fold characteristic of the IL-1 cytokine family, composed primarily of β-strands arranged into three related trefoil-like units. This structural conservation extends beyond IL-1β and IL-1Ra and is found in toxins, mitogens, and growth factors throughout nature, suggesting that this scaffold provides a versatile platform for diverse biological functions (2, 9). IL-1β and IL-1Ra are co-located within the human chromosome 2q13 gene cluster reflecting a common evolutionary ancestry. The interplay between IL-1β and IL-1Ra therefore provides a compelling framework for examining how structurally related cytokines, arising from the same primordial fold, generate profoundly distinct biological outcomes in vertebrate immunity (1, 10–12).

The IL-1 cytokine structural conservation and functional divergence suggests that evolution has acted not only on static receptor-contact residues but also on broader biophysical properties that are less apparent from static single structures alone (13). Molecular recognition is governed not only by shape complementarity and favorable intermolecular contacts, but also by the conformational states populated before binding and the entropy lost or redistributed during complex formation (14). The thermodynamics governing protein-ligand interactions involve a pallet of potential contributions (15, 16). Structural biology and theory have greatly informed the intermolecular interactions that comprise the changes in internal energy that help stabilize protein complexes. On the other hand, much less is known about the role of entropy in setting the free energy, with the changes in the entropy of water being perhaps the most familiar contribution. It has become clear that protein molecules populate an ensemble of states and variation of this residual entropy — the conformational entropy remaining in the folded state —potentially can significantly impact the thermodynamics of fundamental functions such as ligand binding (17, 18) and its natural extension allostery (19).

Here we employ solution NMR spectroscopy to characterize the dynamics of IL-1β and IL-1Ra. Recent developments in methodology allow for comprehensive, site resolved examination of fast internal motion in protein using NMR relaxation methods. These insights can now be interpreted in terms of quantitative measures of conformational entropy (18). Applying this perspective to IL-1β and IL-1Ra could reveal features of cytokine recognition that are not evident from static structures alone. Here we seek to not only contrast the internal motions of the agonist IL-1β and its counterpart antagonist IL-1Ra of the IL-1R receptor, but to also reveal how conformational entropy may be poised to fine tune their individual affinities for their common receptor and to alter the biological outcome.

## Results

### Resonance assignments

IL-1β has been widely characterized by solution NMR spectroscopy. Extensive backbone and side chain resonance assignments have been reported at various temperatures and pH (20–22). Triple resonance and total correlation spectra were acquired to map and extend these assignments to our sample conditions (pH 7.4, 25 °C). Backbone resonance assignments were effectively re-assigned *de novo* using BARASA (23) and agree well with those of Hommel et al. (22). Overall, 99.3% of the non-proline amide N-H and 98.4% of the methyl CH_3_ chemical shift correlations are confidently assigned.

Backbone resonance assignments of the C66A/C122A mutant interleukin-1 receptor antagonist (IL-1Ra) at 35 °C have been previously reported (23). Some resonance assignments for wild-type IL-1Ra have been reported previously (24). The wild-type IL-1Ra was also essentially assigned *de novo* using the same strategy as for IL-1β. Overall, 98.6% of the non-proline amide N-H and 98.6% of the methyl CH_3_ chemical shift correlations are confidently assigned. Formation of the C66-C122 disulfide bond was confirmed by chemical shift analysis (25). Resonance assignments have been deposited to the BMRB under accession codes 53840 and 53846 for IL-1Ra and 53841 and 53847 for IL-1β.

### Macromolecular tumbling and backbone dynamics

The main goal here is to quantitatively characterize the dynamics of methyl-bearing amino acid side chains in the IL-1R agonist IL-1β and its counterpart antagonist IL-1Ra. A critical element of the analysis is a rigorous characterization of the overall tumbling of each protein. Extensive NMR relaxation data was collected at multiple magnetic field strengths to determine backbone amide N-H bond vector 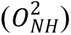 and methyl symmetry axis 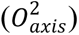 Lipari-Szabo model-free squared generalized order parameters (26) to high precision. Estimates of the precision of model-free order parameters is generally determined by Monte Carlo sampling of relaxation observables based on estimates of their precision obtained from fitting error (27). Here, however, direct knowledge of the precision will be crucial to a comparison of side chain motion of IL-1β and IL-1Ra.

To gain direct access to the experimental precision of obtained model free 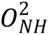 and 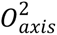 parameters, replicates of the relaxation data set for IL-1β were obtained using distinct samples (i.e., prepared separately). Fitted error of observables was used to generate 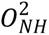 parameters while determining the appropriate tumbling model using a subset of relaxation data filtered for extensive local dynamics as indicated by a hetNOE that is more than 0.15 less than that predicted for an typical amide 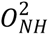 of 0.85 and/or the presence of evidence of chemical exchange effects (see Methods). The extensive backbone relaxation data sets allowed the rotational diffusion of both proteins to be characterized in detail (SI Appendix Tables S1-3).

For comparison, the mass-weighted heavy atoms of each structural model determined by crystallography give structural rhombicity of an equivalent inertia ellipsoid (ρ) of 0.421 and 0.468 for IL-1 β and IL-1Ra, respectively. In other words, both proteins have structural rhombicity near 0.5 and are nearly maximally triaxial. In this range no unique symmetry axis exists, and an axially symmetric model is anticipated to permit two near-degenerate solutions on near-orthogonal axes.

Exhaustive grid-searches for the best solution of the isotropic and axially symmetric tensor models indicates that the latter is required for both proteins. Isotropic tumbling is rejected decisively in every data set, while a fully anisotropic model is also not supported in any data set (IL-1β, p = 0.061 and 0.083; IL-1Ra, p = 0.106). However, as anticipated, the nature of the anisotropy is ill-determined. Extensive jackknife sampling of the relaxation data at the residue level predominately leads to oblate solutions but with a significant minority of prolate solutions also obtained. The 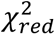 surface shows two minima separated by roughly 90 degrees (Table 1). Both oblate and prolate solutions improve on isotropic tumbling (for both basins of all three data sets, oblate F = 12.4–13.5, prolate F = 5.8–6.3, all p < 5 × 10⁻⁴). Importantly, the effective correlation time is robust to the oblate-prolate ambiguity: τc = 9.0 ± 0.1 ns for IL-1β and 8.5 ± 0.2 ns for IL-1Ra. Fitting of all three models to each data set is summarized in SI Appendix Table S4.

**Table 1.** Rotational diffusion tensors of IL-1β and IL-1Ra.

| Protein | Sample | Basin | D$\parallel$/10<sup>7</sup> s<sup>-1</sup> | D$\perp$/10<sup>7</sup> s<sup>-1</sup> | D$\parallel$/D$\perp$ | $\theta$ (°) | $\phi$ (°) | τc/ns | Nres/Nobs |
| --- | --- | --- | --- | --- | --- | --- | --- | --- | --- |
| IL-1β | 1 | oblate | 1.711 | 1.934 | 0.885 | 65.0 | 102.5 | 8.96 | 115 / 684 |
| IL-1β | 1 | prolate | 1.959 | 1.806 | 1.085 | 32.5 | 332.5 | 8.98 | 115 / 684 |
| IL-1β | 2 | oblate | 1.703 | 1.946 | 0.875 | 117.5 | 285.0 | 8.94 | 116 / 692 |
| IL-1β | 2 | prolate | 1.977 | 1.805 | 1.095 | 155.0 | 125.0 | 8.95 | 116 / 692 |
| IL-1Ra | 1 | oblate | 1.711 | 2.112 | 0.810 | 130.0 | 302.5 | 8.43 | 113 / 556 |
| IL-1Ra | 1 | prolate | 2.151 | 1.870 | 1.150 | 137.5 | 82.5 | 8.49 | 113 / 556 |

For IL-1β, the replicate 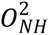 parameters and associated errors were used to generate the weighted average and error at all residues having sufficient data (n_obs_ ≥ 5) of acceptable quality. The replicates correlated with a squared Pearson coefficient (R^2^) of 0.81 and replicate precision (σ_rep) of 0.022 over the 133 residues determined in both samples (SI Appendix Fig. S1; Table S6). IL-1Ra was measured on a single production sample, and its 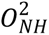 parameters are reported directly and are summarized in SI Appendix Table S7. The 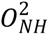 parameters are also deposited in the BMRB under accession numbers 53854 and 53857. Because whether the diffusion tensor is truly oblate or prolate is not determined, each order parameter carries a second, non-statistical uncertainty given by the separation between the two tensor solutions or basins (Δ_basin). This term is small for IL-1β (mean 0.006) but not far removed from the fitting error for IL-1Ra (mean 0.014). Depending on the geometric location of each amide NH within the structure, the shift varies and can change sign from residue to residue. However, this variation largely cancels on averaging and the mean 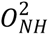 of either protein moves by no more than 0.003 between the two tensors. The tensor ambiguity therefore is negligible for the average and matters only for consideration of individual sites where it has generally quite small impact. Across the 114 amides satisfactorily fit in both preparations — whether or not an exchange term was required — the mean 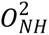 is 0.805 with a standard error of the mean (SEM) of 0.004. Applying the same criteria to the IL-1Ra data set gives 0.805 ± 0.005 over 112 amides, indistinguishable from IL-1β (Welch t = −0.001, p > 0.999); indeed the two IL-1β preparations differ from one another by more than IL-1β differs from IL-1Ra. Amides requiring an exchange term are not less ordered than those that do not (IL-1β, 0.799 ± 0.005 against 0.807 ± 0.005; IL-1Ra, 0.789 ± 0.011 against 0.810 ± 0.005) and are therefore pooled. Ninety-seven residues had determined 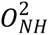 in both IL-1β samples and under both tumbling solutions and agree with an average deviation (*σ*) of 0.0165. Elimination of four outliers (G139, F112, Q116 and S125; Grubbs, α = 0.05) reduces *σ* to 0.0097. Monte Carlo sampling of the fitted observable errors gives a 〈*σ_MC_*〉 of 0.0065. However, each individual *σ_MC_* must be scaled to reflect the inadequacies of the model by multiplication by 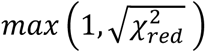, which leads to a 〈*σ*〉 of 0.0145. Propagating the scaled errors through the replicate average gives 0.0091. In summary, despite uncertainty in the symmetry of the tumbling tensors for both proteins, derived macromolecular tumbling correlation times and local 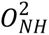 parameters are very well determined.

### Fast methyl-bearing side chain motion

Quantitative characterization of the tumbling parameters for both IL-1β and IL-1Ra sets the stage for an examination of the dynamics of their methyl-bearing side chains. The time dependence of intensities of single and multiple quantum cross-correlated relaxation profiles were used to obtain the relaxation parameters η and *δ* where η is the ratio of the cross correlated relaxation rates and *δ* is the contribution from dipolar relaxation (28). The tumbling models and associated correlation times determined above were then used to extract individual 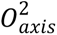 parameters. The replicate 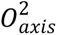 parameters and associated errors were combined to provide the weighted average and error for each methyl group. The derived model-free order parameters are listed in SI Appendix Tables S8-S10 (IL-1β) and S11 (IL-1Ra).

As a segue into the nature of fast motion of methyl-bearing side chains of proteins it is useful to first examine the distribution of 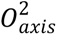 parameters. Numerous examples have indicated that methyl-bearing side chains exhibit a range of rotamer averaging (17, 29, 30). For soluble proteins, three classes of motion are often distinguished in histograms of 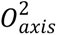 parameters: the so-called ω-class centered around 0.85 and arising from highly restricted motion within a single rotameric well; the *α*-class centered around 0.6 and arising from motion largely within a single rotameric well but with occasional excursions to other rotamers; and J-class centered around 0.35 and involving more extensive rotamer interconversion (29, 30). Recently, a fourth class of motion, termed J’, has been observed in integral membrane proteins and is characterized by extremely low 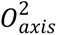 parameters centered around 0.2 (31–33). The J’-class of order parameters reflects extensive averaging about all *χ*-torsion angles (31).

The distributions of 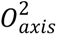 parameters across IL-1β and IL-1Ra are shown in Fig. 1. Assigning each methyl group to the nearest class center — that is, with boundaries at 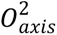 = 0.275, 0.475 and 0.725 — the two proteins populate the four classes of motion (29, 30) similarly: J’ 27% and 26%, J 24% and 21%, *α* 27% and 36%, and ω 22% and 17% for IL-1β and IL-1Ra respectively; a χ^2^ test of these binned populations finds the two distributions indistinguishable (χ^2^ = 1.3, p = 0.74). The comparison is insensitive to shifts of ±0.05 of the boundary positions. Both proteins clearly sample the full range of motional classes, including the J’-class recently revealed in integral membrane proteins. Strikingly, IL-1β and IL-1Ra have indistinguishable average methyl order parameters: 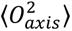 0.484 ± 0.030 and 0.486 ± 0.033, respectively. The two averages differ by 0.002 ± 0.045 (Welch’s t, p = 0.97).

**Fig. 1.**
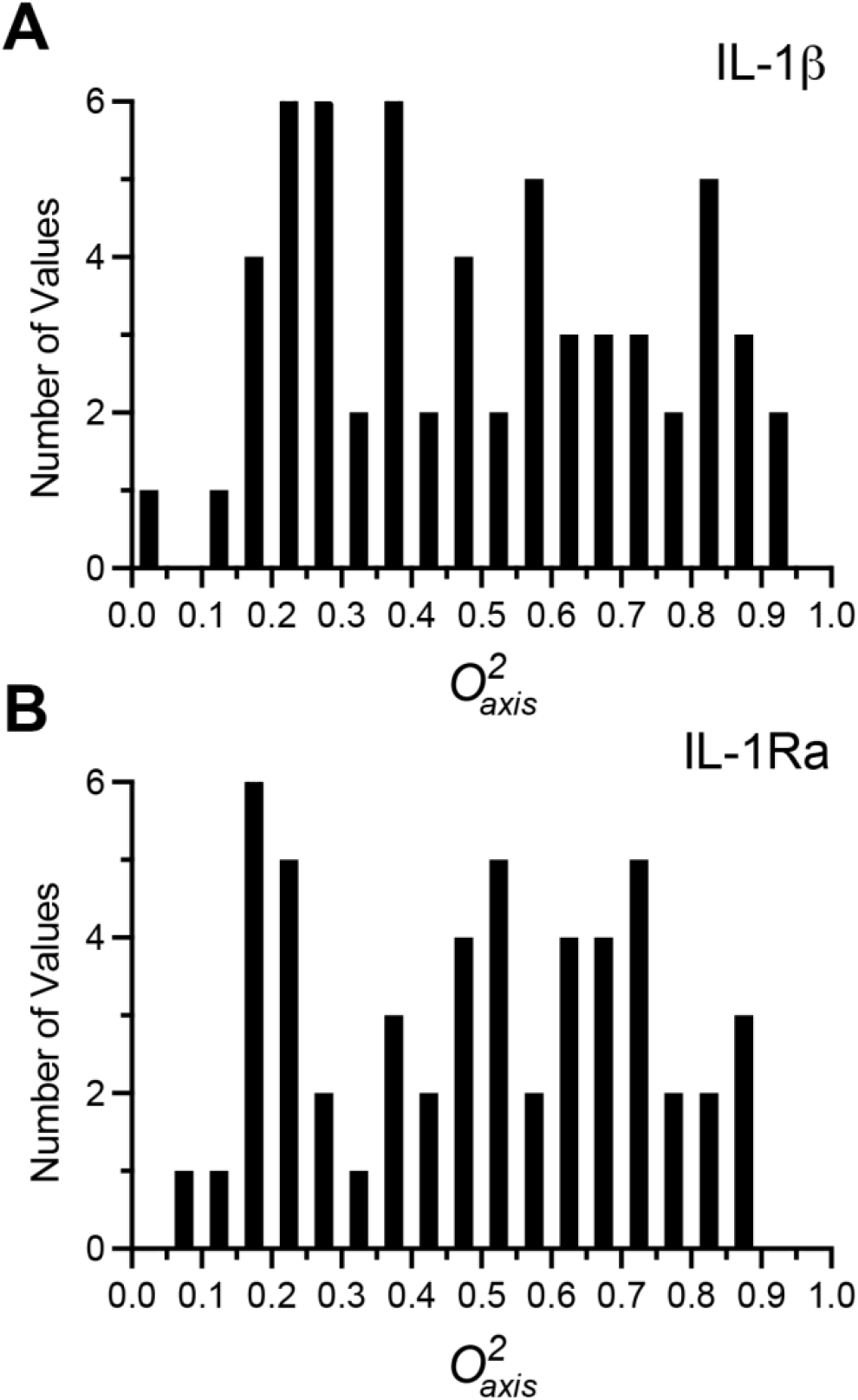
Statistical distributions of methyl-bearing side chain motion in IL-1β and IL-1Ra. Histograms of the determined O^2^*_axis_* parameters of IL-1β (n = 63) and IL-1Ra (n = 53).

The 95% confidence interval on the difference is ± 0.09. As noted above, both averages are insensitive to the undetermined nature (i.e., oblate vs prolate) of the rotational anisotropy, moving by less than 0.002 when every order parameter is re-evaluated under the alternative diffusion tensor. The close correspondence of methyl side chain motion of IL-1β and IL-1Ra, as quantified by the 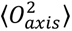, implies they have an essentially equivalent residual conformational entropy (31).

The IL-1β and IL-1Ra constructs used in this study are 153 and 152 residues, respectively, with 31% sequence identity and share a classic trefoil fold (Fig. 2). We examined the spatial distribution of methyl order parameters of each protein seeking clues to the origin of the equivalence of average dynamics. As with all proteins examined thus far by this approach, both proteins have a highly heterogeneous *spatial* distribution of side chain motion. Further, the spatial distributions appear to be quite different (Fig. 2).

**Fig. 2.**
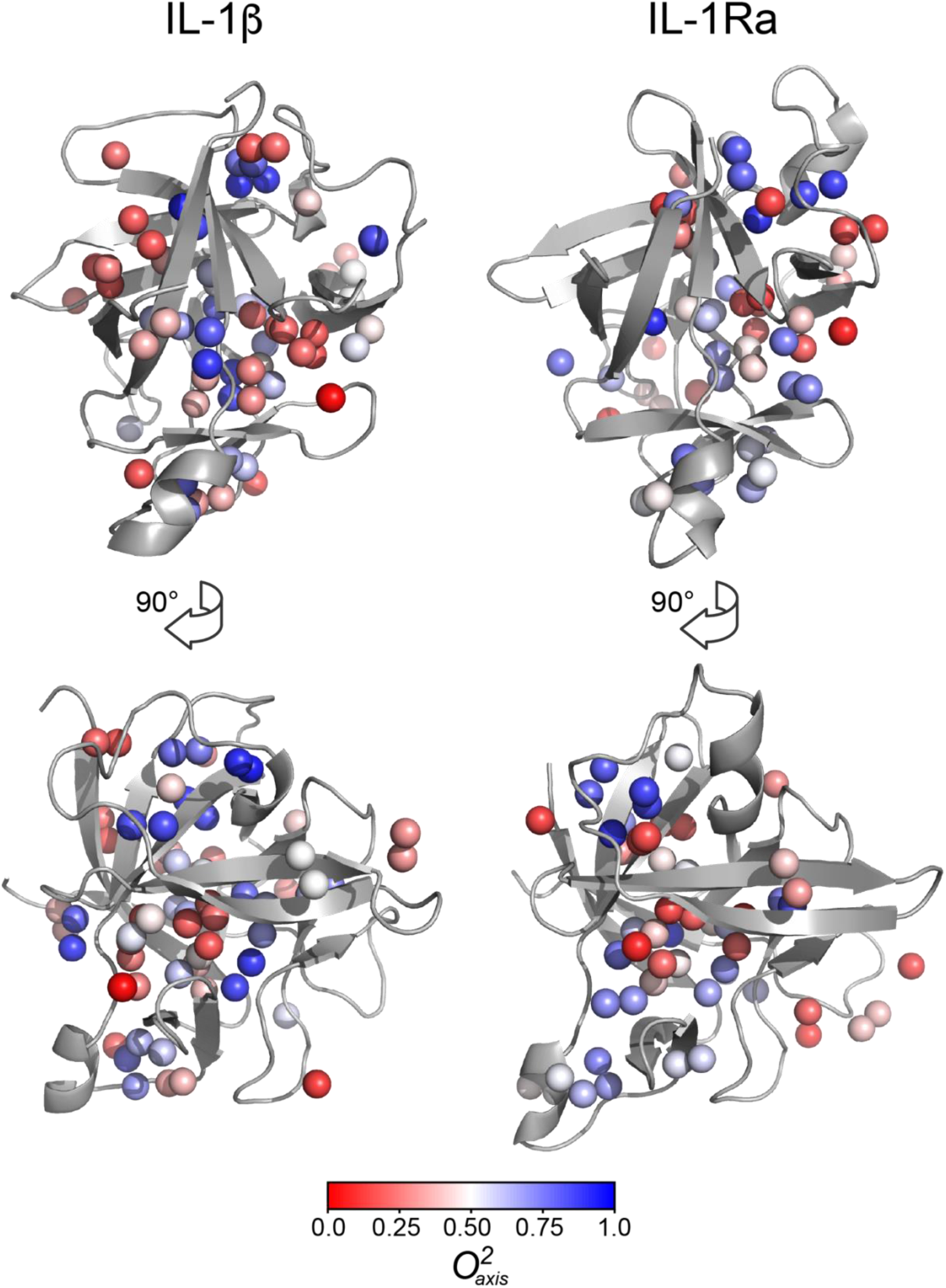
Structural distribution of methyl symmetry-axis order parameters in IL-1b and IL-1Ra. Cartoon representations of the crystal structures of IL-1β (PDB code: 9ILB, chain A) (34) and IL-1Ra (PDB code: 1ILR, chain 2) (35). The structures of IL-1β and IL-1Ra are highly similar, with an alpha carbon r.m.s.d. of 0.83 Å over the 98 of 135 pairs retained after outlier rejection. Methyl groups having determined 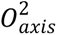 parameters are shown as spheres and are color coded as indicated by the inset color bar. Drawn with PyMol.

To put this on a quantitative footing, we created a spatial map of the O^2^*_axis_* parameters and computed at each alpha carbon a distance-weighted local average of the order parameters of all methyls within 10 Å (Fig. 3). The projection was calculated using the exponential distance dependence of observed perturbation by ligand binding in the Ras cdc42 protein (see Methods) (36). The resulting spatial distributions of IL-1β and IL-1Ra are largely uncorrelated though a simple linear regression gives a slightly negative slope of −0.144 ± 0.082 (Fig. 3).

**Fig. 3.**
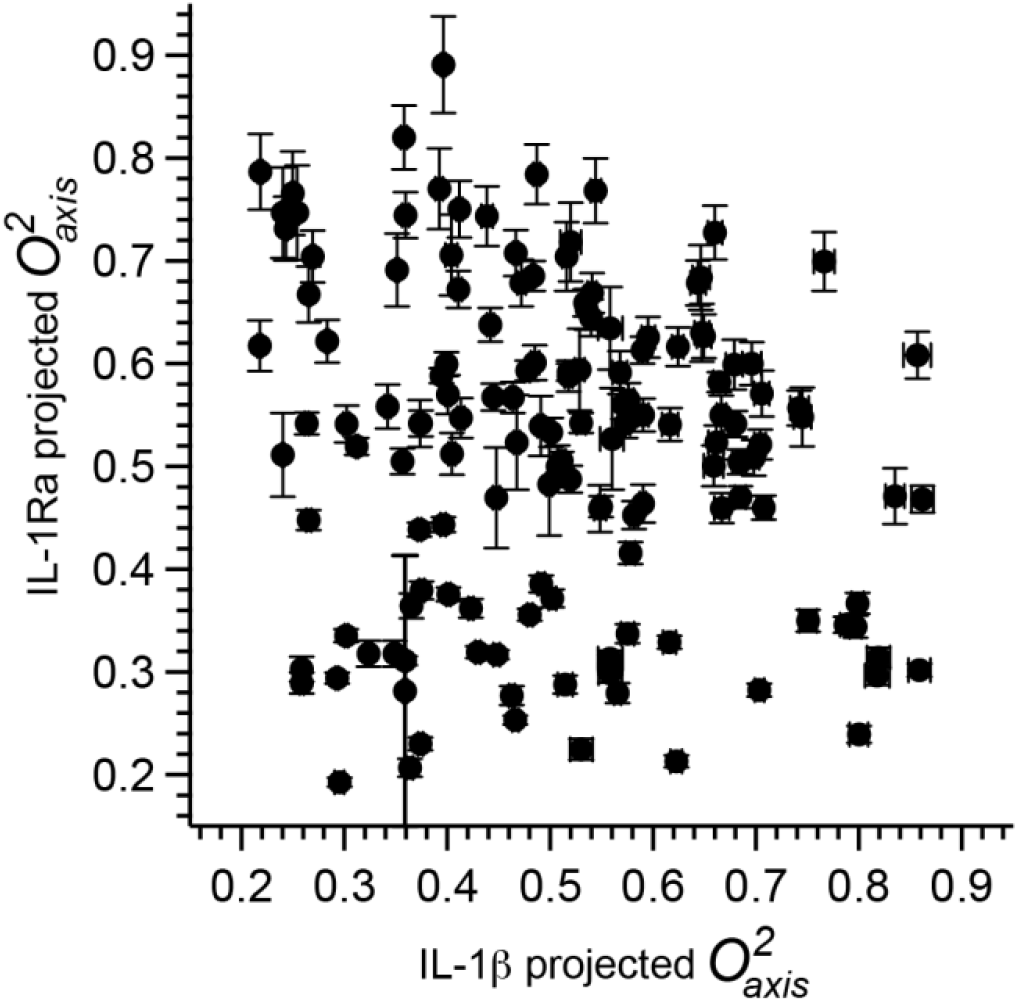
Spatially projected methyl dynamics along the shared β-trefoil fold of IL-1β and IL-1Ra. Statistical correlation of projections of the O^2^*_axis_* values to the alpha carbons of IL-1β and IL-1Ra at the 143 alpha carbons where the spatial dynamics fields are both defined. Each symbol represents a single alpha carbon. Errors of the projections were estimated by Monte Carlo sampling. For most cases, the error is less than the size of the symbol. The point cloud is centered on the shared mean and has a shallow negative slope (−0.14 ± 0.08) but without significant correlation (R^2^ = 0.021).

A comparison of the projected methyl dynamics as a function of the primary structure of the proteins reveals several interesting features (Fig. 4).

**Fig. 4.**
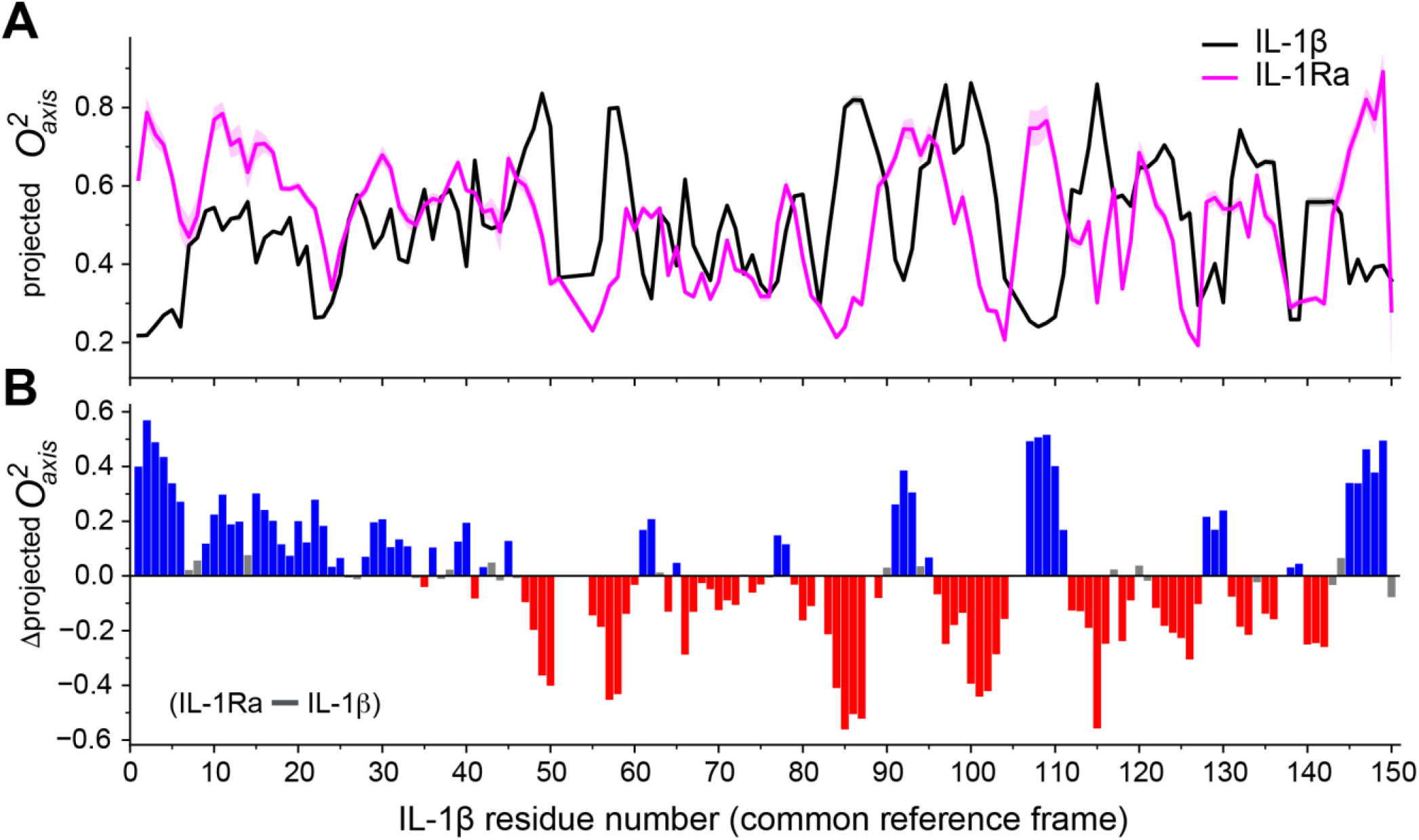
Sequence dependence of projected methyl dynamics along the shared β-trefoil fold of IL-1β and IL-1Ra. (A) The projections of the O^2^*_axis_* parameters to the Cα in Fig. 3 plotted versus residue number for IL-1β (black) and IL-1Ra (magenta). Lightly shaded region represent ± standard error derived by Monte Carlo sampling. (B) Difference (IL-1Ra minus IL-1β) in Cα projections of the O^2^ parameters. Blue, red and grey bars indicate regions where the rigidity of IL-1 β is less, more or not significantly different from IL-1Ra, respectively. IL-1Ra is significantly more rigid (higher O^2^axis projections) in the N-terminal region than IL-1β, suggesting entropic pre-organization for IL-1RAcP recruitment.

## Discussion

IL-1β and IL-1Ra share the β-trefoil fold and bind IL-1RI with high affinity, yet produce opposite biological outcomes: IL-1β recruits IL-1RAcP and initiates inflammatory signaling, while IL-1Ra occupies the receptor without supporting co-receptor recruitment. The central thrust of this study was to employ NMR relaxation to characterize the potential role of conformational entropy in the thermodynamics of molecular recognition by these two opposing allosteric regulatory cytokines. The approach rests on determining the underlying macromolecular reorientation tensor. In this respect, both proteins are close to maximally triaxial. The rhombicity of the equivalent inertia ellipsoid, computed from coordinates alone is ρ = 0.421 for IL-1β and 0.468 for IL-1Ra. In this regime no unique symmetry axis exists and an axially symmetric treatment necessarily admits two near-degenerate solutions on near-orthogonal axes. That is what is observed experimentally: roughly orthogonal oblate and prolate axial tensor solutions describe each data set almost equally well with oblate solutions being represented in the majority of resampled data subsets for both proteins (Table 1, SI Appendix Table S6). Though the crystal structures of both proteins suggest a subtle prolate symmetry, the tendency toward oblate solutions in the jackknife resampling of the data sets can likely be accommodated through dynamical behavior of the extensive network of connecting loops, which account for roughly half of the residues of the β-trefoil fold.

The precision of the data was explicitly examined through comparison of two independently prepared IL-1β samples, which gave a measured deviation in 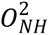 of 0.01 after elimination of four outliers (G139, F112, Q116 and S125). This represents on the order of 1% of a typical order parameter and also calibrates the error estimates conventionally obtained through Monte Carlo analysis. As is the common approach, relaxation in IL-1Ra was measured on a single production sample and the precision of backbone order parameters is estimated in the usual way through Monte Carlo sampling guided by replicate error of individual time points within a single relaxation time course for T_1_ and T_2_ measurements or signal-to-noise estimates in the case of the heteronuclear NOE.

Fortunately, the apparent ambiguity of tumbling tensor matters little to the overall goal. A single axial fit returns a well-conditioned tensor and gives no indication that a second, equally good solution exists on another axis. The consequences were therefore propagated. The effective correlation times of both proteins were found to be essentially basin-invariant, the two solutions differing by 0.013 ns for IL-1β and 0.062 ns for IL-1Ra, respectively eight-fold and three-fold smaller than the quoted uncertainties. This arises because the basins differ in how a fixed amount of anisotropy is oriented rather than in the isotropic part of the tumbling tensor. The obtained 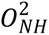 parameters are similarly robust to the ambiguity in the relatively mild axial symmetry: re-evaluating every methyl under the alternative tensor moves the mean by less than 0.002 in either protein.

Remarkably, the two proteins carry indistinguishable average methyl-bearing side chain motion with 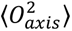 of 0.484 ± 0.030 and 0.486 ± 0.033, a statistically insignificant difference of 0.002 ± 0.045 (Welch’s t, p = 0.97).The NMR-based entropy meter therefore holds them essentially as effectively equivalent resevoirs of residual conformational entropy. The statistical distribution of methyl-bearing side chain motion of both proteins is unremarkable and differ in detail only (Fig. 1). What differs greatly is where that entropy is kept. Projected onto the shared fold, the two distributions of dynamics are uncorrelated (Fig. 3). Regions that are ordered in one protein are, in general, not the regions that are ordered in the other. It is noted that the spatial differences reported here are considerably larger than the variability attributable to sample preparation and measurement, and are taken to reflect genuine differences between the solution ensembles.

Assembly of the signaling complex engages two distinct surfaces of the cytokine. Site A is the central binding pocket that engages IL-1RI and is conserved between the two cytokines; Site B is a separate surface that contacts IL-1RAcP and is required for co-receptor recruitment in agonist complexes (4, 6, 38). The two surfaces barely overlap. In the ternary signaling complex (PDB code 4DEP) Site B comprises twenty residues of IL-1β that bury 657 Å^2^ against IL-1RAcP, of which only seven also contact IL-1RI, together contributing just 117 Å^2^ of the 1981 Å^2^ IL-1RI interface (Fig. 5). IL-1Ra lacks a functionally equivalent Site B. The dynamics of these surfaces differ markedly, and in the direction opposite to what might usually be assumed. Methyl probes lying on the receptor footprint have an 〈*O*^2^ 〉 of 0.325 in IL-1β but 0.495 in IL-1Ra, a difference of 0.17 ± 0.10, in proteins whose whole-molecule averages are indistinguishable. Measured against each protein’s own mean, the IL-1Ra footprint is unexceptional (0.495 against 0.485) while the IL-1β footprint departs sharply from the rest of its structure (0.325 against 0.484). The same asymmetry appears in the alpha carbon projected fields: the IL-1β footprint is significantly more mobile than the remainder of that protein (0.422 against 0.538, p < 0.001), and robust to how the footprint is defined). In contrast, the IL-1Ra footprint is not distinguishable from its own average (0.553 against 0.511, p = 0.089). Site B is likewise significantly more mobile than the remainder of IL-1β (0.442 against 0.557, p = 0.005 across seventeen alpha carbons) and is indistinguishable from Site A in this respect (p = 0.59), so both of the surfaces that the agonist presents to the receptor complex are anomalously mobile. It is noted, however, that only three of the sixty-three methyl probes lie directly on Site B, so this comparison rests on the projected field rather than on probes at the site itself. Perhaps surprisingly, it is the agonist, not the antagonist, that presents an anomalously mobile binding surface. Interestingly, IL-1Ra also buries less surface against the receptor (1656 Å^2^ against 1981 Å^2^, roughly 16% less) while ostensibly binding IL-1RI more tightly. A smaller, already-ordered interface that achieves equal or better affinity is the signature of a surface predisposed toward productive contact: it can form better interactions per unit of buried area while surrendering less conformational entropy on binding.

**Fig. 5.**
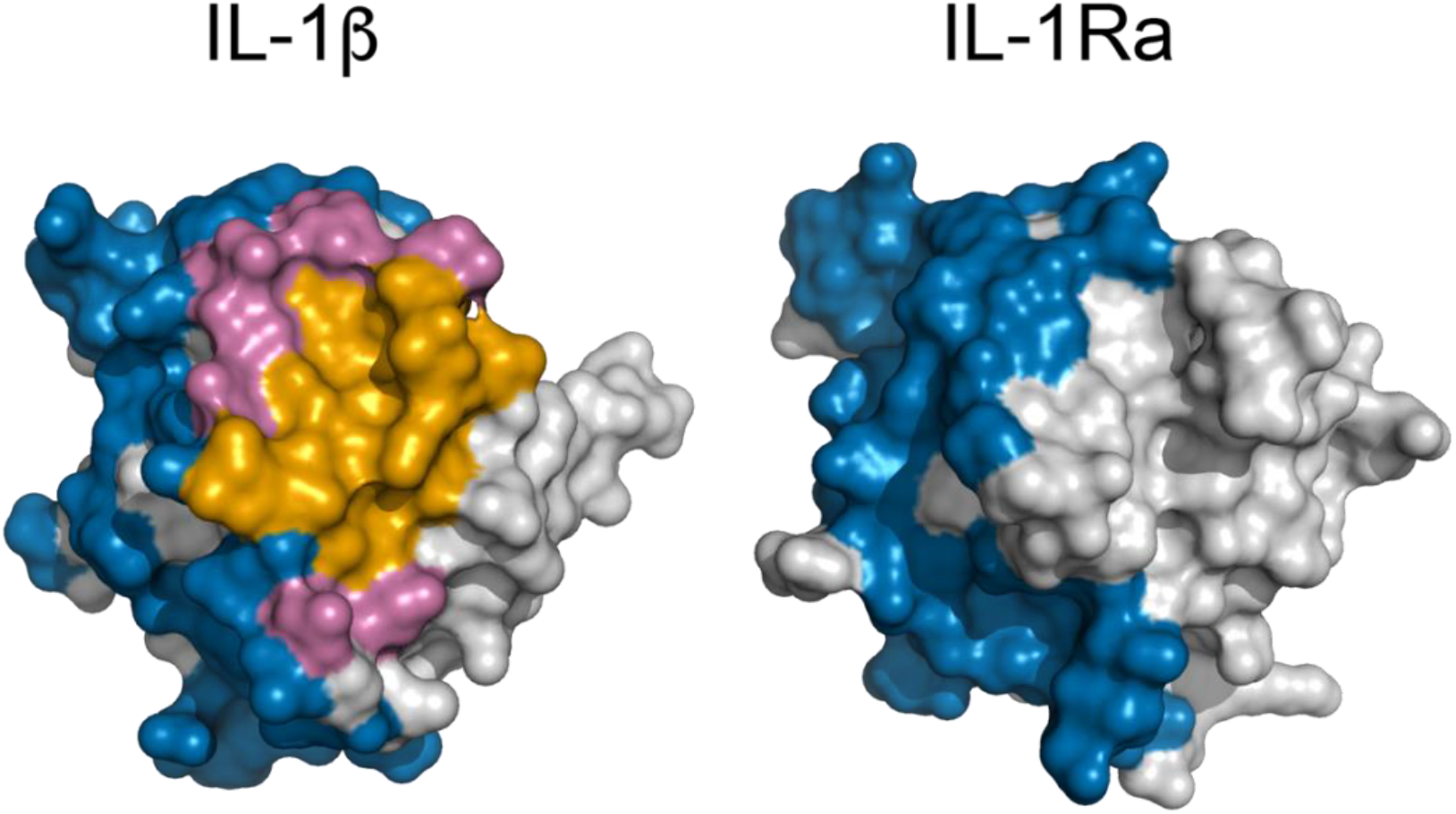
IL-1RI receptor-engaging surfaces of IL-1β and IL-1Ra. Molecular surface representations in a common orientation of IL-1β and IL-1Ra. IL-1β residues are colored by the receptor chain they contact in the assembled signaling complex: IL-1RI only (40 residues; blue), IL-1RAcP only (13 residues, orange), or both (7 residues; pink). IL-1Ra is shown in the same orientation with its IL-1RI contact surface colored equivalently; it forms no ternary complex and therefore presents no IL-1RAcP surface. Residues not in contact with the receptor are colored grey. Contacts are defined by buried solvent accessible surface area computed from PDB entries 1ITB (37), 1IRA (5) and 4DEP (4).

Changes in methyl-bearing side chain motion quantified by the 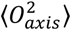 have been successfully linearly and quantitatively correlated with corresponding changes in conformational entropy. Applying this so-called NMR-based conformational entropy meter to the interface differences, and assuming both interfaces reach comparable rigidity in the complex, IL-1Ra surrenders approximately 25 kJ mol^−1^ less conformational entropy than IL-1β on binding. The value is offered to establish scale rather than as a measurement of the binding penalty. Conformational entropy is often a large determinant of the thermodynamics of association: in approximately one quarter of the complexes used to calibrate the meter, removing its contribution would leave affinities biologically ineffective (19). A redistribution of this magnitude is therefore large enough to affect receptor engagement materially. That possibility motivates investigation of motion within the IL-1R complexes, which is in progress.

IL-1β and IL-1Ra diverged functionally without major change to the β-trefoil scaffold. The two proteins hold the same total residual conformational entropy while distributing it in distinct ways. so the total appears constrained while its spatial allocation is not, and redistribution is available as an evolutionary variable at no cost to the overall budget. Since the structures superimpose to 0.83 Å over the 98 of 135 alpha carbon pairs retained after outlier rejection, dynamics has diverged considerably further than structure: selection can act on the dynamic character of a binding surface without redesigning the scaffold that presents it. The redistribution is not uniform over the fold but concentrates at the receptor footprint.

The direction of the change can apparently be assigned from the phylogeny. The IL-1 ligands arose by successive duplication of a prototypical proto-IL-1β gene that appeared alongside proto-IL-1R near the emergence of the vertebrates, some 420 million years ago, with IL-1Ra arising from one of the earliest of these duplications (10). The agonist configuration is therefore ancestral and the comparatively rigid, pre-organized IL-1Ra interface is the derived state. IL-1Ra has evolved under strong selection for specificity (10). In this view, antagonism was acquired not by the loss of a binding site but by the stiffening of one where pre-organization of the components of the interface provided by IL-1Ra helps avoid an entropic penalty upon binding.

Current IL-1-targeted therapeutics, including Anakinra, work by blocking receptor activation through competitive binding. The present data suggest the dynamic properties of the Site B interface as an additional target. Site B must change conformation for the co-receptor to dock: between the binary and ternary complexes residues 54, 106 and 107 move by 1.0, 2.4 and 3.5 Å respectively, against a 0.25 Å reproducibility between the two crystallographically independent copies of the ternary complex. A surface that is anomalously mobile in the free state and must rearrange on assembly is one whose flexibility is functionally required. A molecule that rigidifies the Site B region of IL-1β in its free-state conformation will therefore suppress co-receptor recruitment without preventing IL-1RI binding, separating occupancy from activation. This is the same manoeuvre evolution appears to have used in deriving IL-1Ra, and it is a strategy distinct from a simple competitive blockade.

In conclusion, IL-1β and IL-1Ra hold similar average conformational entropy but house it in different parts of a common fold, and the differences are concentrated at the surfaces that engage the receptor. The agonist presents a mobile binding surface and buries more of it; the antagonist presents a pre-organized one, buries less, and binds at least as tightly. Conformational entropy can therefore distinguish protein function where structure and average flexibility are conserved, and the distinction resides not in how much entropy a protein has but in where it is kept.

## Materials and Methods

### Interleukin-1 Receptor Antagonist (IL-1Ra) and Interleukin-1β (IL-1β)

Isotopically labeled wt-IL-1Ra (UniProt code: P15810) and wt-IL-1β (UniProt code: P01584) were prepared as described previously (23, 38) with the following labeling schemes: uniformly ^15^N,^13^C-labled proteins for triple resonance assignment experiments; trace ^13^C-labeled proteins for assignment of prochiral Leu and Val methyl groups (39); uniformly labeled ^15^N-proteins for backbone ^15^N-relaxation experiments; and selectively ^13^C^1^H_3_-methyl, ^15^N-^1^H-labeled proteins in an otherwise perdeuterated background (> 95% D) for methyl-group cross-correlated ^1^H-^1^H relaxation and ^15^N-relaxation, respectively. Selective methyl labeling employed the 2-keto-3,3-d_2_-4-^13^C-butyrate/2-keto-3-methyl-^13^C-3-d_1_-4-^13^C-butyrate ^13^CH_3_ labeling strategy (40) supplemented with non-deuterated ^13^C_ε_-labeled methionine and expressed during growth on ^15^NH_4_Cl as a sole nitrogen source. Proteins were generally prepared for NMR spectroscopy in 100 mM NaCl, 25 mM HEPES, pH 7.4, 100 μM DSS, 10% D_2_O, and 0.02% (w/v) NaN_3_ at a concentration of 1 mM and 0.4 mM of IL-1Ra and IL-1β, respectively.

### NMR Spectroscopy

Non-uniformly sampled (NUS) (41) triple resonance assignment experiments were acquired on a four-channel Bruker NEO 800 MHz (^1^H) NMR spectrometer. ^13^C-HMQC, HCCH-TOCSY (42) and methyl-TOCSY (43) experiments were acquired on a four-channel Bruker Avance III HD 600 MHz (^1^H) NMR spectrometer. ^15^N-resolved NOESY experiments were acquired on a Bruker Avance III 500 MHz (^1^H) NMR spectrometer. All NMR spectrometers were equipped with cryogenically He cooled probes. All spectra were obtained at 298 K. NMR data was processed using NMRPipe (44) and visualized with NMRFAM-Sparky (45). NUS data was pre-processed with istHMS (41). Most processing was done on the NMRbox virtual platform (46). Resonance assignments were supported by the BARASA package (23).

BARASA was run in the NMRbox virtual platform with predicted chemical shifts generated by ShiftX2 (8) using 1IRA (chain X) as the pdb model for IL-1Ra and 9ILB for IL-1β. The chemical shifts statistics file was extracted from BMRB. In both cases, previously referenced CBCA(CO)NH, HNCACB and ^15^N HSQC peak list files were added to the data folder used by BARASA. Matching tolerances in the peak lists of 0.3, 0.3, 0.02 ppm were used for carbon, nitrogen and hydrogen dimensions respectively. One hundred runs were used. Default values were used for other parameters.

### Model free parameters

Lipari-Szabo model-free (26) squared generalized order parameters and effective correlation times were determined using in-house TUMBLE and RESTLESS software and employed an explicit grid search approach (27). Optimal tumbling models and rotational correlation times were determined (47) for each sample using combinations of ^15^N-relaxation data and crystal structures of IL-1β (PDB code: 9ILB, chain A) and IL-1Ra (PDB code: 1ILR, chain 2). For the tumbling analysis, each residue-level set was screened for chemical exchange effects using an orientation-aware exchange screen, in which residues with elevated R2 are identified iteratively at the fitted tensor rather than from a global R1·R2 statistic (53), with a screen width of 2.0 SD. Residues with inordinately low heteronuclear NOE were also excluded, using a τc- and field-dependent threshold determined automatically rather than a fixed cut-off, since a fixed threshold removes rigid residues at slow tumbling. The choice of screening rule is a dominant systematic in this analysis — on IL-1β sample 2 it moves τc by 0.100 ns and the unique axis by ∼20°, and on IL-1Ra the two screens give unique axes 40.9° apart — and is reported for that reason. An (48)^15^N chemical shift tensor breadth of 170 ppm and an effective N-H bond length (49) of 1.02 Å were employed. This bond length does not consider motion of the hydrogen about the N-H bond. Amide NH squared generalized order parameters 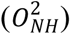 were determined using the determined tumbling model. Methyl order parameters were obtained by fitting the ratio of maximum intensities of individual methyl cross-peaks in the multiple-quantum and single-quantum relaxation experiments as a function of delay times (28). Fits had an R^2^ value above 0.9. Methyl symmetry axis squared generalized order parameters 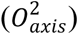 were derived from determined raw order parameters (O^2^) under the assumption of tetrahedral geometry (O^2^/0.111). Monte Carlo sampling of estimated error in relaxation observables was used to generate initial estimates of standard error of derived methyl order parameters.

### Precision of model free order parameters

True replicates (i.e., experiments based on separately prepared samples) provided two independent paired sets of 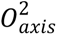 each with a standard error obtained by analytic propagation of the fitted error in η; there is no Monte Carlo step on the methyl pathway, and because a single η is measured per methyl group there is no per-methyl goodness-of-fit. These paired sets were then used to generate the weighted mean and standard error (n = 2) of 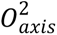 for each individual methyl group. Comparison of the two proteins, by contrast, used the unweighted population mean of each set with its standard error of the mean. Inverse-variance weighting is appropriate for combining two measurements of the same methyl group, where both estimate one quantity, but not for averaging across different sites: the uncertainty in an individual 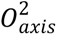 correlates with its value (r = 0.87 for IL-1β), so a precision-weighted average across methyl groups is drawn toward the least ordered sites and does not estimate the mean order parameter of the protein. Comparison of sets of 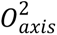 parameters from two different origins (e.g., different proteins) often involves unequal sizes. For two sets A & B of independently obtained 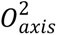 parameters we have:

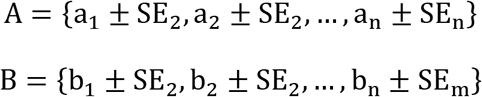

Using the weighted means, 〈A〉 and 〈B〉, and weighted standard errors, SE_A_ and SE_B_, we employ the Welch’s t-test for two independent samples with known standard errors and compare the weighted means of the unpaired sets A and B:

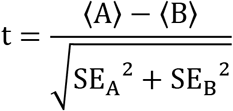

The Welch-Satterthwaite formula for the degrees of freedom is:

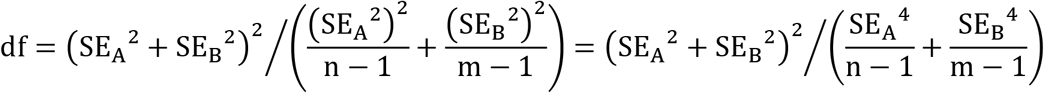

### Spatial analysis of model free order parameters

The crystal structures of IL-1Ra and IL-1β superimpose to an r.m.s.d. of 0.83 Å over the 98 of 135 alpha carbon pairs retained after iterative outlier rejection (PyMOL super, five cycles, 2 Å cutoff). The projection described below uses its own sequence-seeded fit, retaining 136 pairs within 4 Å at an r.m.s.d. of 1.4 Å. Due to the close packing of protein structures, local methyl-bearing side chain ps-ns dynamics measured by NMR relaxation are coupled, albeit weakly, to the motion of their neighbors (18, 50). To create a spatial field of the 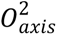 parameter, measured order parameters were projected using the exponential relationship of motional perturbation by ligand binding described by Moorman et al. (36). The normalized projection at a given alpha carbon is then:

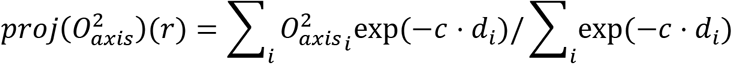

where *r* is the Cα coordinate, *r_i_* is the coordinate of the methyl probe carbon, and *d_i_* is the distance between them (*d_i_* = ‖*r* − *r_i_*‖) and *c* is 0.568 Å^-1^. The sums run over all methyl carbons within 10 Å of the alpha carbon.

### Miscellaneous structural and statistical analyses

Solvent accessible surface area was determined using the BioPython package using the crystal structures of free IL-1Ra and IL-1β and their complexes with the IL-1R (PDB codes 1IRA and 1ITB, respectively). Site B was defined as the IL-1β residues burying surface against the IL-1RAcP chain of the ternary signaling complex (PDB code 4DEP); both crystallographically independent copies of that complex give the same twenty residues. Displacements between the free, binary and ternary states were measured after superposition on the twelve β-strands of IL-1β. Superpositions employed PyMol (53). Statistical quantities were calculated using the SciPy package (51).

## Author Contributions

G.T-M. and A.J.W. designed research; G.T.-M. and R.L. prepared materials; G.T-M. and T.R.C. carried out NMR spectroscopy; G.T-M., T.R.C., A.C.B. and A.J.W. analyzed data; G.T-M. and A.J.W. wrote the paper.

Authors declare no competing interest.

## ACKNOWLEDGEMENTS

We thank Dr. J. A. Caro for helpful discussion during the early phase of this project. This work was supported by the NIH through grants R01 GM129076 and R35 GM158127 to A.J.W. and by Texas A&M University.

---

**Figure S1.**
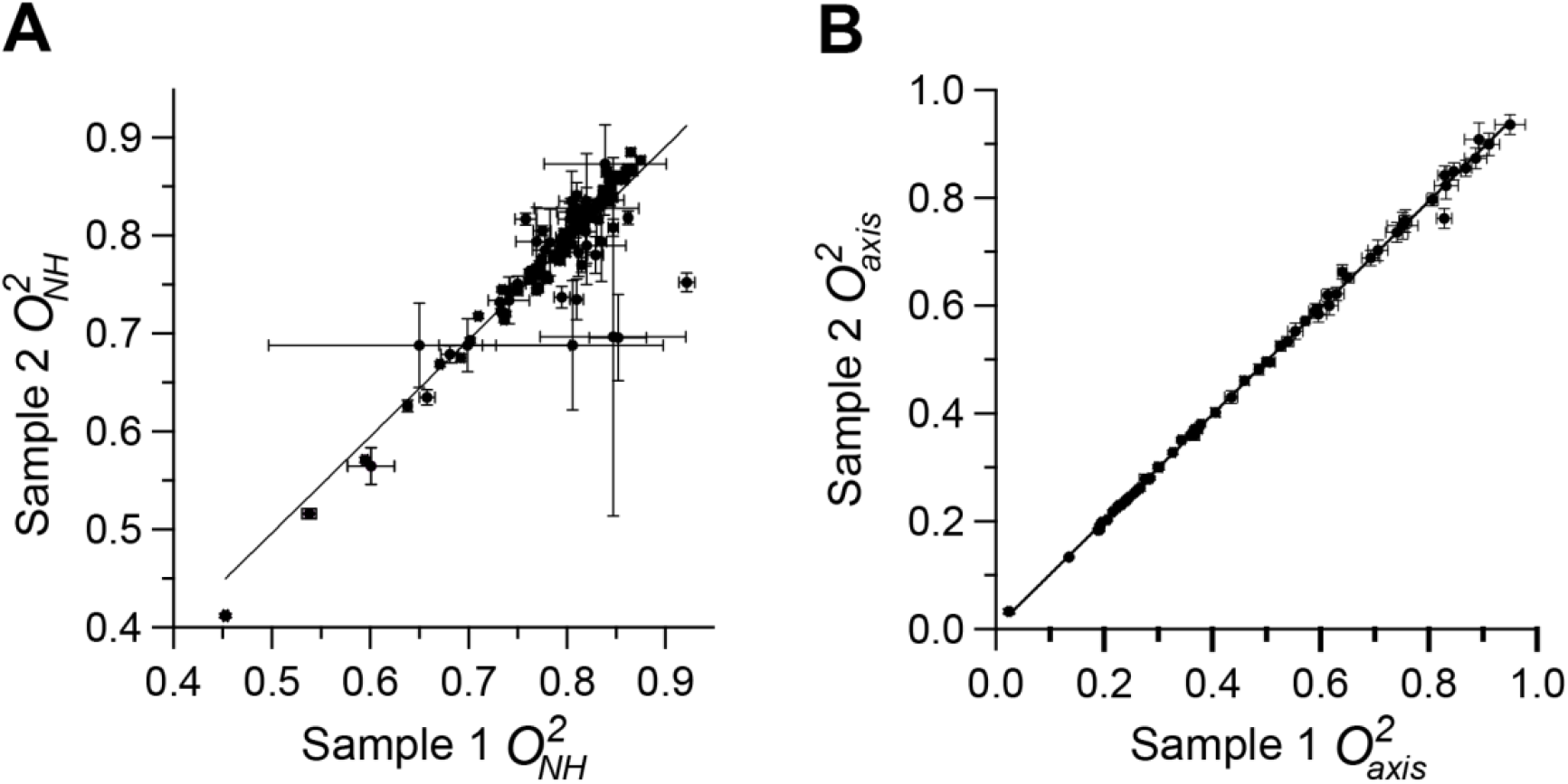
Correlation of model-free squared generalized order parameters from replicate IL-1β samples. Correlation of 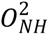 (A) and 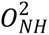 from two independently prepared samples of IL-1β. Each symbol is a single NH group; errors were estimated by Monte Carlo sampling and the dashed line has a slope of unity. Over the 133 residues determined in both samples, y = 0.99x + 0.00 with R^2^ = 0.81. Outliers were identified using the ROUT method (1) and the trimmed statistics are given with the figure. IL-1Ra is not shown as it has a single production sample and so R₂ cannot be decomposed into the intrinsic R₂ and Rex in a similar fashion.

**Table S1.**
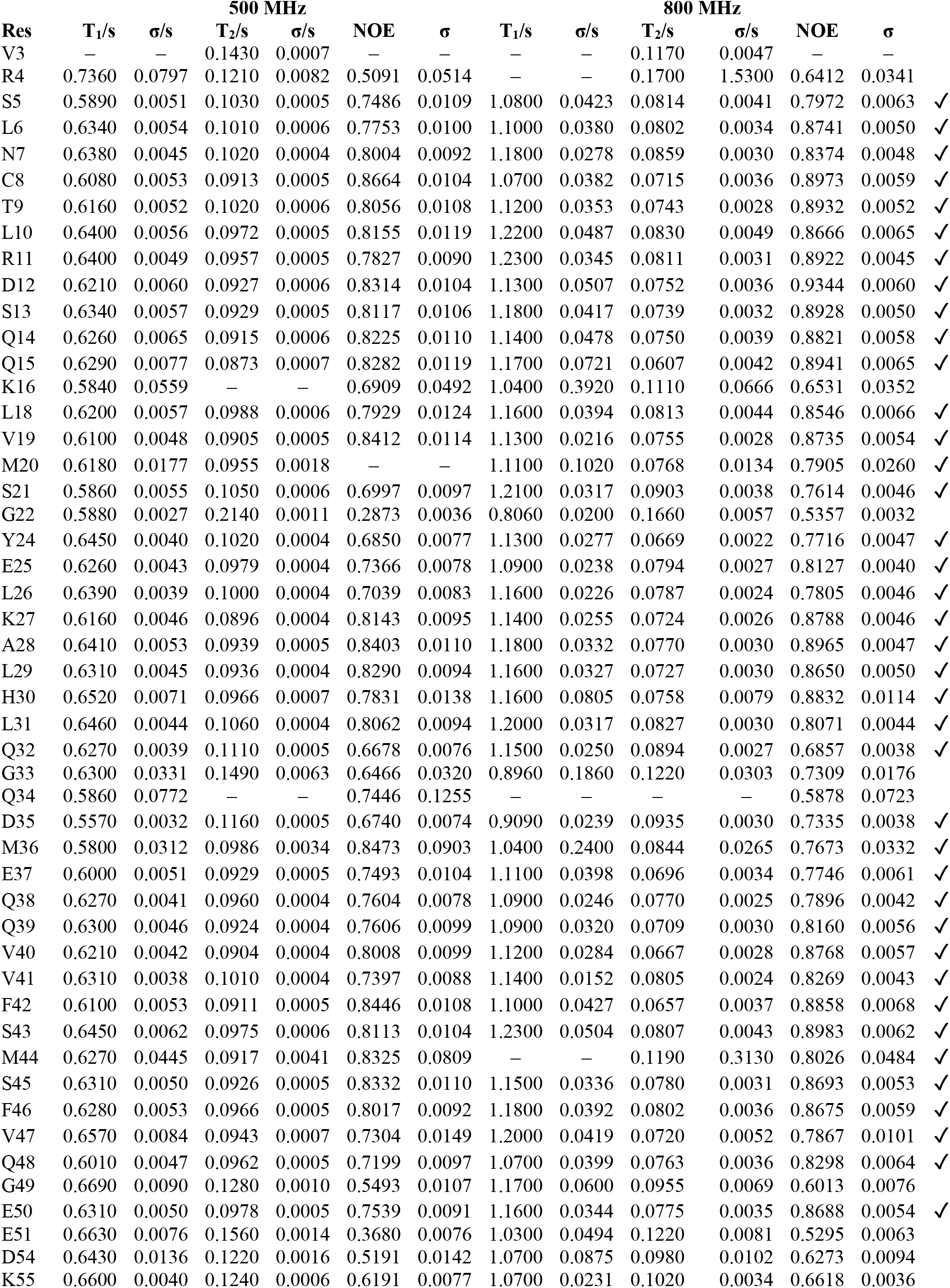
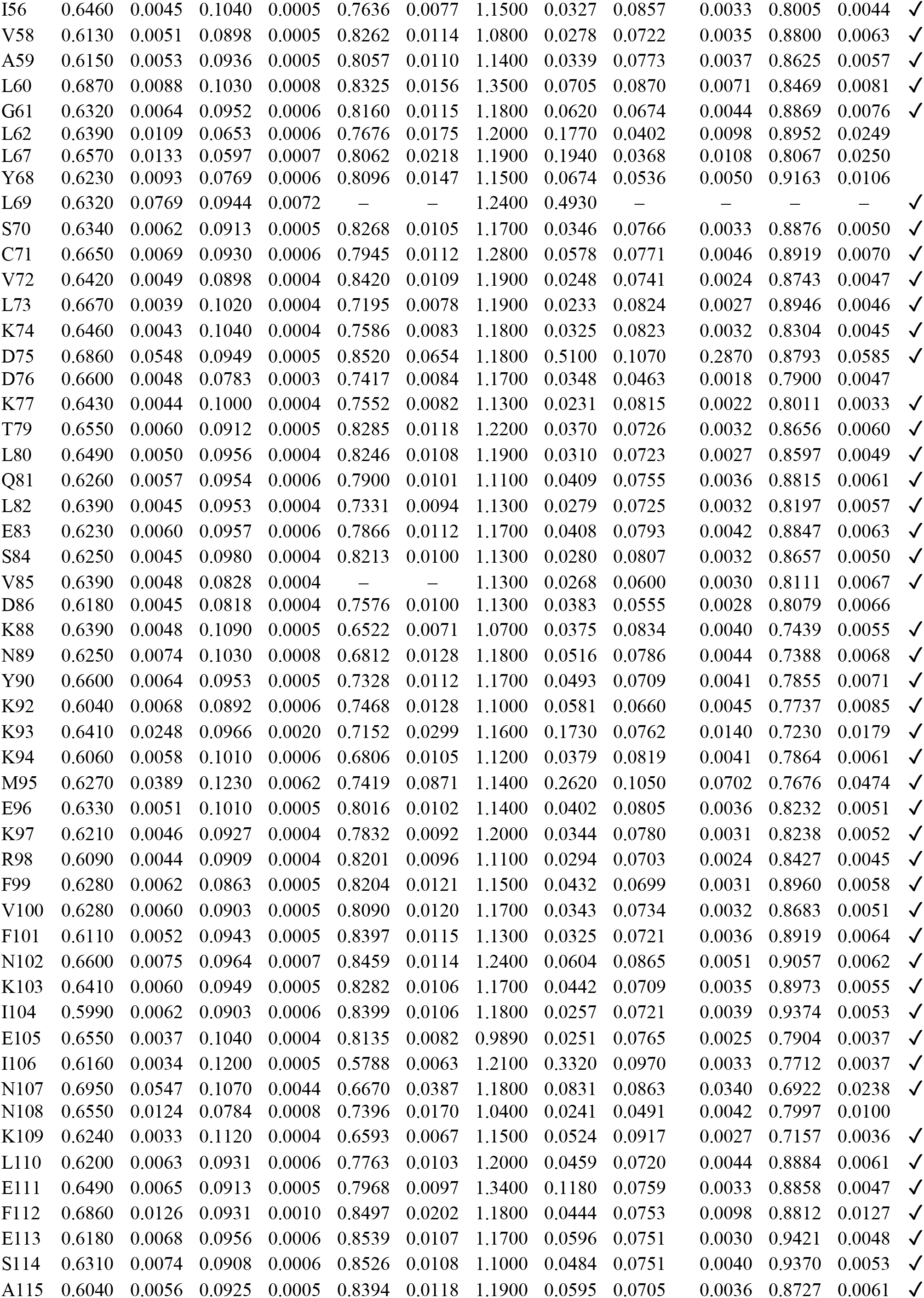
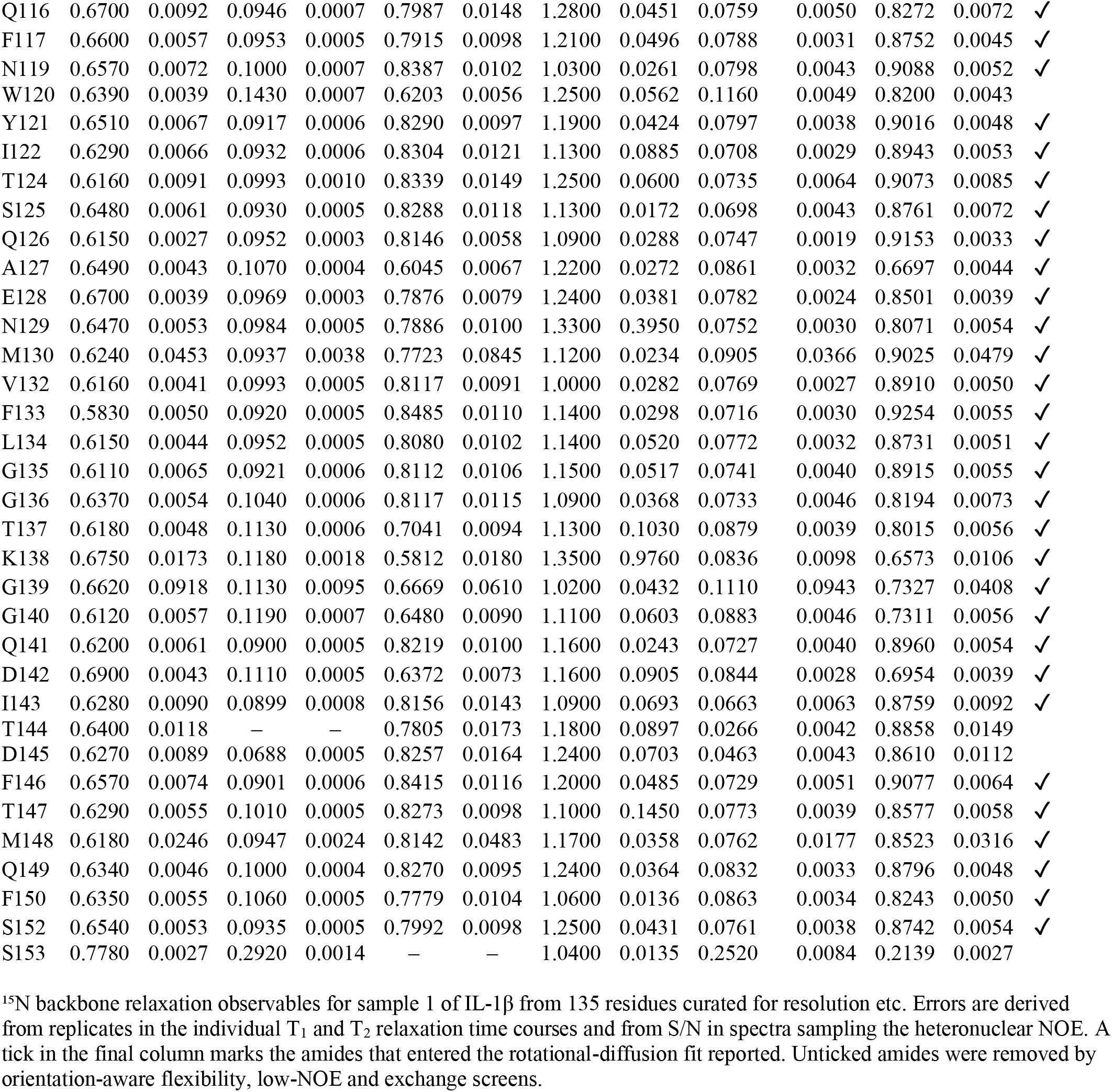
Sample 1 ^15^N-relaxation used to determine the rotational diffusion tensor of IL-1β.

| Res | 500 MHz |  |  |  |  |  | 800 MHz |  |  |  |  |  |
| --- | --- | --- | --- | --- | --- | --- | --- | --- | --- | --- | --- | --- |
|  | T<sub>1</sub>/s | σ/s | T<sub>2</sub>/s | σ/s | NOE | σ | T<sub>1</sub>/s | σ/s | T<sub>2</sub>/s | σ/s | NOE | σ |
| V3 | — | — | 0.1430 | 0.0007 | — | — | — | — | 0.1170 | 0.0047 | — | — |
| R4 | 0.7360 | 0.0797 | 0.1210 | 0.0082 | 0.5091 | 0.0514 | — | — | 0.1700 | 1.5300 | 0.6412 | 0.0341 |
| S5 | 0.5890 | 0.0051 | 0.1030 | 0.0005 | 0.7486 | 0.0109 | 1.0800 | 0.0423 | 0.0814 | 0.0041 | 0.7972 | 0.0063 ✓ |
| L6 | 0.6340 | 0.0054 | 0.1010 | 0.0006 | 0.7753 | 0.0100 | 1.1000 | 0.0380 | 0.0802 | 0.0034 | 0.8741 | 0.0050 ✓ |
| N7 | 0.6380 | 0.0045 | 0.1020 | 0.0004 | 0.8004 | 0.0092 | 1.1800 | 0.0278 | 0.0859 | 0.0030 | 0.8374 | 0.0048 ✓ |
| C8 | 0.6080 | 0.0053 | 0.0913 | 0.0005 | 0.8664 | 0.0104 | 1.0700 | 0.0382 | 0.0715 | 0.0036 | 0.8973 | 0.0059 ✓ |
| T9 | 0.6160 | 0.0052 | 0.1020 | 0.0006 | 0.8056 | 0.0108 | 1.1200 | 0.0353 | 0.0743 | 0.0028 | 0.8932 | 0.0052 ✓ |
| L10 | 0.6400 | 0.0056 | 0.0972 | 0.0005 | 0.8155 | 0.0119 | 1.2200 | 0.0487 | 0.0830 | 0.0049 | 0.8666 | 0.0065 ✓ |
| R11 | 0.6400 | 0.0049 | 0.0957 | 0.0005 | 0.7827 | 0.0090 | 1.2300 | 0.0345 | 0.0811 | 0.0031 | 0.8922 | 0.0045 ✓ |
| D12 | 0.6210 | 0.0060 | 0.0927 | 0.0006 | 0.8314 | 0.0104 | 1.1300 | 0.0507 | 0.0752 | 0.0036 | 0.9344 | 0.0060 ✓ |
| S13 | 0.6340 | 0.0057 | 0.0929 | 0.0005 | 0.8117 | 0.0106 | 1.1800 | 0.0417 | 0.0739 | 0.0032 | 0.8928 | 0.0050 ✓ |
| Q14 | 0.6260 | 0.0065 | 0.0915 | 0.0006 | 0.8225 | 0.0110 | 1.1400 | 0.0478 | 0.0750 | 0.0039 | 0.8821 | 0.0058 ✓ |
| Q15 | 0.6290 | 0.0077 | 0.0873 | 0.0007 | 0.8282 | 0.0119 | 1.1700 | 0.0721 | 0.0607 | 0.0042 | 0.8941 | 0.0065 ✓ |
| K16 | 0.5840 | 0.0559 | — | — | 0.6909 | 0.0492 | 1.0400 | 0.3920 | 0.1110 | 0.0666 | 0.6531 | 0.0352 |
| L18 | 0.6200 | 0.0057 | 0.0988 | 0.0006 | 0.7929 | 0.0124 | 1.1600 | 0.0394 | 0.0813 | 0.0044 | 0.8546 | 0.0066 ✓ |
| V19 | 0.6100 | 0.0048 | 0.0905 | 0.0005 | 0.8412 | 0.0114 | 1.1300 | 0.0216 | 0.0755 | 0.0028 | 0.8735 | 0.0054 ✓ |
| M20 | 0.6180 | 0.0177 | 0.0955 | 0.0018 | — | — | 1.1100 | 0.1020 | 0.0768 | 0.0134 | 0.7905 | 0.0260 ✓ |
| S21 | 0.5860 | 0.0055 | 0.1050 | 0.0006 | 0.6997 | 0.0097 | 1.2100 | 0.0317 | 0.0903 | 0.0038 | 0.7614 | 0.0046 ✓ |
| G22 | 0.5880 | 0.0027 | 0.2140 | 0.0011 | 0.2873 | 0.0036 | 0.8060 | 0.0200 | 0.1660 | 0.0057 | 0.5357 | 0.0032 |
| Y24 | 0.6450 | 0.0040 | 0.1020 | 0.0004 | 0.6850 | 0.0077 | 1.1300 | 0.0277 | 0.0669 | 0.0022 | 0.7716 | 0.0047 ✓ |
| E25 | 0.6260 | 0.0043 | 0.0979 | 0.0004 | 0.7366 | 0.0078 | 1.0900 | 0.0238 | 0.0794 | 0.0027 | 0.8127 | 0.0040 ✓ |
| L26 | 0.6390 | 0.0039 | 0.1000 | 0.0004 | 0.7039 | 0.0083 | 1.1600 | 0.0226 | 0.0787 | 0.0024 | 0.7805 | 0.0046 ✓ |
| K27 | 0.6160 | 0.0046 | 0.0896 | 0.0004 | 0.8143 | 0.0095 | 1.1400 | 0.0255 | 0.0724 | 0.0026 | 0.8788 | 0.0046 ✓ |
| A28 | 0.6410 | 0.0053 | 0.0939 | 0.0005 | 0.8403 | 0.0110 | 1.1800 | 0.0332 | 0.0770 | 0.0030 | 0.8965 | 0.0047 ✓ |
| L29 | 0.6310 | 0.0045 | 0.0936 | 0.0004 | 0.8290 | 0.0094 | 1.1600 | 0.0327 | 0.0727 | 0.0030 | 0.8650 | 0.0050 ✓ |
| H30 | 0.6520 | 0.0071 | 0.0966 | 0.0007 | 0.7831 | 0.0138 | 1.1600 | 0.0805 | 0.0758 | 0.0079 | 0.8832 | 0.0114 ✓ |
| L31 | 0.6460 | 0.0044 | 0.1060 | 0.0004 | 0.8062 | 0.0094 | 1.2000 | 0.0317 | 0.0827 | 0.0030 | 0.8071 | 0.0044 ✓ |
| Q32 | 0.6270 | 0.0039 | 0.1110 | 0.0005 | 0.6678 | 0.0076 | 1.1500 | 0.0250 | 0.0894 | 0.0027 | 0.6857 | 0.0038 ✓ |
| G33 | 0.6300 | 0.0331 | 0.1490 | 0.0063 | 0.6466 | 0.0320 | 0.8960 | 0.1860 | 0.1220 | 0.0303 | 0.7309 | 0.0176 |
| Q34 | 0.5860 | 0.0772 | — | — | 0.7446 | 0.1255 | — | — | — | — | 0.5878 | 0.0723 |
| D35 | 0.5570 | 0.0032 | 0.1160 | 0.0005 | 0.6740 | 0.0074 | 0.9090 | 0.0239 | 0.0935 | 0.0030 | 0.7335 | 0.0038 ✓ |
| M36 | 0.5800 | 0.0312 | 0.0986 | 0.0034 | 0.8473 | 0.0903 | 1.0400 | 0.2400 | 0.0844 | 0.0265 | 0.7673 | 0.0332 ✓ |
| E37 | 0.6000 | 0.0051 | 0.0929 | 0.0005 | 0.7493 | 0.0104 | 1.1100 | 0.0398 | 0.0696 | 0.0034 | 0.7746 | 0.0061 ✓ |
| Q38 | 0.6270 | 0.0041 | 0.0960 | 0.0004 | 0.7604 | 0.0078 | 1.0900 | 0.0246 | 0.0770 | 0.0025 | 0.7896 | 0.0042 ✓ |
| Q39 | 0.6300 | 0.0046 | 0.0924 | 0.0004 | 0.7606 | 0.0099 | 1.0900 | 0.0320 | 0.0709 | 0.0030 | 0.8160 | 0.0056 ✓ |
| V40 | 0.6210 | 0.0042 | 0.0904 | 0.0004 | 0.8008 | 0.0099 | 1.1200 | 0.0284 | 0.0667 | 0.0028 | 0.8768 | 0.0057 ✓ |
| V41 | 0.6310 | 0.0038 | 0.1010 | 0.0004 | 0.7397 | 0.0088 | 1.1400 | 0.0152 | 0.0805 | 0.0024 | 0.8269 | 0.0043 ✓ |
| F42 | 0.6100 | 0.0053 | 0.0911 | 0.0005 | 0.8446 | 0.0108 | 1.1000 | 0.0427 | 0.0657 | 0.0037 | 0.8858 | 0.0068 ✓ |
| S43 | 0.6450 | 0.0062 | 0.0975 | 0.0006 | 0.8113 | 0.0104 | 1.2300 | 0.0504 | 0.0807 | 0.0043 | 0.8983 | 0.0062 ✓ |
| M44 | 0.6270 | 0.0445 | 0.0917 | 0.0041 | 0.8325 | 0.0809 | — | — | 0.1190 | 0.3130 | 0.8026 | 0.0484 ✓ |
| S45 | 0.6310 | 0.0050 | 0.0926 | 0.0005 | 0.8332 | 0.0110 | 1.1500 | 0.0336 | 0.0780 | 0.0031 | 0.8693 | 0.0053 ✓ |
| F46 | 0.6280 | 0.0053 | 0.0966 | 0.0005 | 0.8017 | 0.0092 | 1.1800 | 0.0392 | 0.0802 | 0.0036 | 0.8675 | 0.0059 ✓ |
| V47 | 0.6570 | 0.0084 | 0.0943 | 0.0007 | 0.7304 | 0.0149 | 1.2000 | 0.0419 | 0.0720 | 0.0052 | 0.7867 | 0.0101 ✓ |
| Q48 | 0.6010 | 0.0047 | 0.0962 | 0.0005 | 0.7199 | 0.0097 | 1.0700 | 0.0399 | 0.0763 | 0.0036 | 0.8298 | 0.0064 ✓ |
| G49 | 0.6690 | 0.0090 | 0.1280 | 0.0010 | 0.5493 | 0.0107 | 1.1700 | 0.0600 | 0.0955 | 0.0069 | 0.6013 | 0.0076 |
| E50 | 0.6310 | 0.0050 | 0.0978 | 0.0005 | 0.7539 | 0.0091 | 1.1600 | 0.0344 | 0.0775 | 0.0035 | 0.8688 | 0.0054 ✓ |
| E51 | 0.6630 | 0.0076 | 0.1560 | 0.0014 | 0.3680 | 0.0076 | 1.0300 | 0.0494 | 0.1220 | 0.0081 | 0.5295 | 0.0063 |
| D54 | 0.6430 | 0.0136 | 0.1220 | 0.0016 | 0.5191 | 0.0142 | 1.0700 | 0.0875 | 0.0980 | 0.0102 | 0.6273 | 0.0094 |
| K55 | 0.6600 | 0.0040 | 0.1240 | 0.0006 | 0.6191 | 0.0077 | 1.0700 | 0.0231 | 0.1020 | 0.0034 | 0.6618 | 0.0036 |

|  |  |  |  |  |  |  |  |  |  |  |  |  |  |
| --- | --- | --- | --- | --- | --- | --- | --- | --- | --- | --- | --- | --- | --- |
| I56 | 0.6460 | 0.0045 | 0.1040 | 0.0005 | 0.7636 | 0.0077 | 1.1500 | 0.0327 | 0.0857 | 0.0033 | 0.8005 | 0.0044 | ✓ |
| V58 | 0.6130 | 0.0051 | 0.0898 | 0.0005 | 0.8262 | 0.0114 | 1.0800 | 0.0278 | 0.0722 | 0.0035 | 0.8800 | 0.0063 | ✓ |
| A59 | 0.6150 | 0.0053 | 0.0936 | 0.0005 | 0.8057 | 0.0110 | 1.1400 | 0.0339 | 0.0773 | 0.0037 | 0.8625 | 0.0057 | ✓ |
| L60 | 0.6870 | 0.0088 | 0.1030 | 0.0008 | 0.8325 | 0.0156 | 1.3500 | 0.0705 | 0.0870 | 0.0071 | 0.8469 | 0.0081 | ✓ |
| G61 | 0.6320 | 0.0064 | 0.0952 | 0.0006 | 0.8160 | 0.0115 | 1.1800 | 0.0620 | 0.0674 | 0.0044 | 0.8869 | 0.0076 | ✓ |
| L62 | 0.6390 | 0.0109 | 0.0653 | 0.0006 | 0.7676 | 0.0175 | 1.2000 | 0.1770 | 0.0402 | 0.0098 | 0.8952 | 0.0249 |  |
| L67 | 0.6570 | 0.0133 | 0.0597 | 0.0007 | 0.8062 | 0.0218 | 1.1900 | 0.1940 | 0.0368 | 0.0108 | 0.8067 | 0.0250 |  |
| Y68 | 0.6230 | 0.0093 | 0.0769 | 0.0006 | 0.8096 | 0.0147 | 1.1500 | 0.0674 | 0.0536 | 0.0050 | 0.9163 | 0.0106 |  |
| L69 | 0.6320 | 0.0769 | 0.0944 | 0.0072 | — | — | 1.2400 | 0.4930 | — | — | — | — | ✓ |
| S70 | 0.6340 | 0.0062 | 0.0913 | 0.0005 | 0.8268 | 0.0105 | 1.1700 | 0.0346 | 0.0766 | 0.0033 | 0.8876 | 0.0050 | ✓ |
| C71 | 0.6650 | 0.0069 | 0.0930 | 0.0006 | 0.7945 | 0.0112 | 1.2800 | 0.0578 | 0.0771 | 0.0046 | 0.8919 | 0.0070 | ✓ |
| V72 | 0.6420 | 0.0049 | 0.0898 | 0.0004 | 0.8420 | 0.0109 | 1.1900 | 0.0248 | 0.0741 | 0.0024 | 0.8743 | 0.0047 | ✓ |
| L73 | 0.6670 | 0.0039 | 0.1020 | 0.0004 | 0.7195 | 0.0078 | 1.1900 | 0.0233 | 0.0824 | 0.0027 | 0.8946 | 0.0046 | ✓ |
| K74 | 0.6460 | 0.0043 | 0.1040 | 0.0004 | 0.7586 | 0.0083 | 1.1800 | 0.0325 | 0.0823 | 0.0032 | 0.8304 | 0.0045 | ✓ |
| D75 | 0.6860 | 0.0548 | 0.0949 | 0.0005 | 0.8520 | 0.0654 | 1.1800 | 0.5100 | 0.1070 | 0.2870 | 0.8793 | 0.0585 | ✓ |
| D76 | 0.6600 | 0.0048 | 0.0783 | 0.0003 | 0.7417 | 0.0084 | 1.1700 | 0.0348 | 0.0463 | 0.0018 | 0.7900 | 0.0047 |  |
| K77 | 0.6430 | 0.0044 | 0.1000 | 0.0004 | 0.7552 | 0.0082 | 1.1300 | 0.0231 | 0.0815 | 0.0022 | 0.8011 | 0.0033 | ✓ |
| T79 | 0.6550 | 0.0060 | 0.0912 | 0.0005 | 0.8285 | 0.0118 | 1.2200 | 0.0370 | 0.0726 | 0.0032 | 0.8656 | 0.0060 | ✓ |
| L80 | 0.6490 | 0.0050 | 0.0956 | 0.0004 | 0.8246 | 0.0108 | 1.1900 | 0.0310 | 0.0723 | 0.0027 | 0.8597 | 0.0049 | ✓ |
| Q81 | 0.6260 | 0.0057 | 0.0954 | 0.0006 | 0.7900 | 0.0101 | 1.1100 | 0.0409 | 0.0755 | 0.0036 | 0.8815 | 0.0061 | ✓ |
| L82 | 0.6390 | 0.0045 | 0.0953 | 0.0004 | 0.7331 | 0.0094 | 1.1300 | 0.0279 | 0.0725 | 0.0032 | 0.8197 | 0.0057 | ✓ |
| E83 | 0.6230 | 0.0060 | 0.0957 | 0.0006 | 0.7866 | 0.0112 | 1.1700 | 0.0408 | 0.0793 | 0.0042 | 0.8847 | 0.0063 | ✓ |
| S84 | 0.6250 | 0.0045 | 0.0980 | 0.0004 | 0.8213 | 0.0100 | 1.1300 | 0.0280 | 0.0807 | 0.0032 | 0.8657 | 0.0050 | ✓ |
| V85 | 0.6390 | 0.0048 | 0.0828 | 0.0004 | — | — | 1.1300 | 0.0268 | 0.0600 | 0.0030 | 0.8111 | 0.0067 | ✓ |
| D86 | 0.6180 | 0.0045 | 0.0818 | 0.0004 | 0.7576 | 0.0100 | 1.1300 | 0.0383 | 0.0555 | 0.0028 | 0.8079 | 0.0066 |  |
| K88 | 0.6390 | 0.0048 | 0.1090 | 0.0005 | 0.6522 | 0.0071 | 1.0700 | 0.0375 | 0.0834 | 0.0040 | 0.7439 | 0.0055 | ✓ |
| N89 | 0.6250 | 0.0074 | 0.1030 | 0.0008 | 0.6812 | 0.0128 | 1.1800 | 0.0516 | 0.0786 | 0.0044 | 0.7388 | 0.0068 | ✓ |
| Y90 | 0.6600 | 0.0064 | 0.0953 | 0.0005 | 0.7328 | 0.0112 | 1.1700 | 0.0493 | 0.0709 | 0.0041 | 0.7855 | 0.0071 | ✓ |
| K92 | 0.6040 | 0.0068 | 0.0892 | 0.0006 | 0.7468 | 0.0128 | 1.1000 | 0.0581 | 0.0660 | 0.0045 | 0.7737 | 0.0085 | ✓ |
| K93 | 0.6410 | 0.0248 | 0.0966 | 0.0020 | 0.7152 | 0.0299 | 1.1600 | 0.1730 | 0.0762 | 0.0140 | 0.7230 | 0.0179 | ✓ |
| K94 | 0.6060 | 0.0058 | 0.1010 | 0.0006 | 0.6806 | 0.0105 | 1.1200 | 0.0379 | 0.0819 | 0.0041 | 0.7864 | 0.0061 | ✓ |
| M95 | 0.6270 | 0.0389 | 0.1230 | 0.0062 | 0.7419 | 0.0871 | 1.1400 | 0.2620 | 0.1050 | 0.0702 | 0.7676 | 0.0474 | ✓ |
| E96 | 0.6330 | 0.0051 | 0.1010 | 0.0005 | 0.8016 | 0.0102 | 1.1400 | 0.0402 | 0.0805 | 0.0036 | 0.8232 | 0.0051 | ✓ |
| K97 | 0.6210 | 0.0046 | 0.0927 | 0.0004 | 0.7832 | 0.0092 | 1.2000 | 0.0344 | 0.0780 | 0.0031 | 0.8238 | 0.0052 | ✓ |
| R98 | 0.6090 | 0.0044 | 0.0909 | 0.0004 | 0.8201 | 0.0096 | 1.1100 | 0.0294 | 0.0703 | 0.0024 | 0.8427 | 0.0045 | ✓ |
| F99 | 0.6280 | 0.0062 | 0.0863 | 0.0005 | 0.8204 | 0.0121 | 1.1500 | 0.0432 | 0.0699 | 0.0031 | 0.8960 | 0.0058 | ✓ |
| V100 | 0.6280 | 0.0060 | 0.0903 | 0.0005 | 0.8090 | 0.0120 | 1.1700 | 0.0343 | 0.0734 | 0.0032 | 0.8683 | 0.0051 | ✓ |
| F101 | 0.6110 | 0.0052 | 0.0943 | 0.0005 | 0.8397 | 0.0115 | 1.1300 | 0.0325 | 0.0721 | 0.0036 | 0.8919 | 0.0064 | ✓ |
| N102 | 0.6600 | 0.0075 | 0.0964 | 0.0007 | 0.8459 | 0.0114 | 1.2400 | 0.0604 | 0.0865 | 0.0051 | 0.9057 | 0.0062 | ✓ |
| K103 | 0.6410 | 0.0060 | 0.0949 | 0.0005 | 0.8282 | 0.0106 | 1.1700 | 0.0442 | 0.0709 | 0.0035 | 0.8973 | 0.0055 | ✓ |
| I104 | 0.5990 | 0.0062 | 0.0903 | 0.0006 | 0.8399 | 0.0106 | 1.1800 | 0.0257 | 0.0721 | 0.0039 | 0.9374 | 0.0053 | ✓ |
| E105 | 0.6550 | 0.0037 | 0.1040 | 0.0004 | 0.8135 | 0.0082 | 0.9890 | 0.0251 | 0.0765 | 0.0025 | 0.7904 | 0.0037 | ✓ |
| I106 | 0.6160 | 0.0034 | 0.1200 | 0.0005 | 0.5788 | 0.0063 | 1.2100 | 0.3320 | 0.0970 | 0.0033 | 0.7712 | 0.0037 | ✓ |
| N107 | 0.6950 | 0.0547 | 0.1070 | 0.0044 | 0.6670 | 0.0387 | 1.1800 | 0.0831 | 0.0863 | 0.0340 | 0.6922 | 0.0238 | ✓ |
| N108 | 0.6550 | 0.0124 | 0.0784 | 0.0008 | 0.7396 | 0.0170 | 1.0400 | 0.0241 | 0.0491 | 0.0042 | 0.7997 | 0.0100 |  |
| K109 | 0.6240 | 0.0033 | 0.1120 | 0.0004 | 0.6593 | 0.0067 | 1.1500 | 0.0524 | 0.0917 | 0.0027 | 0.7157 | 0.0036 | ✓ |
| L110 | 0.6200 | 0.0063 | 0.0931 | 0.0006 | 0.7763 | 0.0103 | 1.2000 | 0.0459 | 0.0720 | 0.0044 | 0.8884 | 0.0061 | ✓ |
| E111 | 0.6490 | 0.0065 | 0.0913 | 0.0005 | 0.7968 | 0.0097 | 1.3400 | 0.1180 | 0.0759 | 0.0033 | 0.8858 | 0.0047 | ✓ |
| F112 | 0.6860 | 0.0126 | 0.0931 | 0.0010 | 0.8497 | 0.0202 | 1.1800 | 0.0444 | 0.0753 | 0.0098 | 0.8812 | 0.0127 | ✓ |
| E113 | 0.6180 | 0.0068 | 0.0956 | 0.0006 | 0.8539 | 0.0107 | 1.1700 | 0.0596 | 0.0751 | 0.0030 | 0.9421 | 0.0048 | ✓ |
| S114 | 0.6310 | 0.0074 | 0.0908 | 0.0006 | 0.8526 | 0.0108 | 1.1000 | 0.0484 | 0.0751 | 0.0040 | 0.9370 | 0.0053 | ✓ |
| A115 | 0.6040 | 0.0056 | 0.0925 | 0.0005 | 0.8394 | 0.0118 | 1.1900 | 0.0595 | 0.0705 | 0.0036 | 0.8727 | 0.0061 | ✓ |
| Q116 | 0.6700 | 0.0092 | 0.0946 | 0.0007 | 0.7987 | 0.0148 | 1.2800 | 0.0451 | 0.0759 | 0.0050 | 0.8272 | 0.0072 | ✓ |
| F117 | 0.6600 | 0.0057 | 0.0953 | 0.0005 | 0.7915 | 0.0098 | 1.2100 | 0.0496 | 0.0788 | 0.0031 | 0.8752 | 0.0045 | ✓ |
| N119 | 0.6570 | 0.0072 | 0.1000 | 0.0007 | 0.8387 | 0.0102 | 1.0300 | 0.0261 | 0.0798 | 0.0043 | 0.9088 | 0.0052 | ✓ |
| W120 | 0.6390 | 0.0039 | 0.1430 | 0.0007 | 0.6203 | 0.0056 | 1.2500 | 0.0562 | 0.1160 | 0.0049 | 0.8200 | 0.0043 |  |
| Y121 | 0.6510 | 0.0067 | 0.0917 | 0.0006 | 0.8290 | 0.0097 | 1.1900 | 0.0424 | 0.0797 | 0.0038 | 0.9016 | 0.0048 | ✓ |
| I122 | 0.6290 | 0.0066 | 0.0932 | 0.0006 | 0.8304 | 0.0121 | 1.1300 | 0.0885 | 0.0708 | 0.0029 | 0.8943 | 0.0053 | ✓ |
| T124 | 0.6160 | 0.0091 | 0.0993 | 0.0010 | 0.8339 | 0.0149 | 1.2500 | 0.0600 | 0.0735 | 0.0064 | 0.9073 | 0.0085 | ✓ |
| S125 | 0.6480 | 0.0061 | 0.0930 | 0.0005 | 0.8288 | 0.0118 | 1.1300 | 0.0172 | 0.0698 | 0.0043 | 0.8761 | 0.0072 | ✓ |
| Q126 | 0.6150 | 0.0027 | 0.0952 | 0.0003 | 0.8146 | 0.0058 | 1.0900 | 0.0288 | 0.0747 | 0.0019 | 0.9153 | 0.0033 | ✓ |
| A127 | 0.6490 | 0.0043 | 0.1070 | 0.0004 | 0.6045 | 0.0067 | 1.2200 | 0.0272 | 0.0861 | 0.0032 | 0.6697 | 0.0044 | ✓ |
| E128 | 0.6700 | 0.0039 | 0.0969 | 0.0003 | 0.7876 | 0.0079 | 1.2400 | 0.0381 | 0.0782 | 0.0024 | 0.8501 | 0.0039 | ✓ |
| N129 | 0.6470 | 0.0053 | 0.0984 | 0.0005 | 0.7886 | 0.0100 | 1.3300 | 0.3950 | 0.0752 | 0.0030 | 0.8071 | 0.0054 | ✓ |
| M130 | 0.6240 | 0.0453 | 0.0937 | 0.0038 | 0.7723 | 0.0845 | 1.1200 | 0.0234 | 0.0905 | 0.0366 | 0.9025 | 0.0479 | ✓ |
| V132 | 0.6160 | 0.0041 | 0.0993 | 0.0005 | 0.8117 | 0.0091 | 1.0000 | 0.0282 | 0.0769 | 0.0027 | 0.8910 | 0.0050 | ✓ |
| F133 | 0.5830 | 0.0050 | 0.0920 | 0.0005 | 0.8485 | 0.0110 | 1.1400 | 0.0298 | 0.0716 | 0.0030 | 0.9254 | 0.0055 | ✓ |
| L134 | 0.6150 | 0.0044 | 0.0952 | 0.0005 | 0.8080 | 0.0102 | 1.1400 | 0.0520 | 0.0772 | 0.0032 | 0.8731 | 0.0051 | ✓ |
| G135 | 0.6110 | 0.0065 | 0.0921 | 0.0006 | 0.8112 | 0.0106 | 1.1500 | 0.0517 | 0.0741 | 0.0040 | 0.8915 | 0.0055 | ✓ |
| G136 | 0.6370 | 0.0054 | 0.1040 | 0.0006 | 0.8117 | 0.0115 | 1.0900 | 0.0368 | 0.0733 | 0.0046 | 0.8194 | 0.0073 | ✓ |
| T137 | 0.6180 | 0.0048 | 0.1130 | 0.0006 | 0.7041 | 0.0094 | 1.1300 | 0.1030 | 0.0879 | 0.0039 | 0.8015 | 0.0056 | ✓ |
| K138 | 0.6750 | 0.0173 | 0.1180 | 0.0018 | 0.5812 | 0.0180 | 1.3500 | 0.9760 | 0.0836 | 0.0098 | 0.6573 | 0.0106 | ✓ |
| G139 | 0.6620 | 0.0918 | 0.1130 | 0.0095 | 0.6669 | 0.0610 | 1.0200 | 0.0432 | 0.1110 | 0.0943 | 0.7327 | 0.0408 | ✓ |
| G140 | 0.6120 | 0.0057 | 0.1190 | 0.0007 | 0.6480 | 0.0090 | 1.1100 | 0.0603 | 0.0883 | 0.0046 | 0.7311 | 0.0056 | ✓ |
| Q141 | 0.6200 | 0.0061 | 0.0900 | 0.0005 | 0.8219 | 0.0100 | 1.1600 | 0.0243 | 0.0727 | 0.0040 | 0.8960 | 0.0054 | ✓ |
| D142 | 0.6900 | 0.0043 | 0.1110 | 0.0005 | 0.6372 | 0.0073 | 1.1600 | 0.0905 | 0.0844 | 0.0028 | 0.6954 | 0.0039 | ✓ |
| I143 | 0.6280 | 0.0090 | 0.0899 | 0.0008 | 0.8156 | 0.0143 | 1.0900 | 0.0693 | 0.0663 | 0.0063 | 0.8759 | 0.0092 | ✓ |
| T144 | 0.6400 | 0.0118 | — | — | 0.7805 | 0.0173 | 1.1800 | 0.0897 | 0.0266 | 0.0042 | 0.8858 | 0.0149 |  |
| D145 | 0.6270 | 0.0089 | 0.0688 | 0.0005 | 0.8257 | 0.0164 | 1.2400 | 0.0703 | 0.0463 | 0.0043 | 0.8610 | 0.0112 |  |
| F146 | 0.6570 | 0.0074 | 0.0901 | 0.0006 | 0.8415 | 0.0116 | 1.2000 | 0.0485 | 0.0729 | 0.0051 | 0.9077 | 0.0064 | ✓ |
| T147 | 0.6290 | 0.0055 | 0.1010 | 0.0005 | 0.8273 | 0.0098 | 1.1000 | 0.1450 | 0.0773 | 0.0039 | 0.8577 | 0.0058 | ✓ |
| M148 | 0.6180 | 0.0246 | 0.0947 | 0.0024 | 0.8142 | 0.0483 | 1.1700 | 0.0358 | 0.0762 | 0.0177 | 0.8523 | 0.0316 | ✓ |
| Q149 | 0.6340 | 0.0046 | 0.1000 | 0.0004 | 0.8270 | 0.0095 | 1.2400 | 0.0364 | 0.0832 | 0.0033 | 0.8796 | 0.0048 | ✓ |
| F150 | 0.6350 | 0.0055 | 0.1060 | 0.0005 | 0.7779 | 0.0104 | 1.0600 | 0.0136 | 0.0863 | 0.0034 | 0.8243 | 0.0050 | ✓ |
| S152 | 0.6540 | 0.0053 | 0.0935 | 0.0005 | 0.7992 | 0.0098 | 1.2500 | 0.0431 | 0.0761 | 0.0038 | 0.8742 | 0.0054 | ✓ |
| S153 | 0.7780 | 0.0027 | 0.2920 | 0.0014 | — | — | 1.0400 | 0.0135 | 0.2520 | 0.0084 | 0.2139 | 0.0027 |  |

<sup>15</sup>N backbone relaxation observables for sample 1 of IL-1 $\beta$ from 135 residues curated for resolution etc. Errors are derived from replicates in the individual T<sub>1</sub> and T<sub>2</sub> relaxation time courses and from S/N in spectra sampling the heteronuclear NOE. A tick in the final column marks the amides that entered the rotational-diffusion fit reported. Unticked amides were removed by orientation-aware flexibility, low-NOE and exchange screens.

**Table S2:**
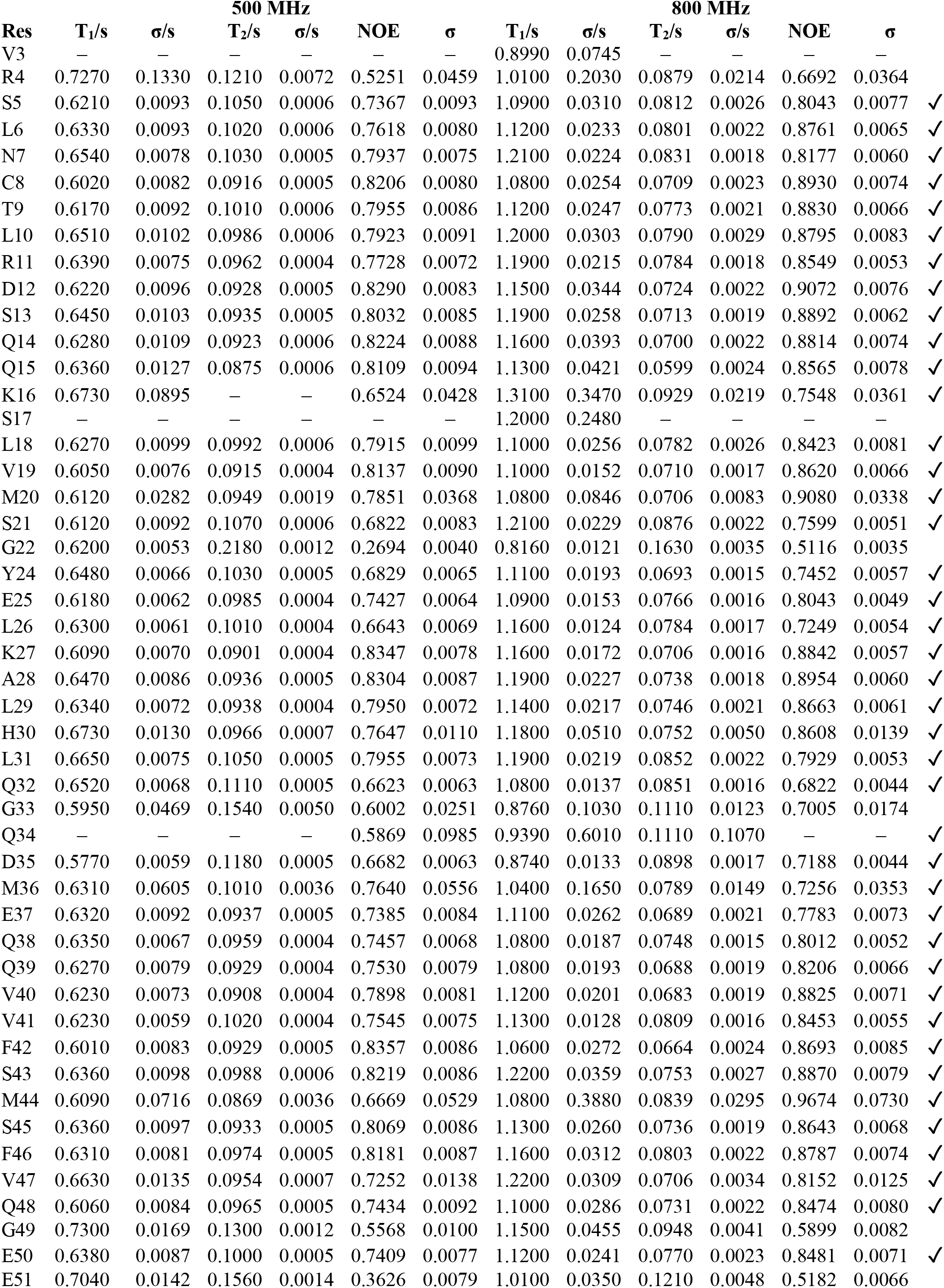
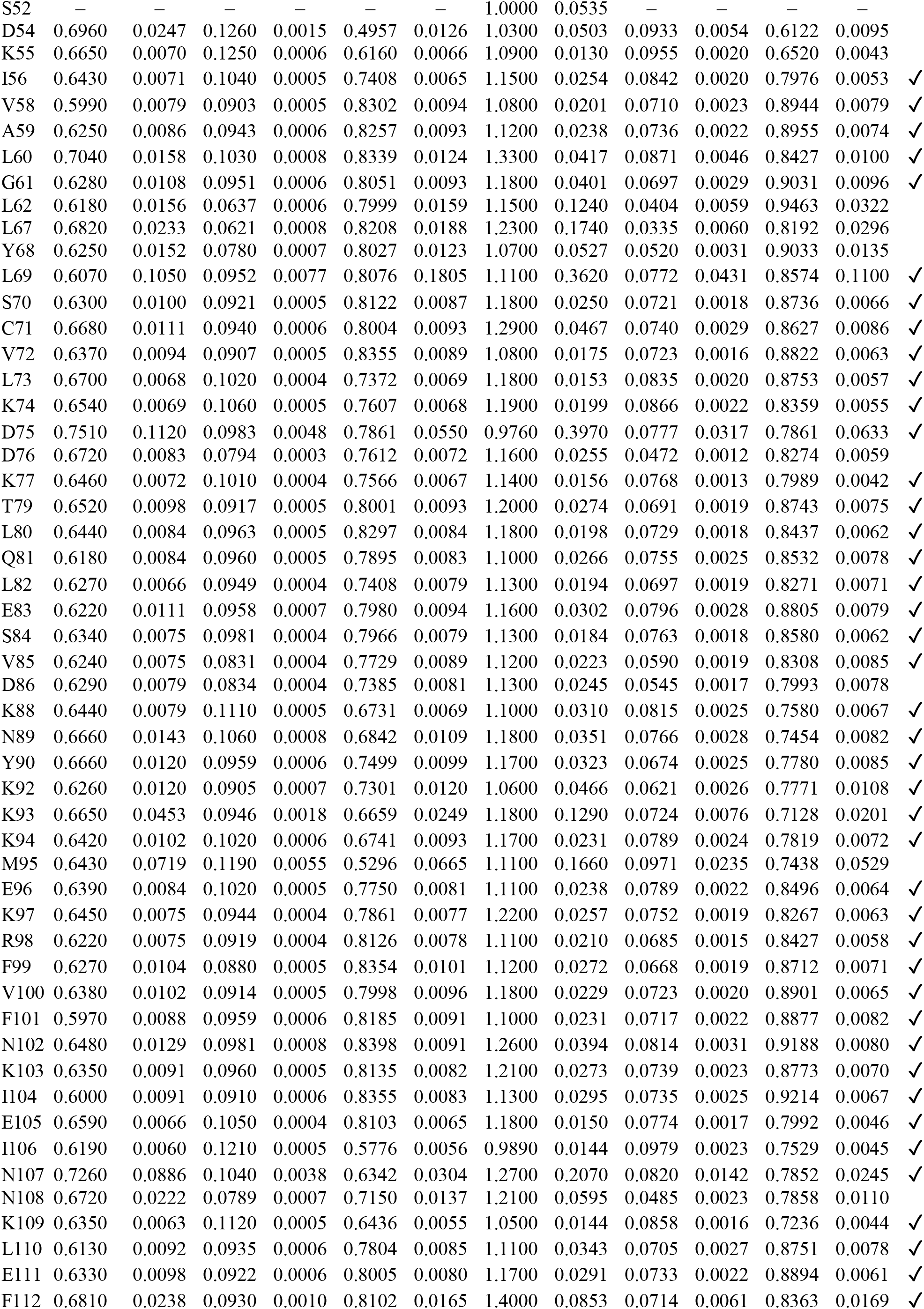
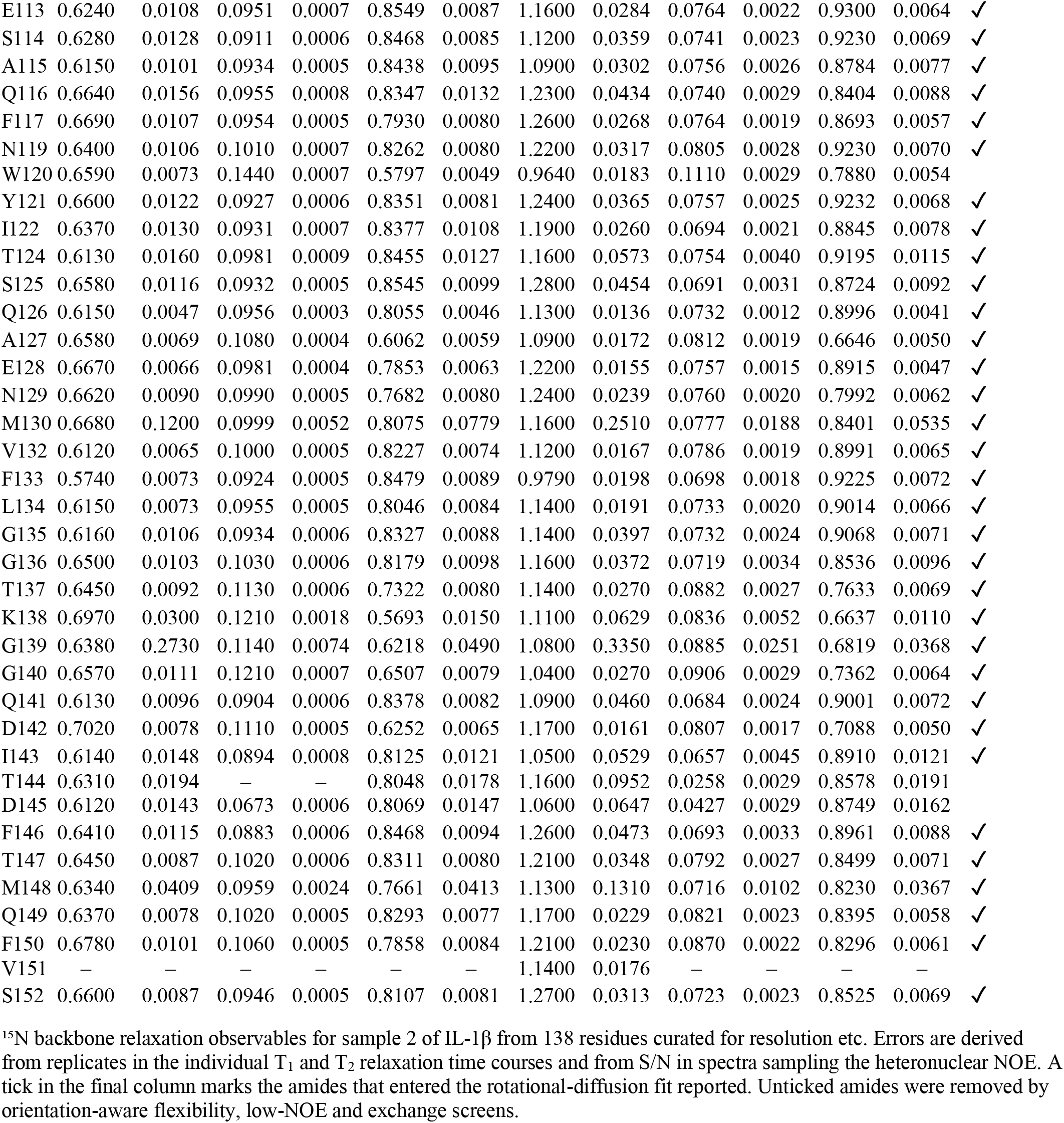
Sample 2 ^15^N-relaxation used to determine the rotational diffusion tensor of IL-1β.

| Res | 500 MHz |  |  |  |  |  | 800 MHz |  |  |  |  |  |
| --- | --- | --- | --- | --- | --- | --- | --- | --- | --- | --- | --- | --- |
| | T<sub>1</sub>/s | $\sigma$/s | T<sub>2</sub>/s | $\sigma$/s | NOE | $\sigma$ | T<sub>1</sub>/s | $\sigma$/s | T<sub>2</sub>/s | $\sigma$/s | NOE | $\sigma$ |
| V3 | — | — | — | — | — | — | 0.8990 | 0.0745 | — | — | — | — |
| R4 | 0.7270 | 0.1330 | 0.1210 | 0.0072 | 0.5251 | 0.0459 | 1.0100 | 0.2030 | 0.0879 | 0.0214 | 0.6692 | 0.0364 |
| S5 | 0.6210 | 0.0093 | 0.1050 | 0.0006 | 0.7367 | 0.0093 | 1.0900 | 0.0310 | 0.0812 | 0.0026 | 0.8043 | 0.0077 ✓ |
| L6 | 0.6330 | 0.0093 | 0.1020 | 0.0006 | 0.7618 | 0.0080 | 1.1200 | 0.0233 | 0.0801 | 0.0022 | 0.8761 | 0.0065 ✓ |
| N7 | 0.6540 | 0.0078 | 0.1030 | 0.0005 | 0.7937 | 0.0075 | 1.2100 | 0.0224 | 0.0831 | 0.0018 | 0.8177 | 0.0060 ✓ |
| C8 | 0.6020 | 0.0082 | 0.0916 | 0.0005 | 0.8206 | 0.0080 | 1.0800 | 0.0254 | 0.0709 | 0.0023 | 0.8930 | 0.0074 ✓ |
| T9 | 0.6170 | 0.0092 | 0.1010 | 0.0006 | 0.7955 | 0.0086 | 1.1200 | 0.0247 | 0.0773 | 0.0021 | 0.8830 | 0.0066 ✓ |
| L10 | 0.6510 | 0.0102 | 0.0986 | 0.0006 | 0.7923 | 0.0091 | 1.2000 | 0.0303 | 0.0790 | 0.0029 | 0.8795 | 0.0083 ✓ |
| R11 | 0.6390 | 0.0075 | 0.0962 | 0.0004 | 0.7728 | 0.0072 | 1.1900 | 0.0215 | 0.0784 | 0.0018 | 0.8549 | 0.0053 ✓ |
| D12 | 0.6220 | 0.0096 | 0.0928 | 0.0005 | 0.8290 | 0.0083 | 1.1500 | 0.0344 | 0.0724 | 0.0022 | 0.9072 | 0.0076 ✓ |
| S13 | 0.6450 | 0.0103 | 0.0935 | 0.0005 | 0.8032 | 0.0085 | 1.1900 | 0.0258 | 0.0713 | 0.0019 | 0.8892 | 0.0062 ✓ |
| Q14 | 0.6280 | 0.0109 | 0.0923 | 0.0006 | 0.8224 | 0.0088 | 1.1600 | 0.0393 | 0.0700 | 0.0022 | 0.8814 | 0.0074 ✓ |
| Q15 | 0.6360 | 0.0127 | 0.0875 | 0.0006 | 0.8109 | 0.0094 | 1.1300 | 0.0421 | 0.0599 | 0.0024 | 0.8565 | 0.0078 ✓ |
| K16 | 0.6730 | 0.0895 | — | — | 0.6524 | 0.0428 | 1.3100 | 0.3470 | 0.0929 | 0.0219 | 0.7548 | 0.0361 ✓ |
| S17 | — | — | — | — | — | — | 1.2000 | 0.2480 | — | — | — | — |
| L18 | 0.6270 | 0.0099 | 0.0992 | 0.0006 | 0.7915 | 0.0099 | 1.1000 | 0.0256 | 0.0782 | 0.0026 | 0.8423 | 0.0081 ✓ |
| V19 | 0.6050 | 0.0076 | 0.0915 | 0.0004 | 0.8137 | 0.0090 | 1.1000 | 0.0152 | 0.0710 | 0.0017 | 0.8620 | 0.0066 ✓ |
| M20 | 0.6120 | 0.0282 | 0.0949 | 0.0019 | 0.7851 | 0.0368 | 1.0800 | 0.0846 | 0.0706 | 0.0083 | 0.9080 | 0.0338 ✓ |
| S21 | 0.6120 | 0.0092 | 0.1070 | 0.0006 | 0.6822 | 0.0083 | 1.2100 | 0.0229 | 0.0876 | 0.0022 | 0.7599 | 0.0051 ✓ |
| G22 | 0.6200 | 0.0053 | 0.2180 | 0.0012 | 0.2694 | 0.0040 | 0.8160 | 0.0121 | 0.1630 | 0.0035 | 0.5116 | 0.0035 |
| Y24 | 0.6480 | 0.0066 | 0.1030 | 0.0005 | 0.6829 | 0.0065 | 1.1100 | 0.0193 | 0.0693 | 0.0015 | 0.7452 | 0.0057 ✓ |
| E25 | 0.6180 | 0.0062 | 0.0985 | 0.0004 | 0.7427 | 0.0064 | 1.0900 | 0.0153 | 0.0766 | 0.0016 | 0.8043 | 0.0049 ✓ |
| L26 | 0.6300 | 0.0061 | 0.1010 | 0.0004 | 0.6643 | 0.0069 | 1.1600 | 0.0124 | 0.0784 | 0.0017 | 0.7249 | 0.0054 ✓ |
| K27 | 0.6090 | 0.0070 | 0.0901 | 0.0004 | 0.8347 | 0.0078 | 1.1600 | 0.0172 | 0.0706 | 0.0016 | 0.8842 | 0.0057 ✓ |
| A28 | 0.6470 | 0.0086 | 0.0936 | 0.0005 | 0.8304 | 0.0087 | 1.1900 | 0.0227 | 0.0738 | 0.0018 | 0.8954 | 0.0060 ✓ |
| L29 | 0.6340 | 0.0072 | 0.0938 | 0.0004 | 0.7950 | 0.0072 | 1.1400 | 0.0217 | 0.0746 | 0.0021 | 0.8663 | 0.0061 ✓ |
| H30 | 0.6730 | 0.0130 | 0.0966 | 0.0007 | 0.7647 | 0.0110 | 1.1800 | 0.0510 | 0.0752 | 0.0050 | 0.8608 | 0.0139 ✓ |
| L31 | 0.6650 | 0.0075 | 0.1050 | 0.0005 | 0.7955 | 0.0073 | 1.1900 | 0.0219 | 0.0852 | 0.0022 | 0.7929 | 0.0053 ✓ |
| Q32 | 0.6520 | 0.0068 | 0.1110 | 0.0005 | 0.6623 | 0.0063 | 1.0800 | 0.0137 | 0.0851 | 0.0016 | 0.6822 | 0.0044 ✓ |
| G33 | 0.5950 | 0.0469 | 0.1540 | 0.0050 | 0.6002 | 0.0251 | 0.8760 | 0.1030 | 0.1110 | 0.0123 | 0.7005 | 0.0174 |
| Q34 | — | — | — | — | 0.5869 | 0.0985 | 0.9390 | 0.6010 | 0.1110 | 0.1070 | — | — ✓ |
| D35 | 0.5770 | 0.0059 | 0.1180 | 0.0005 | 0.6682 | 0.0063 | 0.8740 | 0.0133 | 0.0898 | 0.0017 | 0.7188 | 0.0044 ✓ |
| M36 | 0.6310 | 0.0605 | 0.1010 | 0.0036 | 0.7640 | 0.0556 | 1.0400 | 0.1650 | 0.0789 | 0.0149 | 0.7256 | 0.0353 ✓ |
| E37 | 0.6320 | 0.0092 | 0.0937 | 0.0005 | 0.7385 | 0.0084 | 1.1100 | 0.0262 | 0.0689 | 0.0021 | 0.7783 | 0.0073 ✓ |
| Q38 | 0.6350 | 0.0067 | 0.0959 | 0.0004 | 0.7457 | 0.0068 | 1.0800 | 0.0187 | 0.0748 | 0.0015 | 0.8012 | 0.0052 ✓ |
| Q39 | 0.6270 | 0.0079 | 0.0929 | 0.0004 | 0.7530 | 0.0079 | 1.0800 | 0.0193 | 0.0688 | 0.0019 | 0.8206 | 0.0066 ✓ |
| V40 | 0.6230 | 0.0073 | 0.0908 | 0.0004 | 0.7898 | 0.0081 | 1.1200 | 0.0201 | 0.0683 | 0.0019 | 0.8825 | 0.0071 ✓ |
| V41 | 0.6230 | 0.0059 | 0.1020 | 0.0004 | 0.7545 | 0.0075 | 1.1300 | 0.0128 | 0.0809 | 0.0016 | 0.8453 | 0.0055 ✓ |
| F42 | 0.6010 | 0.0083 | 0.0929 | 0.0005 | 0.8357 | 0.0086 | 1.0600 | 0.0272 | 0.0664 | 0.0024 | 0.8693 | 0.0085 ✓ |
| S43 | 0.6360 | 0.0098 | 0.0988 | 0.0006 | 0.8219 | 0.0086 | 1.2200 | 0.0359 | 0.0753 | 0.0027 | 0.8870 | 0.0079 ✓ |
| M44 | 0.6090 | 0.0716 | 0.0869 | 0.0036 | 0.6669 | 0.0529 | 1.0800 | 0.3880 | 0.0839 | 0.0295 | 0.9674 | 0.0730 ✓ |
| S45 | 0.6360 | 0.0097 | 0.0933 | 0.0005 | 0.8069 | 0.0086 | 1.1300 | 0.0260 | 0.0736 | 0.0019 | 0.8643 | 0.0068 ✓ |
| F46 | 0.6310 | 0.0081 | 0.0974 | 0.0005 | 0.8181 | 0.0087 | 1.1600 | 0.0312 | 0.0803 | 0.0022 | 0.8787 | 0.0074 ✓ |
| V47 | 0.6630 | 0.0135 | 0.0954 | 0.0007 | 0.7252 | 0.0138 | 1.2200 | 0.0309 | 0.0706 | 0.0034 | 0.8152 | 0.0125 ✓ |
| Q48 | 0.6060 | 0.0084 | 0.0965 | 0.0005 | 0.7434 | 0.0092 | 1.1000 | 0.0286 | 0.0731 | 0.0022 | 0.8474 | 0.0080 ✓ |
| G49 | 0.7300 | 0.0169 | 0.1300 | 0.0012 | 0.5568 | 0.0100 | 1.1500 | 0.0455 | 0.0948 | 0.0041 | 0.5899 | 0.0082 |
| E50 | 0.6380 | 0.0087 | 0.1000 | 0.0005 | 0.7409 | 0.0077 | 1.1200 | 0.0241 | 0.0770 | 0.0023 | 0.8481 | 0.0071 ✓ |
| E51 | 0.7040 | 0.0142 | 0.1560 | 0.0014 | 0.3626 | 0.0079 | 1.0100 | 0.0350 | 0.1210 | 0.0048 | 0.5182 | 0.0066 |

|  |  |  |  |  |  |  |  |  |  |  |  |  |  |
| --- | --- | --- | --- | --- | --- | --- | --- | --- | --- | --- | --- | --- | --- |
| S52 | — | — | — | — | — | — | 1.0000 | 0.0535 | — | — | — | — |  |
| D54 | 0.6960 | 0.0247 | 0.1260 | 0.0015 | 0.4957 | 0.0126 | 1.0300 | 0.0503 | 0.0933 | 0.0054 | 0.6122 | 0.0095 |  |
| K55 | 0.6650 | 0.0070 | 0.1250 | 0.0006 | 0.6160 | 0.0066 | 1.0900 | 0.0130 | 0.0955 | 0.0020 | 0.6520 | 0.0043 |  |
| I56 | 0.6430 | 0.0071 | 0.1040 | 0.0005 | 0.7408 | 0.0065 | 1.1500 | 0.0254 | 0.0842 | 0.0020 | 0.7976 | 0.0053 | ✓ |
| V58 | 0.5990 | 0.0079 | 0.0903 | 0.0005 | 0.8302 | 0.0094 | 1.0800 | 0.0201 | 0.0710 | 0.0023 | 0.8944 | 0.0079 | ✓ |
| A59 | 0.6250 | 0.0086 | 0.0943 | 0.0006 | 0.8257 | 0.0093 | 1.1200 | 0.0238 | 0.0736 | 0.0022 | 0.8955 | 0.0074 | ✓ |
| L60 | 0.7040 | 0.0158 | 0.1030 | 0.0008 | 0.8339 | 0.0124 | 1.3300 | 0.0417 | 0.0871 | 0.0046 | 0.8427 | 0.0100 | ✓ |
| G61 | 0.6280 | 0.0108 | 0.0951 | 0.0006 | 0.8051 | 0.0093 | 1.1800 | 0.0401 | 0.0697 | 0.0029 | 0.9031 | 0.0096 | ✓ |
| L62 | 0.6180 | 0.0156 | 0.0637 | 0.0006 | 0.7999 | 0.0159 | 1.1500 | 0.1240 | 0.0404 | 0.0059 | 0.9463 | 0.0322 |  |
| L67 | 0.6820 | 0.0233 | 0.0621 | 0.0008 | 0.8208 | 0.0188 | 1.2300 | 0.1740 | 0.0335 | 0.0060 | 0.8192 | 0.0296 |  |
| Y68 | 0.6250 | 0.0152 | 0.0780 | 0.0007 | 0.8027 | 0.0123 | 1.0700 | 0.0527 | 0.0520 | 0.0031 | 0.9033 | 0.0135 |  |
| L69 | 0.6070 | 0.1050 | 0.0952 | 0.0077 | 0.8076 | 0.1805 | 1.1100 | 0.3620 | 0.0772 | 0.0431 | 0.8574 | 0.1100 | ✓ |
| S70 | 0.6300 | 0.0100 | 0.0921 | 0.0005 | 0.8122 | 0.0087 | 1.1800 | 0.0250 | 0.0721 | 0.0018 | 0.8736 | 0.0066 | ✓ |
| C71 | 0.6680 | 0.0111 | 0.0940 | 0.0006 | 0.8004 | 0.0093 | 1.2900 | 0.0467 | 0.0740 | 0.0029 | 0.8627 | 0.0086 | ✓ |
| V72 | 0.6370 | 0.0094 | 0.0907 | 0.0005 | 0.8355 | 0.0089 | 1.0800 | 0.0175 | 0.0723 | 0.0016 | 0.8822 | 0.0063 | ✓ |
| L73 | 0.6700 | 0.0068 | 0.1020 | 0.0004 | 0.7372 | 0.0069 | 1.1800 | 0.0153 | 0.0835 | 0.0020 | 0.8753 | 0.0057 | ✓ |
| K74 | 0.6540 | 0.0069 | 0.1060 | 0.0005 | 0.7607 | 0.0068 | 1.1900 | 0.0199 | 0.0866 | 0.0022 | 0.8359 | 0.0055 | ✓ |
| D75 | 0.7510 | 0.1120 | 0.0983 | 0.0048 | 0.7861 | 0.0550 | 0.9760 | 0.3970 | 0.0777 | 0.0317 | 0.7861 | 0.0633 | ✓ |
| D76 | 0.6720 | 0.0083 | 0.0794 | 0.0003 | 0.7612 | 0.0072 | 1.1600 | 0.0255 | 0.0472 | 0.0012 | 0.8274 | 0.0059 |  |
| K77 | 0.6460 | 0.0072 | 0.1010 | 0.0004 | 0.7566 | 0.0067 | 1.1400 | 0.0156 | 0.0768 | 0.0013 | 0.7989 | 0.0042 | ✓ |
| T79 | 0.6520 | 0.0098 | 0.0917 | 0.0005 | 0.8001 | 0.0093 | 1.2000 | 0.0274 | 0.0691 | 0.0019 | 0.8743 | 0.0075 | ✓ |
| L80 | 0.6440 | 0.0084 | 0.0963 | 0.0005 | 0.8297 | 0.0084 | 1.1800 | 0.0198 | 0.0729 | 0.0018 | 0.8437 | 0.0062 | ✓ |
| Q81 | 0.6180 | 0.0084 | 0.0960 | 0.0005 | 0.7895 | 0.0083 | 1.1000 | 0.0266 | 0.0755 | 0.0025 | 0.8532 | 0.0078 | ✓ |
| L82 | 0.6270 | 0.0066 | 0.0949 | 0.0004 | 0.7408 | 0.0079 | 1.1300 | 0.0194 | 0.0697 | 0.0019 | 0.8271 | 0.0071 | ✓ |
| E83 | 0.6220 | 0.0111 | 0.0958 | 0.0007 | 0.7980 | 0.0094 | 1.1600 | 0.0302 | 0.0796 | 0.0028 | 0.8805 | 0.0079 | ✓ |
| S84 | 0.6340 | 0.0075 | 0.0981 | 0.0004 | 0.7966 | 0.0079 | 1.1300 | 0.0184 | 0.0763 | 0.0018 | 0.8580 | 0.0062 | ✓ |
| V85 | 0.6240 | 0.0075 | 0.0831 | 0.0004 | 0.7729 | 0.0089 | 1.1200 | 0.0223 | 0.0590 | 0.0019 | 0.8308 | 0.0085 | ✓ |
| D86 | 0.6290 | 0.0079 | 0.0834 | 0.0004 | 0.7385 | 0.0081 | 1.1300 | 0.0245 | 0.0545 | 0.0017 | 0.7993 | 0.0078 |  |
| K88 | 0.6440 | 0.0079 | 0.1110 | 0.0005 | 0.6731 | 0.0069 | 1.1000 | 0.0310 | 0.0815 | 0.0025 | 0.7580 | 0.0067 | ✓ |
| N89 | 0.6660 | 0.0143 | 0.1060 | 0.0008 | 0.6842 | 0.0109 | 1.1800 | 0.0351 | 0.0766 | 0.0028 | 0.7454 | 0.0082 | ✓ |
| Y90 | 0.6660 | 0.0120 | 0.0959 | 0.0006 | 0.7499 | 0.0099 | 1.1700 | 0.0323 | 0.0674 | 0.0025 | 0.7780 | 0.0085 | ✓ |
| K92 | 0.6260 | 0.0120 | 0.0905 | 0.0007 | 0.7301 | 0.0120 | 1.0600 | 0.0466 | 0.0621 | 0.0026 | 0.7771 | 0.0108 | ✓ |
| K93 | 0.6650 | 0.0453 | 0.0946 | 0.0018 | 0.6659 | 0.0249 | 1.1800 | 0.1290 | 0.0724 | 0.0076 | 0.7128 | 0.0201 | ✓ |
| K94 | 0.6420 | 0.0102 | 0.1020 | 0.0006 | 0.6741 | 0.0093 | 1.1700 | 0.0231 | 0.0789 | 0.0024 | 0.7819 | 0.0072 | ✓ |
| M95 | 0.6430 | 0.0719 | 0.1190 | 0.0055 | 0.5296 | 0.0665 | 1.1100 | 0.1660 | 0.0971 | 0.0235 | 0.7438 | 0.0529 |  |
| E96 | 0.6390 | 0.0084 | 0.1020 | 0.0005 | 0.7750 | 0.0081 | 1.1100 | 0.0238 | 0.0789 | 0.0022 | 0.8496 | 0.0064 | ✓ |
| K97 | 0.6450 | 0.0075 | 0.0944 | 0.0004 | 0.7861 | 0.0077 | 1.2200 | 0.0257 | 0.0752 | 0.0019 | 0.8267 | 0.0063 | ✓ |
| R98 | 0.6220 | 0.0075 | 0.0919 | 0.0004 | 0.8126 | 0.0078 | 1.1100 | 0.0210 | 0.0685 | 0.0015 | 0.8427 | 0.0058 | ✓ |
| F99 | 0.6270 | 0.0104 | 0.0880 | 0.0005 | 0.8354 | 0.0101 | 1.1200 | 0.0272 | 0.0668 | 0.0019 | 0.8712 | 0.0071 | ✓ |
| V100 | 0.6380 | 0.0102 | 0.0914 | 0.0005 | 0.7998 | 0.0096 | 1.1800 | 0.0229 | 0.0723 | 0.0020 | 0.8901 | 0.0065 | ✓ |
| F101 | 0.5970 | 0.0088 | 0.0959 | 0.0006 | 0.8185 | 0.0091 | 1.1000 | 0.0231 | 0.0717 | 0.0022 | 0.8877 | 0.0082 | ✓ |
| N102 | 0.6480 | 0.0129 | 0.0981 | 0.0008 | 0.8398 | 0.0091 | 1.2600 | 0.0394 | 0.0814 | 0.0031 | 0.9188 | 0.0080 | ✓ |
| K103 | 0.6350 | 0.0091 | 0.0960 | 0.0005 | 0.8135 | 0.0082 | 1.2100 | 0.0273 | 0.0739 | 0.0023 | 0.8773 | 0.0070 | ✓ |
| I104 | 0.6000 | 0.0091 | 0.0910 | 0.0006 | 0.8355 | 0.0083 | 1.1300 | 0.0295 | 0.0735 | 0.0025 | 0.9214 | 0.0067 | ✓ |
| E105 | 0.6590 | 0.0066 | 0.1050 | 0.0004 | 0.8103 | 0.0065 | 1.1800 | 0.0150 | 0.0774 | 0.0017 | 0.7992 | 0.0046 | ✓ |
| I106 | 0.6190 | 0.0060 | 0.1210 | 0.0005 | 0.5776 | 0.0056 | 0.9890 | 0.0144 | 0.0979 | 0.0023 | 0.7529 | 0.0045 | ✓ |
| N107 | 0.7260 | 0.0886 | 0.1040 | 0.0038 | 0.6342 | 0.0304 | 1.2700 | 0.2070 | 0.0820 | 0.0142 | 0.7852 | 0.0245 | ✓ |
| N108 | 0.6720 | 0.0222 | 0.0789 | 0.0007 | 0.7150 | 0.0137 | 1.2100 | 0.0595 | 0.0485 | 0.0023 | 0.7858 | 0.0110 |  |
| K109 | 0.6350 | 0.0063 | 0.1120 | 0.0005 | 0.6436 | 0.0055 | 1.0500 | 0.0144 | 0.0858 | 0.0016 | 0.7236 | 0.0044 | ✓ |
| L110 | 0.6130 | 0.0092 | 0.0935 | 0.0006 | 0.7804 | 0.0085 | 1.1100 | 0.0343 | 0.0705 | 0.0027 | 0.8751 | 0.0078 | ✓ |
| E111 | 0.6330 | 0.0098 | 0.0922 | 0.0006 | 0.8005 | 0.0080 | 1.1700 | 0.0291 | 0.0733 | 0.0022 | 0.8894 | 0.0061 | ✓ |
| F112 | 0.6810 | 0.0238 | 0.0930 | 0.0010 | 0.8102 | 0.0165 | 1.4000 | 0.0853 | 0.0714 | 0.0061 | 0.8363 | 0.0169 | ✓ |
| E113 | 0.6240 | 0.0108 | 0.0951 | 0.0007 | 0.8549 | 0.0087 | 1.1600 | 0.0284 | 0.0764 | 0.0022 | 0.9300 | 0.0064 | ✓ |
| S114 | 0.6280 | 0.0128 | 0.0911 | 0.0006 | 0.8468 | 0.0085 | 1.1200 | 0.0359 | 0.0741 | 0.0023 | 0.9230 | 0.0069 | ✓ |
| A115 | 0.6150 | 0.0101 | 0.0934 | 0.0005 | 0.8438 | 0.0095 | 1.0900 | 0.0302 | 0.0756 | 0.0026 | 0.8784 | 0.0077 | ✓ |
| Q116 | 0.6640 | 0.0156 | 0.0955 | 0.0008 | 0.8347 | 0.0132 | 1.2300 | 0.0434 | 0.0740 | 0.0029 | 0.8404 | 0.0088 | ✓ |
| F117 | 0.6690 | 0.0107 | 0.0954 | 0.0005 | 0.7930 | 0.0080 | 1.2600 | 0.0268 | 0.0764 | 0.0019 | 0.8693 | 0.0057 | ✓ |
| N119 | 0.6400 | 0.0106 | 0.1010 | 0.0007 | 0.8262 | 0.0080 | 1.2200 | 0.0317 | 0.0805 | 0.0028 | 0.9230 | 0.0070 | ✓ |
| W120 | 0.6590 | 0.0073 | 0.1440 | 0.0007 | 0.5797 | 0.0049 | 0.9640 | 0.0183 | 0.1110 | 0.0029 | 0.7880 | 0.0054 |  |
| Y121 | 0.6600 | 0.0122 | 0.0927 | 0.0006 | 0.8351 | 0.0081 | 1.2400 | 0.0365 | 0.0757 | 0.0025 | 0.9232 | 0.0068 | ✓ |
| I122 | 0.6370 | 0.0130 | 0.0931 | 0.0007 | 0.8377 | 0.0108 | 1.1900 | 0.0260 | 0.0694 | 0.0021 | 0.8845 | 0.0078 | ✓ |
| T124 | 0.6130 | 0.0160 | 0.0981 | 0.0009 | 0.8455 | 0.0127 | 1.1600 | 0.0573 | 0.0754 | 0.0040 | 0.9195 | 0.0115 | ✓ |
| S125 | 0.6580 | 0.0116 | 0.0932 | 0.0005 | 0.8545 | 0.0099 | 1.2800 | 0.0454 | 0.0691 | 0.0031 | 0.8724 | 0.0092 | ✓ |
| Q126 | 0.6150 | 0.0047 | 0.0956 | 0.0003 | 0.8055 | 0.0046 | 1.1300 | 0.0136 | 0.0732 | 0.0012 | 0.8996 | 0.0041 | ✓ |
| A127 | 0.6580 | 0.0069 | 0.1080 | 0.0004 | 0.6062 | 0.0059 | 1.0900 | 0.0172 | 0.0812 | 0.0019 | 0.6646 | 0.0050 | ✓ |
| E128 | 0.6670 | 0.0066 | 0.0981 | 0.0004 | 0.7853 | 0.0063 | 1.2200 | 0.0155 | 0.0757 | 0.0015 | 0.8915 | 0.0047 | ✓ |
| N129 | 0.6620 | 0.0090 | 0.0990 | 0.0005 | 0.7682 | 0.0080 | 1.2400 | 0.0239 | 0.0760 | 0.0020 | 0.7992 | 0.0062 | ✓ |
| M130 | 0.6680 | 0.1200 | 0.0999 | 0.0052 | 0.8075 | 0.0779 | 1.1600 | 0.2510 | 0.0777 | 0.0188 | 0.8401 | 0.0535 | ✓ |
| V132 | 0.6120 | 0.0065 | 0.1000 | 0.0005 | 0.8227 | 0.0074 | 1.1200 | 0.0167 | 0.0786 | 0.0019 | 0.8991 | 0.0065 | ✓ |
| F133 | 0.5740 | 0.0073 | 0.0924 | 0.0005 | 0.8479 | 0.0089 | 0.9790 | 0.0198 | 0.0698 | 0.0018 | 0.9225 | 0.0072 | ✓ |
| L134 | 0.6150 | 0.0073 | 0.0955 | 0.0005 | 0.8046 | 0.0084 | 1.1400 | 0.0191 | 0.0733 | 0.0020 | 0.9014 | 0.0066 | ✓ |
| G135 | 0.6160 | 0.0106 | 0.0934 | 0.0006 | 0.8327 | 0.0088 | 1.1400 | 0.0397 | 0.0732 | 0.0024 | 0.9068 | 0.0071 | ✓ |
| G136 | 0.6500 | 0.0103 | 0.1030 | 0.0006 | 0.8179 | 0.0098 | 1.1600 | 0.0372 | 0.0719 | 0.0034 | 0.8536 | 0.0096 | ✓ |
| T137 | 0.6450 | 0.0092 | 0.1130 | 0.0006 | 0.7322 | 0.0080 | 1.1400 | 0.0270 | 0.0882 | 0.0027 | 0.7633 | 0.0069 | ✓ |
| K138 | 0.6970 | 0.0300 | 0.1210 | 0.0018 | 0.5693 | 0.0150 | 1.1100 | 0.0629 | 0.0836 | 0.0052 | 0.6637 | 0.0110 | ✓ |
| G139 | 0.6380 | 0.2730 | 0.1140 | 0.0074 | 0.6218 | 0.0490 | 1.0800 | 0.3350 | 0.0885 | 0.0251 | 0.6819 | 0.0368 | ✓ |
| G140 | 0.6570 | 0.0111 | 0.1210 | 0.0007 | 0.6507 | 0.0079 | 1.0400 | 0.0270 | 0.0906 | 0.0029 | 0.7362 | 0.0064 | ✓ |
| Q141 | 0.6130 | 0.0096 | 0.0904 | 0.0006 | 0.8378 | 0.0082 | 1.0900 | 0.0460 | 0.0684 | 0.0024 | 0.9001 | 0.0072 | ✓ |
| D142 | 0.7020 | 0.0078 | 0.1110 | 0.0005 | 0.6252 | 0.0065 | 1.1700 | 0.0161 | 0.0807 | 0.0017 | 0.7088 | 0.0050 | ✓ |
| I143 | 0.6140 | 0.0148 | 0.0894 | 0.0008 | 0.8125 | 0.0121 | 1.0500 | 0.0529 | 0.0657 | 0.0045 | 0.8910 | 0.0121 | ✓ |
| T144 | 0.6310 | 0.0194 | – | – | 0.8048 | 0.0178 | 1.1600 | 0.0952 | 0.0258 | 0.0029 | 0.8578 | 0.0191 |  |
| D145 | 0.6120 | 0.0143 | 0.0673 | 0.0006 | 0.8069 | 0.0147 | 1.0600 | 0.0647 | 0.0427 | 0.0029 | 0.8749 | 0.0162 |  |
| F146 | 0.6410 | 0.0115 | 0.0883 | 0.0006 | 0.8468 | 0.0094 | 1.2600 | 0.0473 | 0.0693 | 0.0033 | 0.8961 | 0.0088 | ✓ |
| T147 | 0.6450 | 0.0087 | 0.1020 | 0.0006 | 0.8311 | 0.0080 | 1.2100 | 0.0348 | 0.0792 | 0.0027 | 0.8499 | 0.0071 | ✓ |
| M148 | 0.6340 | 0.0409 | 0.0959 | 0.0024 | 0.7661 | 0.0413 | 1.1300 | 0.1310 | 0.0716 | 0.0102 | 0.8230 | 0.0367 | ✓ |
| Q149 | 0.6370 | 0.0078 | 0.1020 | 0.0005 | 0.8293 | 0.0077 | 1.1700 | 0.0229 | 0.0821 | 0.0023 | 0.8395 | 0.0058 | ✓ |
| F150 | 0.6780 | 0.0101 | 0.1060 | 0.0005 | 0.7858 | 0.0084 | 1.2100 | 0.0230 | 0.0870 | 0.0022 | 0.8296 | 0.0061 | ✓ |
| V151 | – | – | – | – | – | – | 1.1400 | 0.0176 | – | – | – | – |  |
| S152 | 0.6600 | 0.0087 | 0.0946 | 0.0005 | 0.8107 | 0.0081 | 1.2700 | 0.0313 | 0.0723 | 0.0023 | 0.8525 | 0.0069 | ✓ |

<sup>15</sup>N backbone relaxation observables for sample 2 of IL-1 $\beta$ from 138 residues curated for resolution etc. Errors are derived from replicates in the individual T<sub>1</sub> and T<sub>2</sub> relaxation time courses and from S/N in spectra sampling the heteronuclear NOE. A tick in the final column marks the amides that entered the rotational-diffusion fit reported. Unticked amides were removed by orientation-aware flexibility, low-NOE and exchange screens.

**Table S3:**
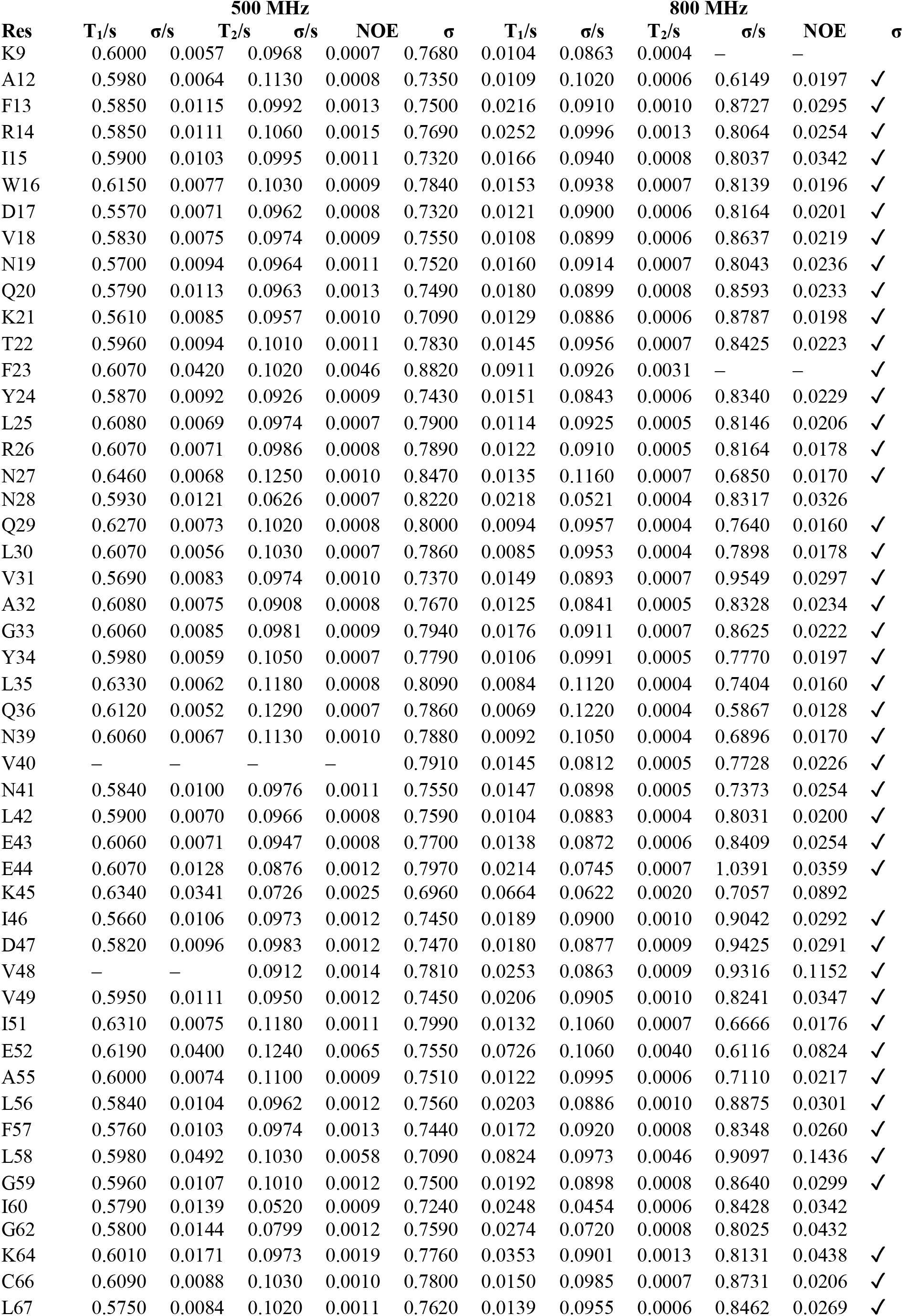
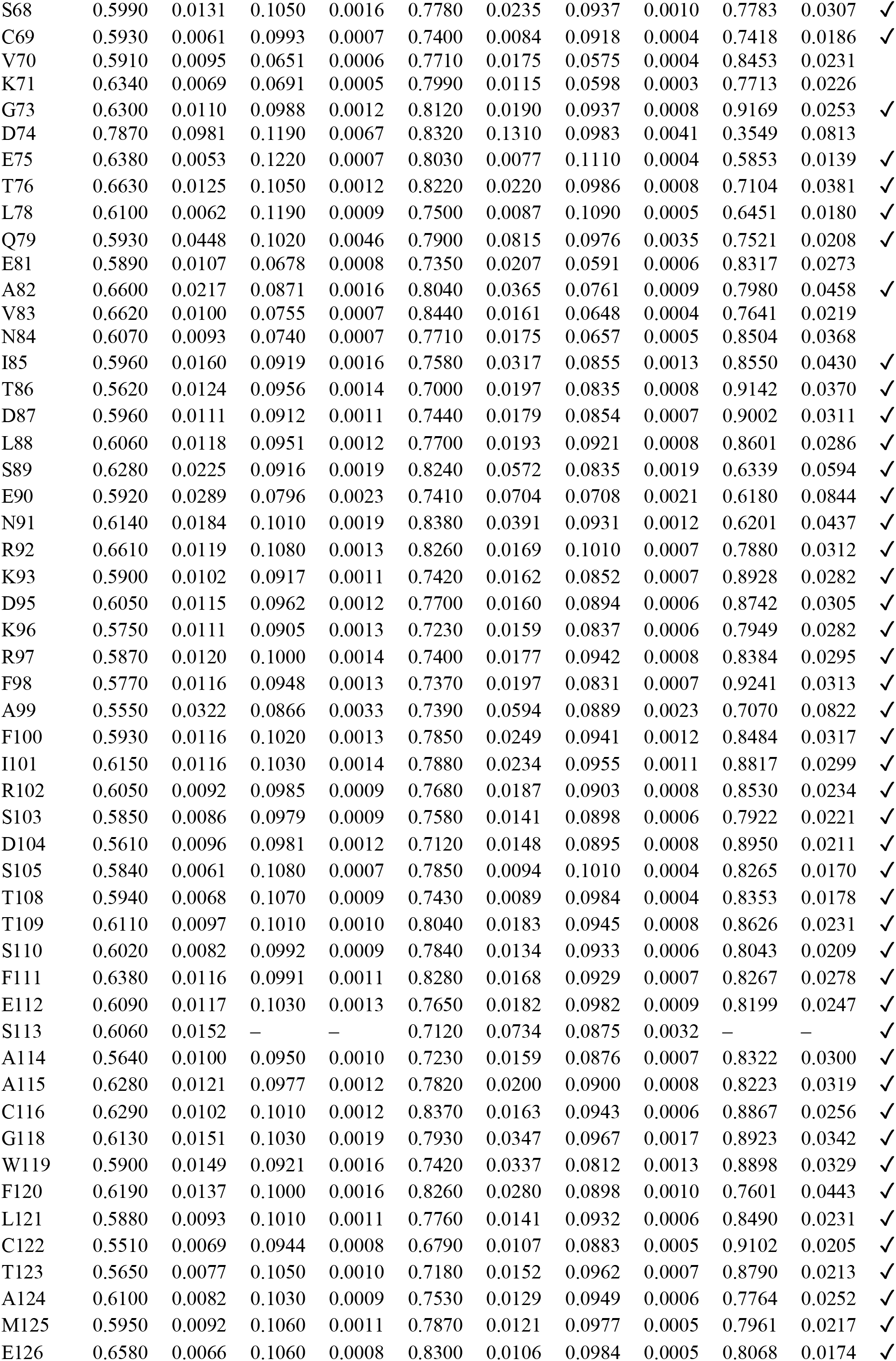
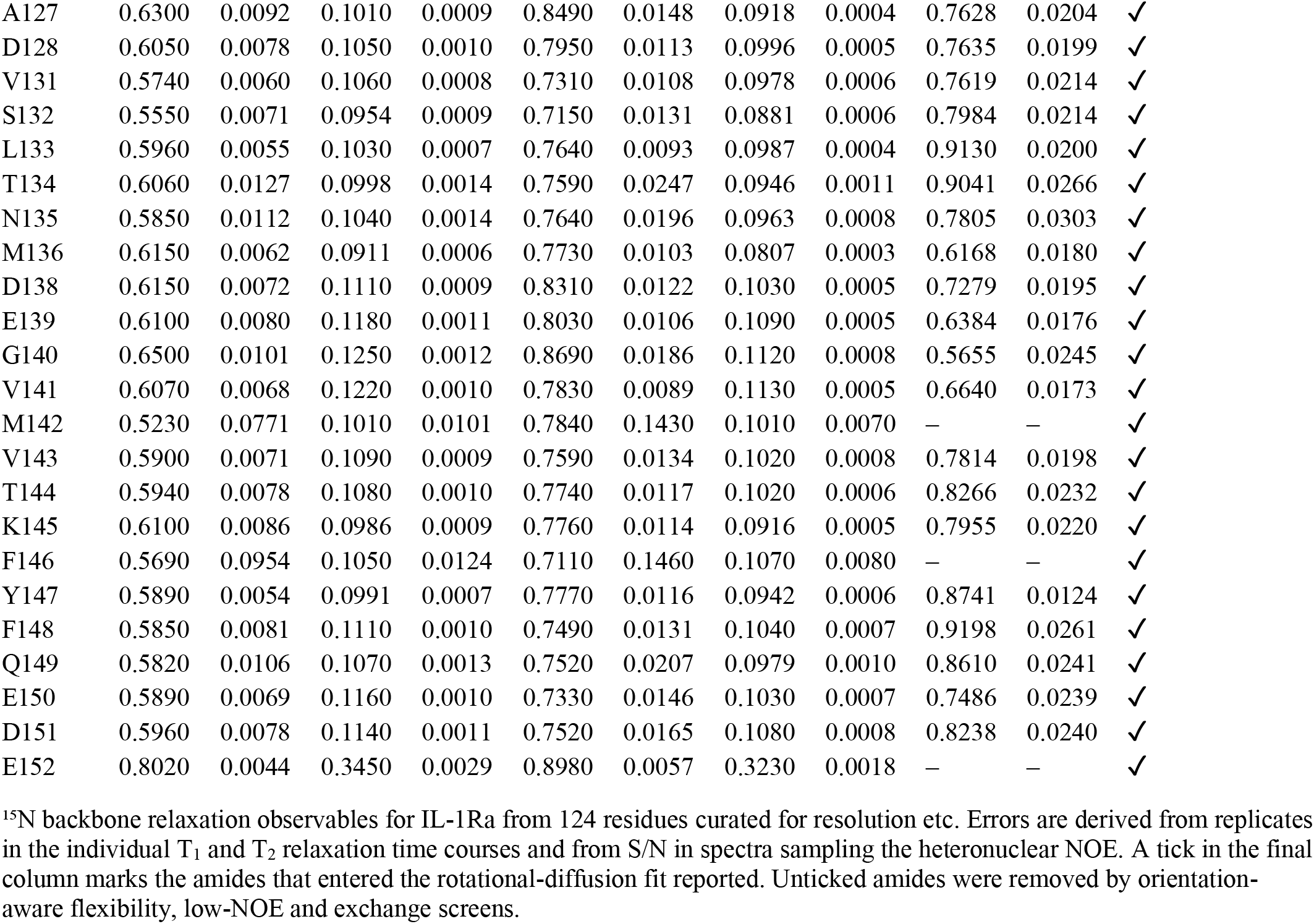
^15^N-relaxation used to determine the rotational diffusion tensor of IL-1Ra.

**Table S4:**
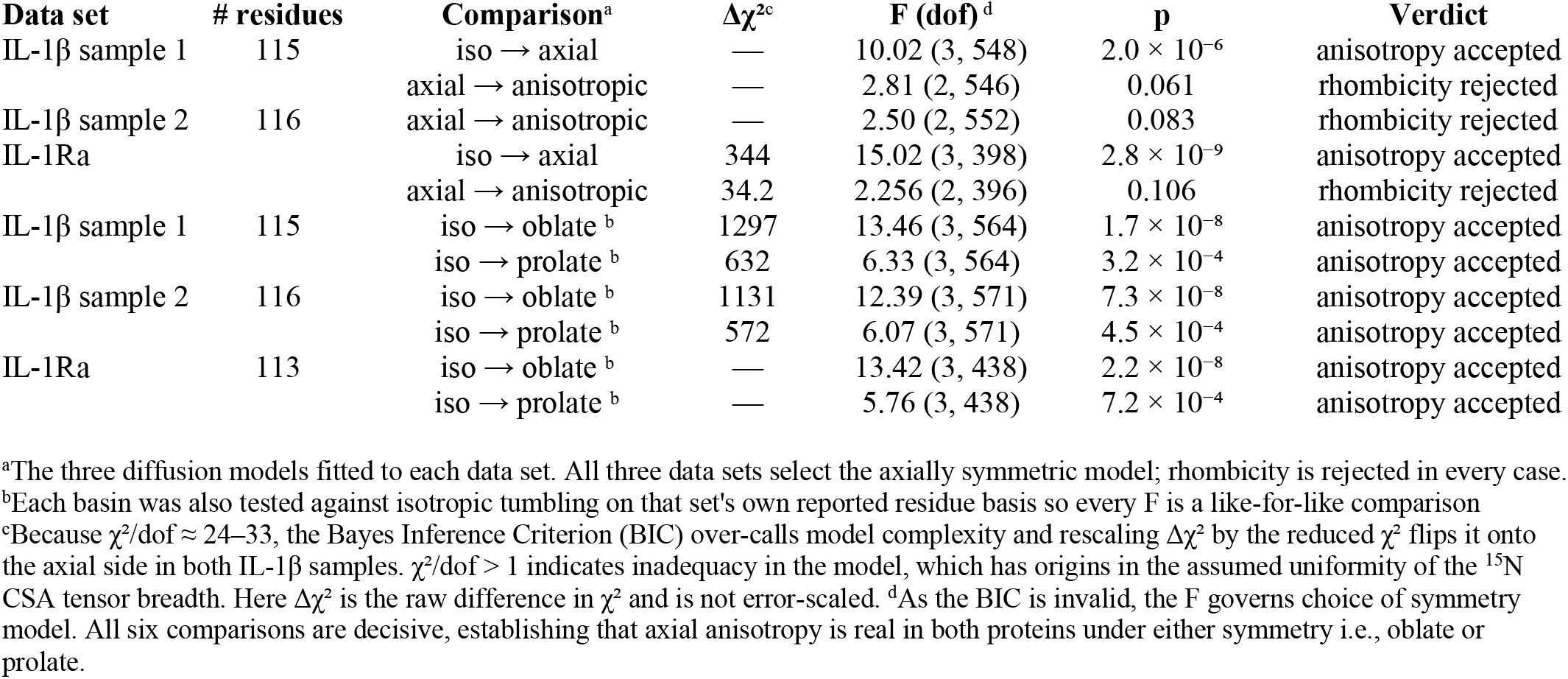
Symmetry ladder statistics for rotational diffusion tensor of IL-1β and IL-1Ra.

**Table S5:** Determinacy of the rotational tensor anisotropy symmetry.

| | IL-1β Sample 1 | IL-1β Sample 2 | IL-1Ra |
| --- | --- | --- | --- |
| axis separation, basin A vs B <sup>a</sup> | 87.5° | 86.2° | 85.6° |
| $\Delta\chi^2$ (prolate – oblate), raw <sup>b</sup> | 664.6 | 559.2 | 719.4 |
| $\Delta\chi^2$, error-scaled <sup>b</sup> | 20.7 | 18.4 | 22.1 |
| resamples selecting basin B <sup>c</sup> | 34% | 28% | 25% |
| $\lambda_{\text{max}}$ of the axis-orientation tensor <sup>c</sup> | 0.616 | 0.682 | 0.705 / 0.707 |
| axis cone, median / 90th percentile <sup>d</sup> | 24.0° / 84.5° | 15.9° / 82.5° | 18° / 81° |
| 90th ÷ median <sup>d</sup> | 3.5 | 5.2 | 4.6 |
| $\tau\text{c}$, eff difference between basins <sup>e</sup> | 0.013 ns (0.14%) | 0.013 ns (0.14%) | 0.062 ns (0.74%) |
| structural rhombicity $\rho$ <sup>f</sup> | 0.421 | 0.421 | 0.468 |
| N<sub>res</sub> / N<sub>obs</sub> <sup>g</sup> | 115 / 684 | 116 / 692 | 113 / 556 |

**Table S6:**
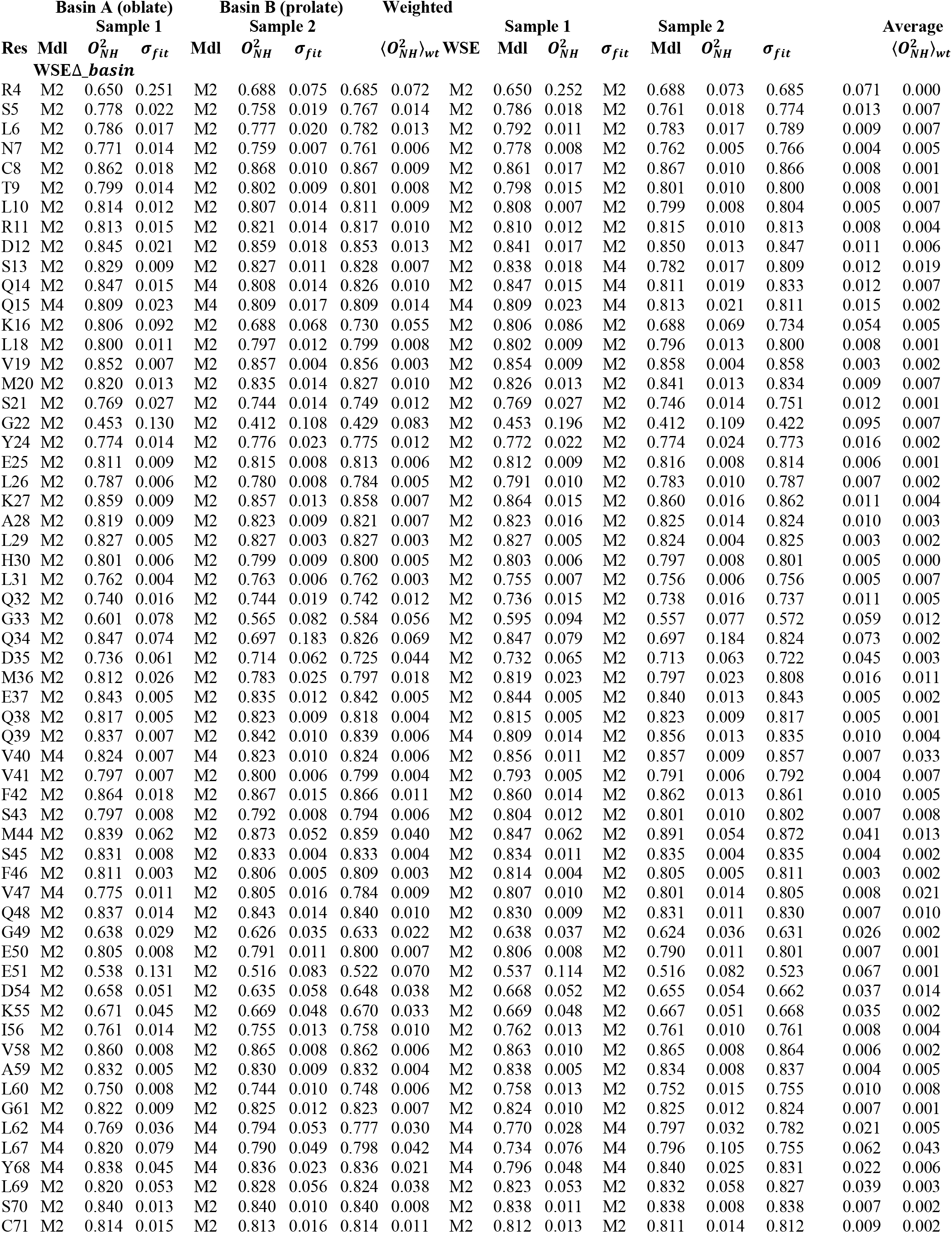
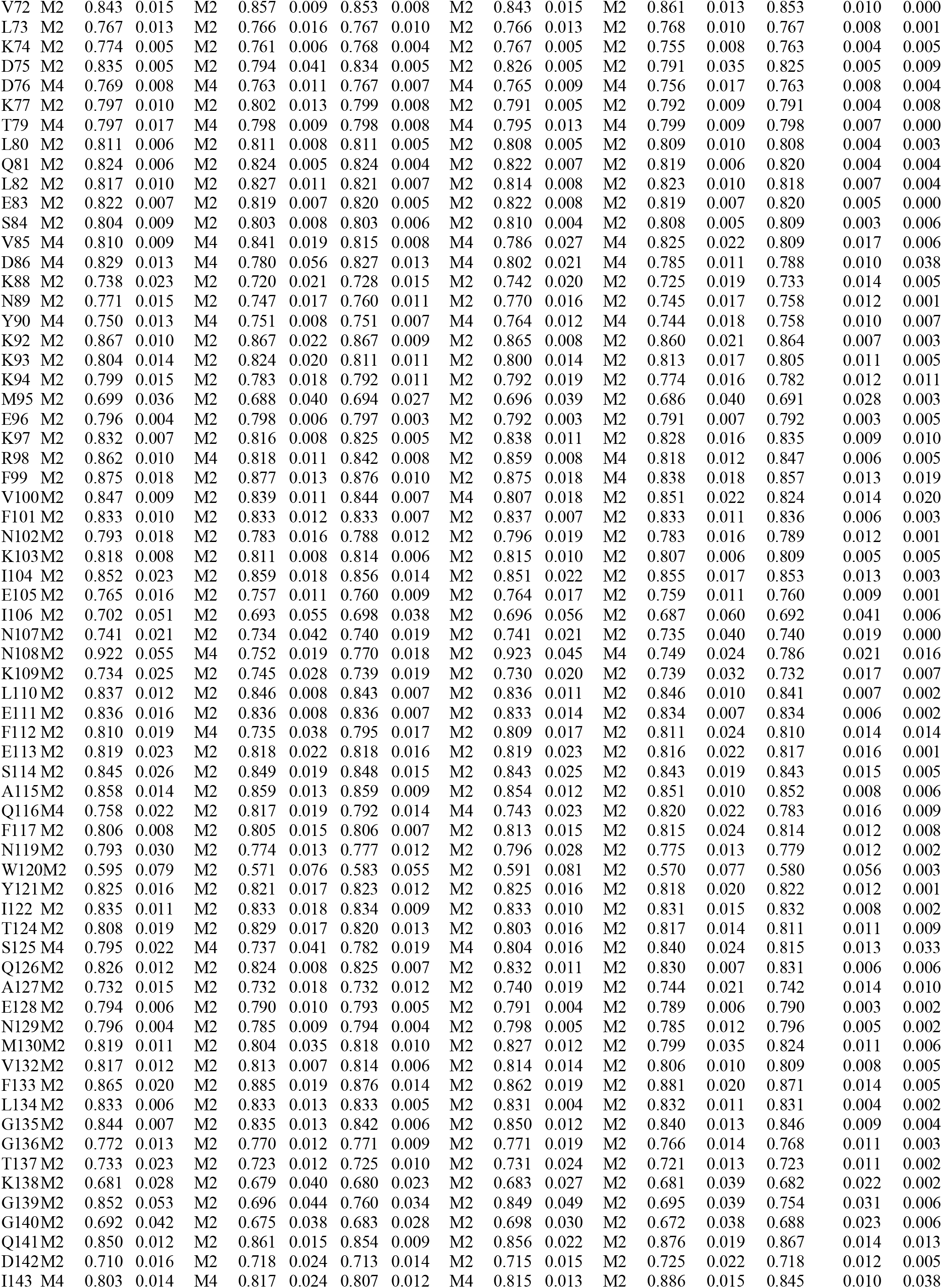
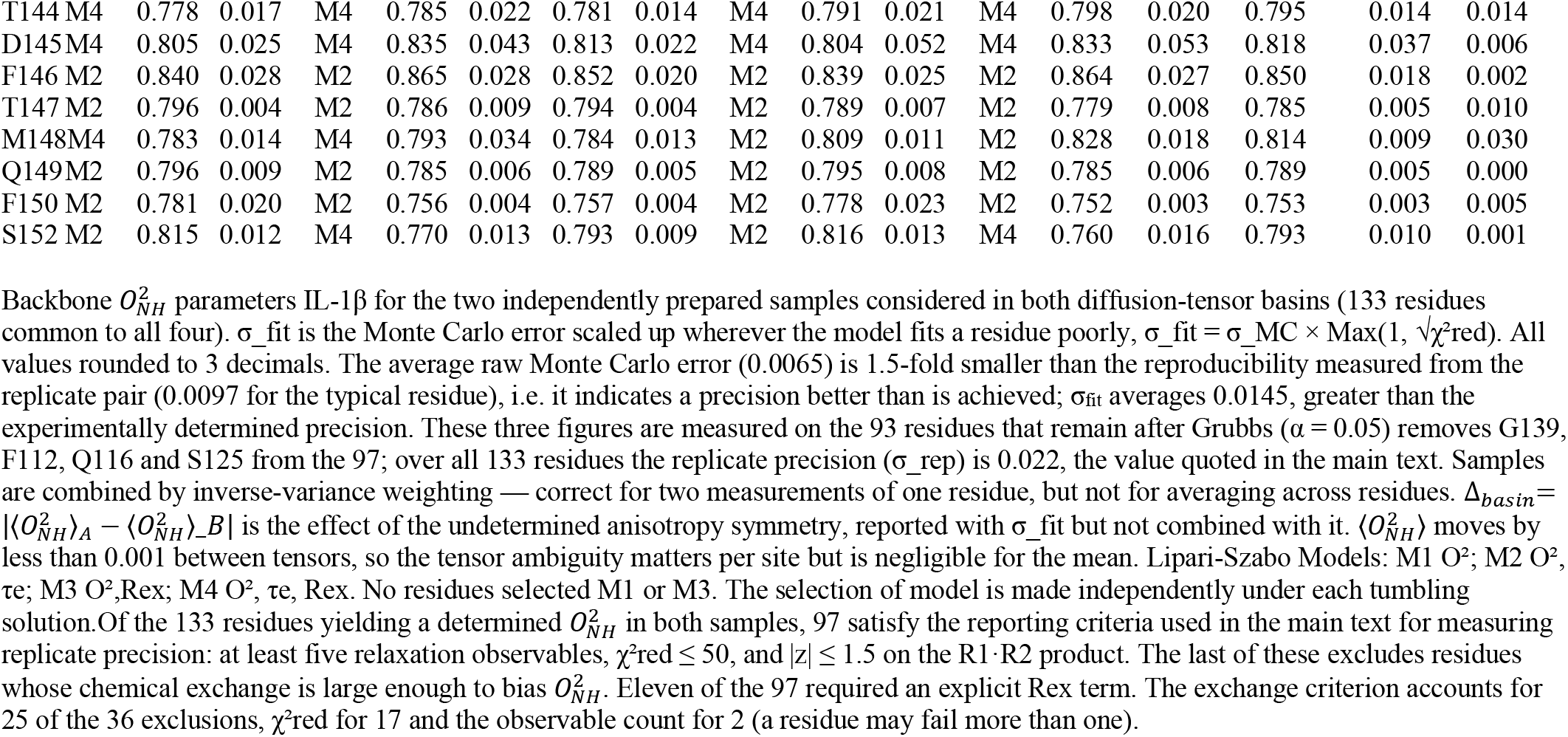
Dependence of IL-1β 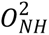 parameters on tensor symmetry.

**Table S7:** Dependence of IL-1Ra 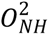 parameters on tensor symmetry.

| Res | Basin A (oblate) | | | Basin B (prolate) | | | Weighted Average<br>$\Delta_{\text{basin}}$ |
| --- | --- | --- | --- | --- | --- | --- | --- |
| | Mdl | $O_{NH}^2$ | $\sigma_{fit}$ | Mdl | $O_{NH}^2$ | $\sigma_{fit}$ | |
| A12 | M2 | 0.736 | 0.023 | M2 | 0.745 | 0.018 | 0.009 |
| F13 | M2 | 0.847 | 0.013 | M2 | 0.847 | 0.013 | 0.000 |
| R14 | M2 | 0.800 | 0.008 | M2 | 0.801 | 0.008 | 0.001 |
| I15 | M2 | 0.830 | 0.006 | M2 | 0.830 | 0.006 | 0.000 |
| W16 | M2 | 0.801 | 0.007 | M2 | 0.807 | 0.008 | 0.006 |
| D17 | M2 | 0.859 | 0.007 | M2 | 0.870 | 0.005 | 0.011 |
| V18 | M2 | 0.845 | 0.006 | M2 | 0.840 | 0.009 | 0.005 |
| N19 | M2 | 0.874 | 0.022 | M2 | 0.860 | 0.008 | 0.014 |
| Q20 | M2 | 0.846 | 0.008 | M2 | 0.857 | 0.007 | 0.011 |
| K21 | M2 | 0.867 | 0.008 | M2 | 0.857 | 0.017 | 0.010 |
| T22 | M2 | 0.805 | 0.011 | M2 | 0.820 | 0.006 | 0.015 |
| F23* | M2 | 0.802 | 0.026 | M2 | 0.819 | 0.029 | 0.017 |
| Y24 | M2 | 0.877 | 0.011 | M2 | 0.877 | 0.011 | 0.000 |
| L25 | M2 | 0.825 | 0.004 | M2 | 0.822 | 0.004 | 0.003 |
| R26 | M2 | 0.824 | 0.004 | M2 | 0.821 | 0.004 | 0.003 |
| N27 | M2 | 0.667 | 0.029 | M2 | 0.660 | 0.034 | 0.007 |
| N28 | M4 | 0.825 | 0.015 | M4 | 0.822 | 0.014 | 0.003 |
| Q29 | M2 | 0.796 | 0.005 | M2 | 0.798 | 0.006 | 0.002 |
| L30 | M2 | 0.810 | 0.006 | M2 | 0.803 | 0.004 | 0.007 |
| V31 | M2 | 0.852 | 0.014 | M2 | 0.843 | 0.019 | 0.009 |
| A32 | M4 | 0.818 | 0.010 | M4 | 0.823 | 0.014 | 0.005 |
| G33 | M2 | 0.829 | 0.007 | M2 | 0.838 | 0.014 | 0.009 |
| Y34 | M2 | 0.776 | 0.015 | M2 | 0.799 | 0.004 | 0.023 |
| L35 | M2 | 0.694 | 0.022 | M2 | 0.713 | 0.010 | 0.019 |
| Q36 | M2 | 0.665 | 0.025 | M2 | 0.660 | 0.020 | 0.005 |
| N39 | M2 | 0.764 | 0.005 | M2 | 0.753 | 0.007 | 0.011 |
| V40* | M2 | 0.917 | 0.065 | M2 | 0.912 | 0.051 | 0.005 |
| N41 | M2 | 0.837 | 0.005 | M2 | 0.851 | 0.009 | 0.014 |
| L42 | M4 | 0.810 | 0.015 | M4 | 0.801 | 0.010 | 0.009 |
| E43 | M4 | 0.814 | 0.010 | M2 | 0.849 | 0.009 | 0.035 |
| E44 | M2 | 0.928 | 0.080 | M2 | 0.924 | 0.064 | 0.004 |
| K45 | M4 | 0.784 | 0.050 | M4 | 0.776 | 0.045 | 0.008 |
| I46 | M2 | 0.872 | 0.020 | M2 | 0.862 | 0.012 | 0.010 |
| D47 | M2 | 0.849 | 0.016 | M2 | 0.856 | 0.020 | 0.007 |
| V48* | M2 | 0.877 | 0.014 | M2 | 0.878 | 0.013 | 0.001 |
| V49 | M2 | 0.846 | 0.007 | M2 | 0.852 | 0.008 | 0.006 |
| I51 | M2 | 0.739 | 0.012 | M2 | 0.729 | 0.011 | 0.010 |
| E52 | M2 | 0.714 | 0.028 | M2 | 0.705 | 0.031 | 0.009 |
| A55 | M2 | 0.762 | 0.015 | M2 | 0.779 | 0.010 | 0.017 |
| L56 | M2 | 0.847 | 0.008 | M2 | 0.853 | 0.008 | 0.006 |
| F57 | M2 | 0.837 | 0.010 | M2 | 0.847 | 0.007 | 0.010 |
| L58 | M2 | 0.806 | 0.030 | M2 | 0.804 | 0.031 | 0.002 |
| G59 | M2 | 0.838 | 0.013 | M2 | 0.837 | 0.013 | 0.001 |
| I60 | M4 | 0.858 | 0.100 | M4 | 0.864 | 0.033 | 0.006 |
| G62 | M4 | 0.805 | 0.021 | M4 | 0.848 | 0.038 | 0.043 |
| K64 | M4 | 0.803 | 0.021 | M2 | 0.841 | 0.011 | 0.038 |
| C66 | M2 | 0.815 | 0.019 | M2 | 0.805 | 0.007 | 0.010 |
| L67 | M2 | 0.820 | 0.008 | M2 | 0.824 | 0.006 | 0.004 |
| S68 | M2 | 0.794 | 0.014 | M2 | 0.814 | 0.013 | 0.020 |
| C69 | M2 | 0.823 | 0.006 | M2 | 0.844 | 0.014 | 0.021 |
| V70 | M4 | 0.872 | 0.147 | M4 | 0.783 | 0.190 | 0.089 |
| K71 | M4 | 0.720 | 0.224 | M4 | 0.722 | 0.147 | 0.002 |
| G73 | M2 | 0.819 | 0.020 | M2 | 0.823 | 0.024 | 0.004 |
| D74 | M2 | 0.686 | 0.039 | M2 | 0.701 | 0.047 | 0.015 |
| E75 | M2 | 0.716 | 0.009 | M2 | 0.699 | 0.012 | 0.017 |
| T76 | M4 | 0.700 | 0.027 | M2 | 0.785 | 0.020 | 0.085 |
| L78 | M2 | 0.712 | 0.027 | M2 | 0.724 | 0.017 | 0.012 |
| Q79 | M2 | 0.793 | 0.026 | M2 | 0.807 | 0.026 | 0.014 |
| E81 | M4 | 0.851 | 0.020 | M4 | 0.852 | 0.063 | 0.001 |
| A82 | M4 | 0.760 | 0.027 | M4 | 0.766 | 0.036 | 0.006 |
| V83 | M4 | 0.652 | 0.232 | M4 | 0.739 | 0.014 | 0.087 |
| N84 | M4 | 0.780 | 0.016 | M4 | 0.770 | 0.094 | 0.010 |
| I85 | M2 | 0.872 | 0.014 | M2 | 0.872 | 0.013 | 0.000 |
| T86 | M2 | 0.885 | 0.019 | M2 | 0.886 | 0.019 | 0.001 |
| D87 | M2 | 0.878 | 0.015 | M2 | 0.891 | 0.028 | 0.013 |
| L88 | M2 | 0.843 | 0.013 | M2 | 0.837 | 0.009 | 0.006 |
| S89 | M4 | 0.776 | 0.030 | M4 | 0.757 | 0.029 | 0.019 |
| E90 | M4 | 0.823 | 0.043 | M4 | 0.830 | 0.044 | 0.007 |
| N91 | M2 | 0.804 | 0.015 | M2 | 0.792 | 0.010 | 0.012 |
| R92 | M4 | 0.715 | 0.014 | M2 | 0.769 | 0.011 | 0.054 |
| K93 | M2 | 0.891 | 0.030 | M4 | 0.831 | 0.020 | 0.060 |
| D95 | M2 | 0.847 | 0.011 | M4 | 0.793 | 0.015 | 0.054 |
| K96 | M4 | 0.849 | 0.016 | M4 | 0.855 | 0.015 | 0.006 |
| R97 | M4 | 0.790 | 0.017 | M2 | 0.838 | 0.007 | 0.048 |
| F98 | M2 | 0.895 | 0.030 | M2 | 0.896 | 0.030 | 0.001 |
| A99 | M2 | 0.901 | 0.028 | M2 | 0.891 | 0.023 | 0.010 |
| F100 | M2 | 0.809 | 0.009 | M2 | 0.815 | 0.007 | 0.006 |
| I101 | M2 | 0.796 | 0.008 | M2 | 0.806 | 0.010 | 0.010 |
| R102 | M2 | 0.836 | 0.011 | M4 | 0.802 | 0.012 | 0.034 |
| S103 | M2 | 0.835 | 0.006 | M2 | 0.851 | 0.009 | 0.016 |
| D104 | M2 | 0.849 | 0.020 | M2 | 0.860 | 0.011 | 0.011 |
| S105 | M2 | 0.767 | 0.017 | M2 | 0.767 | 0.017 | 0.000 |
| T108 | M2 | 0.804 | 0.007 | M2 | 0.787 | 0.017 | 0.017 |
| T109 | M2 | 0.803 | 0.006 | M2 | 0.818 | 0.013 | 0.015 |
| S110 | M2 | 0.815 | 0.005 | M2 | 0.833 | 0.014 | 0.018 |
| F111 | M2 | 0.810 | 0.012 | M4 | 0.763 | 0.020 | 0.047 |
| E112 | M2 | 0.788 | 0.012 | M2 | 0.808 | 0.007 | 0.020 |
| S113* | M2 | 0.820 | 0.031 | M2 | 0.812 | 0.039 | 0.008 |
| A114 | M2 | 0.872 | 0.007 | M2 | 0.857 | 0.010 | 0.015 |
| A115 | M2 | 0.825 | 0.010 | M2 | 0.820 | 0.006 | 0.005 |
| C116 | M2 | 0.801 | 0.010 | M2 | 0.798 | 0.008 | 0.003 |
| G118 | M2 | 0.793 | 0.010 | M2 | 0.798 | 0.010 | 0.005 |
| W119 | M4 | 0.821 | 0.025 | M4 | 0.821 | 0.026 | 0.000 |
| F120 | M4 | 0.765 | 0.018 | M4 | 0.767 | 0.018 | 0.002 |
| L121 | M2 | 0.815 | 0.010 | M2 | 0.835 | 0.012 | 0.020 |
| C122 | M2 | 0.903 | 0.018 | M2 | 0.892 | 0.011 | 0.011 |
| T123 | M2 | 0.833 | 0.008 | M2 | 0.820 | 0.014 | 0.013 |
| A124 | M2 | 0.821 | 0.014 | M2 | 0.804 | 0.006 | 0.017 |
| M125 | M2 | 0.779 | 0.015 | M2 | 0.777 | 0.016 | 0.002 |
| E126 | M2 | 0.773 | 0.012 | M2 | 0.767 | 0.005 | 0.006 |
| A127 | M4 | 0.732 | 0.023 | M4 | 0.757 | 0.027 | 0.025 |
| D128 | M2 | 0.773 | 0.010 | M2 | 0.791 | 0.004 | 0.018 |
| V131 | M2 | 0.806 | 0.008 | M2 | 0.788 | 0.020 | 0.018 |
| S132 | M4 | 0.832 | 0.011 | M2 | 0.883 | 0.008 | 0.051 |
| L133 | M2 | 0.795 | 0.013 | M2 | 0.807 | 0.007 | 0.012 |
| T134 | M2 | 0.812 | 0.012 | M2 | 0.827 | 0.014 | 0.015 |
| N135 | M2 | 0.792 | 0.016 | M2 | 0.815 | 0.007 | 0.023 |
| M136 | M4 | 0.766 | 0.010 | M4 | 0.770 | 0.004 | 0.004 |
| D138 | M2 | 0.755 | 0.008 | M2 | 0.744 | 0.013 | 0.011 |
| E139 | M2 | 0.729 | 0.008 | M2 | 0.716 | 0.015 | 0.013 |
| G140 | M2 | 0.689 | 0.012 | M2 | 0.684 | 0.012 | 0.005 |
| V141 | M2 | 0.699 | 0.019 | M2 | 0.702 | 0.024 | 0.003 |
| M142* | M2 | 0.757 | 0.063 | M2 | 0.753 | 0.063 | 0.004 |
| V143 | M2 | 0.776 | 0.014 | M2 | 0.774 | 0.016 | 0.002 |
| T144 | M2 | 0.802 | 0.011 | M2 | 0.783 | 0.008 | 0.019 |
| K145 | M2 | 0.832 | 0.008 | M4 | 0.795 | 0.011 | 0.037 |
| F146* | M2 | 0.709 | 0.069 | M2 | 0.735 | 0.067 | 0.026 |
| Y147 | M2 | 0.823 | 0.010 | M2 | 0.833 | 0.010 | 0.010 |
| F148 | M2 | 0.774 | 0.018 | M2 | 0.771 | 0.020 | 0.003 |
| Q149 | M2 | 0.786 | 0.022 | M2 | 0.802 | 0.010 | 0.016 |
| E150 | M2 | 0.760 | 0.015 | M2 | 0.750 | 0.025 | 0.010 |
| D151 | M2 | 0.749 | 0.019 | M2 | 0.740 | 0.026 | 0.009 |
| E152* | M2 | 0.178 | 0.007 | M2 | 0.173 | 0.005 | 0.005 |

**Table S8:** Sample 1 IL-1β 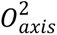 parameters.

| Methyl | basin A (oblate) | | | basin B (prolate) | | | $\Delta_{\text{basin}}$ |
| --- | --- | --- | --- | --- | --- | --- | --- |
| | $O_{axis}^2$ | $\sigma$ | $\tau c, \text{eff (ns)}$ | $O_{axis}^2$ | $\sigma$ | $\tau c, \text{eff (ns)}$ | |
| V3 CG1 | 0.2339 | 0.0040 | 8.6816 | 0.2286 | 0.0040 | 8.8827 | 0.0053 |
| V3 CG2 | 0.2061 | 0.0040 | 8.9430 | 0.2026 | 0.0039 | 9.0974 | 0.0035 |
| L6 CD1 | 0.1887 | 0.0028 | 9.1525 | 0.1873 | 0.0028 | 9.2235 | 0.0014 |
| L6 CD2 | 0.1965 | 0.0039 | 8.7461 | 0.1936 | 0.0038 | 8.8762 | 0.0029 |
| L10 CD1 | 0.6405 | 0.0081 | 9.1441 | 0.6612 | 0.0083 | 8.8584 | 0.0207 |
| L10 CD2 | 0.6139 | 0.0119 | 8.7553 | 0.6022 | 0.0116 | 8.9244 | 0.0117 |
| L18 CD1 | 0.3697 | 0.0041 | 9.1462 | 0.3746 | 0.0041 | 9.0275 | 0.0049 |
| L18 CD2 | 0.3789 | 0.0058 | 9.0926 | 0.3765 | 0.0058 | 9.1512 | 0.0024 |
| V19 CG1 | 0.6292 | 0.0147 | 9.0486 | 0.6427 | 0.0150 | 8.8584 | 0.0135 |
| V19 CG2 | 0.5937 | 0.0080 | 9.1356 | 0.5931 | 0.0080 | 9.1434 | 0.0006 |
| M20 CE | 0.0242 | 0.0033 | 8.8002 | 0.0238 | 0.0032 | 8.9510 | 0.0004 |
| L26 CD1 | 0.4862 | 0.0083 | 8.7907 | 0.4768 | 0.0082 | 8.9637 | 0.0094 |
| L26 CD2 | 0.5905 | 0.0116 | 9.0101 | 0.5996 | 0.0118 | 8.8734 | 0.0091 |
| L29 CD1 | 0.3433 | 0.0059 | 9.1489 | 0.3544 | 0.0061 | 8.8626 | 0.0111 |
| L29 CD2 | 0.3667 | 0.0057 | 8.7355 | 0.3616 | 0.0056 | 8.8588 | 0.0051 |
| L31 CD1 | 0.7065 | 0.0183 | 8.9794 | 0.7024 | 0.0182 | 9.0311 | 0.0041 |
| L31 CD2 | 0.7557 | 0.0100 | 9.1283 | 0.7538 | 0.0099 | 9.1509 | 0.0019 |
| M36 CE | 0.1923 | 0.0032 | 9.0694 | 0.1945 | 0.0033 | 8.9661 | 0.0022 |
| V40 CG1 | 0.2858 | 0.0047 | 9.1242 | 0.2859 | 0.0047 | 9.1212 | 0.0001 |
| V40 CG2 | 0.3000 | 0.0059 | 8.9751 | 0.2972 | 0.0059 | 9.0596 | 0.0028 |
| V41 CG1 | 0.8288 | 0.0140 | 9.0182 | 0.8201 | 0.0138 | 9.1131 | 0.0087 |
| V41 CG2 | 0.8687 | 0.0111 | 9.1536 | 0.8863 | 0.0113 | 8.9722 | 0.0176 |
| M44 CE | 0.3674 | 0.0089 | 9.1517 | 0.3648 | 0.0088 | 9.2185 | 0.0026 |
| V47 CG1 | 0.7518 | 0.0290 | 8.7910 | 0.7437 | 0.0286 | 8.8865 | 0.0081 |
| V47 CG2 | 0.7413 | 0.0206 | 9.0016 | 0.7331 | 0.0204 | 9.1028 | 0.0082 |
| I56 CD1 | 0.2564 | 0.0020 | 9.0691 | 0.2539 | 0.0020 | 9.1577 | 0.0025 |
| V58 CG1 | 0.9115 | 0.0194 | 8.8901 | 0.9101 | 0.0194 | 8.9041 | 0.0014 |
| V58 CG2 | 0.8460 | 0.0134 | 9.1038 | 0.8408 | 0.0134 | 9.1599 | 0.0052 |
| L60 CD1 | 0.5261 | 0.0092 | 9.0545 | 0.5208 | 0.0092 | 9.1464 | 0.0053 |
| L60 CD2 | 0.5723 | 0.0064 | 9.1387 | 0.5897 | 0.0066 | 8.8694 | 0.0174 |
| L62 CD1 | 0.2415 | 0.0028 | 8.8924 | 0.2423 | 0.0028 | 8.8634 | 0.0008 |
| L62 CD2 | 0.2422 | 0.0029 | 8.8662 | 0.2393 | 0.0029 | 8.9728 | 0.0029 |
| L67 CD1 | 0.5398 | 0.0108 | 9.1360 | 0.5428 | 0.0109 | 9.0857 | 0.0030 |
| L67 CD2 | 0.4599 | 0.0082 | 9.1376 | 0.4734 | 0.0085 | 8.8774 | 0.0135 |
| L69 CD1 | 0.2671 | 0.0050 | 8.7637 | 0.2635 | 0.0049 | 8.8830 | 0.0036 |
| L69 CD2 | 0.2462 | 0.0035 | 9.0531 | 0.2448 | 0.0034 | 9.1070 | 0.0014 |
| V72 CG1 | 0.5066 | 0.0097 | 9.1130 | 0.5074 | 0.0097 | 9.0975 | 0.0008 |
| V72 CG2 | 0.5004 | 0.0068 | 9.0943 | 0.4999 | 0.0068 | 9.1042 | 0.0005 |
| L73 CD1 | 0.3019 | 0.0060 | 9.1534 | 0.3097 | 0.0062 | 8.9213 | 0.0078 |
| L73 CD2 | 0.3270 | 0.0052 | 8.9289 | 0.3213 | 0.0051 | 9.0883 | 0.0057 |
| L80 CD1 | 0.5960 | 0.0153 | 9.0384 | 0.5898 | 0.0151 | 9.1334 | 0.0062 |
| L80 CD2 | 0.6507 | 0.0113 | 9.0852 | 0.6607 | 0.0115 | 8.9480 | 0.0100 |
| L82 CD1 | 0.2160 | 0.0035 | 9.0267 | 0.2142 | 0.0034 | 9.1008 | 0.0018 |
| L82 CD2 | 0.1918 | 0.0026 | 8.9074 | 0.1888 | 0.0026 | 9.0477 | 0.0030 |
| V85 CG1 | 0.8297 | 0.0125 | 8.8884 | 0.8184 | 0.0123 | 9.0110 | 0.0113 |
| V85 CG2 | 0.8067 | 0.0105 | 9.1468 | 0.8328 | 0.0108 | 8.8595 | 0.0261 |
| M95 CE | 0.4350 | 0.0123 | 9.1530 | 0.4322 | 0.0122 | 9.2126 | 0.0028 |
| V100 CG1 | 0.9503 | 0.0281 | 8.7210 | 0.9346 | 0.0276 | 8.8674 | 0.0157 |
| V100 CG2 | 0.8930 | 0.0266 | 8.9673 | 0.8785 | 0.0261 | 9.1153 | 0.0145 |
| I104 CD1 | 0.4061 | 0.0079 | 9.1554 | 0.4138 | 0.0080 | 8.9854 | 0.0077 |
| I106 CD1 | 0.2610 | 0.0025 | 9.1556 | 0.2599 | 0.0025 | 9.1966 | 0.0011 |
| L110 CD1 | 0.2255 | 0.0039 | 8.7501 | 0.2214 | 0.0038 | 8.9113 | 0.0041 |
| L110 CD2 | 0.2272 | 0.0032 | 8.9632 | 0.2248 | 0.0032 | 9.0588 | 0.0024 |
| I122 CD1 | 0.7573 | 0.0111 | 9.0865 | 0.7663 | 0.0113 | 8.9807 | 0.0090 |
| M130 CE | 0.1356 | 0.0015 | 9.1137 | 0.1394 | 0.0015 | 8.8651 | 0.0038 |
| V132 CG1 | 0.8867 | 0.0202 | 8.8493 | 0.8729 | 0.0199 | 8.9893 | 0.0138 |
| V132 CG2 | 0.8323 | 0.0219 | 9.1555 | 0.8576 | 0.0225 | 8.8855 | 0.0253 |
| L134 CD1 | 0.6160 | 0.0169 | 9.1556 | 0.6310 | 0.0173 | 8.9372 | 0.0150 |
| L134 CD2 | 0.6930 | 0.0169 | 9.0909 | 0.7000 | 0.0170 | 8.9997 | 0.0070 |
| I143 CD1 | 0.5539 | 0.0142 | 9.0551 | 0.5637 | 0.0145 | 8.8980 | 0.0098 |
| M148 CE | 0.2762 | 0.0104 | 8.6217 | 0.2688 | 0.0101 | 8.8588 | 0.0074 |
| V151 CG1 | 0.3608 | 0.0055 | 9.0710 | 0.3569 | 0.0054 | 9.1719 | 0.0039 |
| V151 CG2 | 0.3735 | 0.0076 | 9.0878 | 0.3809 | 0.0078 | 8.9118 | 0.0074 |

Methyl symmetry-axis order parameters for IL-1 $\beta$ from Sample 1 considered under both tumbling tensor solutions (63 methyls over 37 residues). $O_{axis}^2 = \eta/K\tau_{c,eff}$ ; $K = 3.609 \times 10^9 \text{ s}^{-2}$ , and carries analytic propagation of the $\eta$ error alone. $\sigma$ carries no contribution from tensor uncertainty. $\Delta_{\text{basin}} = |O^2_{\text{A}} - O^2_{\text{B}}|$ acts through the orientation-dependent $\tau_{c,eff}$ and averages 0.0070 and exceeds $\sigma$ at 16 of 63 methyls. $\tau_{c,eff}$ spans 8.6217 to 9.1556 ns under basin A (oblate) and 8.8584 to 9.2235 ns under basin B (prolate), with means of 9.0058 and 9.0057 ns, respectively. The two solutions redistribute $\tau_{c,eff}$ among methyls rather than rescaling it, which is why the basin term is appreciable per site while $\langle O_{axis}^2 \rangle$ moves only from 0.4854 to 0.4862.

**Table S9:** Sample 2 IL-1β 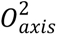 parameters.

| Methyl | basin A (oblate) | | | basin B (prolate) | | | $\Delta_{\text{basin}}$ |
| --- | --- | --- | --- | --- | --- | --- | --- |
| | $O_{axis}^2$ | $\sigma$ | $\tau c, \text{eff (ns)}$ | $O_{axis}^2$ | $\sigma$ | $\tau c, \text{eff (ns)}$ | |
| V3 CG1 | 0.2319 | 0.0025 | 8.6322 | 0.2264 | 0.0025 | 8.8399 | 0.0055 |
| V3 CG2 | 0.2025 | 0.0028 | 8.9007 | 0.1986 | 0.0027 | 9.0757 | 0.0039 |
| L6 CD1 | 0.1836 | 0.0031 | 9.1475 | 0.1823 | 0.0031 | 9.2088 | 0.0013 |
| L6 CD2 | 0.1985 | 0.0034 | 8.7321 | 0.1955 | 0.0033 | 8.8661 | 0.0030 |
| L10 CD1 | 0.6629 | 0.0132 | 9.1439 | 0.6854 | 0.0137 | 8.8437 | 0.0225 |
| L10 CD2 | 0.6197 | 0.0119 | 8.7360 | 0.6073 | 0.0116 | 8.9136 | 0.0124 |
| L18 CD1 | 0.3715 | 0.0055 | 9.1350 | 0.3809 | 0.0057 | 8.9114 | 0.0094 |
| L18 CD2 | 0.3808 | 0.0057 | 9.0632 | 0.3756 | 0.0056 | 9.1887 | 0.0052 |
| V19 CG1 | 0.6219 | 0.0130 | 9.0564 | 0.6371 | 0.0133 | 8.8400 | 0.0152 |
| V19 CG2 | 0.5934 | 0.0091 | 9.1154 | 0.5877 | 0.0090 | 9.2040 | 0.0057 |
| M20 CE | 0.0326 | 0.0044 | 8.7342 | 0.0318 | 0.0043 | 8.9521 | 0.0008 |
| L26 CD1 | 0.4822 | 0.0099 | 8.7649 | 0.4731 | 0.0097 | 8.9334 | 0.0091 |
| L26 CD2 | 0.5903 | 0.0134 | 8.9589 | 0.5950 | 0.0135 | 8.8878 | 0.0047 |
| L29 CD1 | 0.3511 | 0.0055 | 9.1334 | 0.3632 | 0.0057 | 8.8298 | 0.0121 |
| L29 CD2 | 0.3704 | 0.0054 | 8.7191 | 0.3657 | 0.0054 | 8.8323 | 0.0047 |
| L31 CD1 | 0.7026 | 0.0197 | 8.9827 | 0.6960 | 0.0195 | 9.0685 | 0.0066 |
| L31 CD2 | 0.7563 | 0.0114 | 9.1194 | 0.7621 | 0.0115 | 9.0493 | 0.0058 |
| M36 CE | 0.1845 | 0.0036 | 9.0784 | 0.1855 | 0.0036 | 9.0313 | 0.0010 |
| V40 CG1 | 0.2801 | 0.0037 | 9.1165 | 0.2834 | 0.0037 | 9.0110 | 0.0033 |
| V40 CG2 | 0.3014 | 0.0039 | 8.9750 | 0.2978 | 0.0039 | 9.0815 | 0.0036 |
| V41 CG1 | 0.7624 | 0.0188 | 9.0158 | 0.7541 | 0.0186 | 9.1155 | 0.0083 |
| V41 CG2 | 0.8556 | 0.0154 | 9.1484 | 0.8831 | 0.0159 | 8.8637 | 0.0275 |
| M44 CE | 0.3599 | 0.0095 | 9.1472 | 0.3571 | 0.0094 | 9.2188 | 0.0028 |
| V47 CG1 | 0.7488 | 0.0252 | 8.7357 | 0.7408 | 0.0249 | 8.8308 | 0.0080 |
| V47 CG2 | 0.7364 | 0.0182 | 8.9583 | 0.7230 | 0.0179 | 9.1237 | 0.0134 |
| I56 CD1 | 0.2528 | 0.0022 | 9.0658 | 0.2507 | 0.0022 | 9.1405 | 0.0021 |
| V58 CG1 | 0.8992 | 0.0216 | 8.8931 | 0.8959 | 0.0215 | 8.9255 | 0.0033 |
| V58 CG2 | 0.8494 | 0.0161 | 9.0769 | 0.8382 | 0.0159 | 9.1975 | 0.0112 |
| L60 CD1 | 0.5251 | 0.0096 | 9.0202 | 0.5168 | 0.0094 | 9.1656 | 0.0083 |
| L60 CD2 | 0.5724 | 0.0054 | 9.1405 | 0.5884 | 0.0056 | 8.8922 | 0.0160 |
| L62 CD1 | 0.2407 | 0.0037 | 8.8345 | 0.2410 | 0.0037 | 8.8224 | 0.0003 |
| L62 CD2 | 0.2389 | 0.0022 | 8.8614 | 0.2355 | 0.0022 | 8.9874 | 0.0034 |
| L67 CD1 | 0.5341 | 0.0098 | 9.1249 | 0.5431 | 0.0100 | 8.9732 | 0.0090 |
| L67 CD2 | 0.4613 | 0.0076 | 9.1398 | 0.4732 | 0.0078 | 8.9101 | 0.0119 |
| L69 CD1 | 0.2617 | 0.0065 | 8.7424 | 0.2592 | 0.0065 | 8.8267 | 0.0025 |
| L69 CD2 | 0.2443 | 0.0039 | 9.0560 | 0.2420 | 0.0038 | 9.1419 | 0.0023 |
| V72 CG1 | 0.4951 | 0.0082 | 9.0849 | 0.4907 | 0.0082 | 9.1654 | 0.0044 |
| V72 CG2 | 0.4964 | 0.0068 | 9.0808 | 0.5004 | 0.0069 | 9.0080 | 0.0040 |
| L73 CD1 | 0.3008 | 0.0054 | 9.1427 | 0.3114 | 0.0056 | 8.8327 | 0.0106 |
| L73 CD2 | 0.3282 | 0.0043 | 8.8909 | 0.3227 | 0.0042 | 9.0429 | 0.0055 |
| L80 CD1 | 0.5849 | 0.0150 | 9.0306 | 0.5807 | 0.0149 | 9.0957 | 0.0042 |
| L80 CD2 | 0.6523 | 0.0096 | 9.0850 | 0.6696 | 0.0099 | 8.8500 | 0.0173 |
| L82 CD1 | 0.2178 | 0.0038 | 9.0278 | 0.2155 | 0.0037 | 9.1241 | 0.0023 |
| L82 CD2 | 0.1921 | 0.0028 | 8.8540 | 0.1879 | 0.0027 | 9.0533 | 0.0042 |
| V85 CG1 | 0.8420 | 0.0173 | 8.8298 | 0.8237 | 0.0169 | 9.0259 | 0.0183 |
| V85 CG2 | 0.7972 | 0.0118 | 9.1459 | 0.8230 | 0.0122 | 8.8599 | 0.0258 |
| M95 CE | 0.4306 | 0.0120 | 9.1471 | 0.4303 | 0.0120 | 9.1519 | 0.0003 |
| V100 CG1 | 0.9364 | 0.0186 | 8.7030 | 0.9209 | 0.0183 | 8.8497 | 0.0155 |
| V100 CG2 | 0.9088 | 0.0312 | 8.9288 | 0.8929 | 0.0307 | 9.0883 | 0.0159 |
| I104 CD1 | 0.4023 | 0.0082 | 9.1481 | 0.4058 | 0.0083 | 9.0686 | 0.0035 |
| I106 CD1 | 0.2577 | 0.0030 | 9.1479 | 0.2553 | 0.0029 | 9.2328 | 0.0024 |
| L110 CD1 | 0.2265 | 0.0044 | 8.6809 | 0.2208 | 0.0043 | 8.9036 | 0.0057 |
| L110 CD2 | 0.2297 | 0.0033 | 8.9430 | 0.2284 | 0.0033 | 8.9954 | 0.0013 |
| I122 CD1 | 0.7573 | 0.0214 | 9.0491 | 0.7573 | 0.0214 | 9.0492 | 0.0000 |
| M130 CE | 0.1335 | 0.0023 | 9.0861 | 0.1375 | 0.0024 | 8.8258 | 0.0040 |
| V132 CG1 | 0.8739 | 0.0191 | 8.8200 | 0.8625 | 0.0188 | 8.9372 | 0.0114 |
| V132 CG2 | 0.8228 | 0.0249 | 9.1483 | 0.8533 | 0.0258 | 8.8212 | 0.0305 |
| L134 CD1 | 0.6008 | 0.0179 | 9.1463 | 0.6100 | 0.0181 | 9.0080 | 0.0092 |
| L134 CD2 | 0.6891 | 0.0143 | 9.0878 | 0.7044 | 0.0146 | 8.8892 | 0.0153 |
| I143 CD1 | 0.5525 | 0.0151 | 9.0202 | 0.5647 | 0.0155 | 8.8260 | 0.0122 |
| M148 CE | 0.2787 | 0.0096 | 8.5656 | 0.2707 | 0.0094 | 8.8218 | 0.0080 |
| V151 CG1 | 0.3606 | 0.0041 | 9.0416 | 0.3553 | 0.0041 | 9.1757 | 0.0053 |
| V151 CG2 | 0.3715 | 0.0073 | 9.0899 | 0.3825 | 0.0075 | 8.8295 | 0.0110 |

Methyl symmetry-axis order parameters for IL-1 $\beta$ from Sample 2 considered under both tumbling tensor solutions (63 methyls over 37 residues). $O_{axis}^2 = \eta/K\tau_{c,eff}$ ; $K = 3.609 \times 10^9 \text{ s}^{-2}$ , and carries analytic propagation of the $\eta$ error alone. $\sigma$ carries no contribution from tensor uncertainty. $\Delta_{\text{basin}} = |O^2_{\text{A}} - O^2_{\text{B}}|$ acts through the orientation-dependent $\tau_{c,eff}$ and averages 0.0082 and exceeds $\sigma$ at 23 of 63 methyls. $\tau_{c,eff}$ spans 8.5656 to 9.1484 ns under basin A (oblate) and 8.8212 to 9.228 ns under basin B (prolate), with means of 8.9862 and 8.9879 ns, respectively. The two solutions redistribute $\tau_{c,eff}$ among methyls rather than rescaling it, which is why the basin term is appreciable per site while $\langle O_{axis}^2 \rangle$ moves only from 0.4826 to 0.4833.

**Table S10:** Weighted average IL-1β 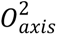 parameters.

|  | basin A (oblate) |  | basin B (prolate) |  |  |
| --- | --- | --- | --- | --- | --- |
| Methyl | $O_{axis}^2$ $\sigma$ | WSE | $O_{axis}^2$ | WSE | $\Delta_{basin}$ |
| V3 CG1 | 0.2325 | 0.0021 | 0.2270 | 0.0021 | 0.0054 |
| V3 CG2 | 0.2037 | 0.0023 | 0.1999 | 0.0022 | 0.0038 |
| L6 CD1 | 0.1864 | 0.0021 | 0.1851 | 0.0021 | 0.0014 |
| L6 CD2 | 0.1976 | 0.0026 | 0.1947 | 0.0025 | 0.0030 |
| L10 CD1 | 0.6466 | 0.0069 | 0.6677 | 0.0071 | 0.0211 |
| L10 CD2 | 0.6168 | 0.0084 | 0.6048 | 0.0082 | 0.0121 |
| L18 CD1 | 0.3703 | 0.0033 | 0.3767 | 0.0033 | 0.0064 |
| L18 CD2 | 0.3799 | 0.0041 | 0.3760 | 0.0040 | 0.0038 |
| V19 CG1 | 0.6251 | 0.0097 | 0.6396 | 0.0100 | 0.0145 |
| V19 CG2 | 0.5936 | 0.0060 | 0.5907 | 0.0060 | 0.0029 |
| M20 CE | 0.0272 | 0.0026 | 0.0267 | 0.0026 | 0.0006 |
| L26 CD1 | 0.4845 | 0.0064 | 0.4753 | 0.0063 | 0.0093 |
| L26 CD2 | 0.5904 | 0.0088 | 0.5976 | 0.0089 | 0.0072 |
| L29 CD1 | 0.3475 | 0.0040 | 0.3591 | 0.0042 | 0.0116 |
| L29 CD2 | 0.3686 | 0.0039 | 0.3637 | 0.0039 | 0.0049 |
| L31 CD1 | 0.7047 | 0.0134 | 0.6994 | 0.0133 | 0.0053 |
| L31 CD2 | 0.7560 | 0.0075 | 0.7573 | 0.0075 | 0.0014 |
| M36 CE | 0.1889 | 0.0024 | 0.1904 | 0.0024 | 0.0015 |
| V40 CG1 | 0.2823 | 0.0029 | 0.2844 | 0.0029 | 0.0021 |
| V40 CG2 | 0.3010 | 0.0033 | 0.2976 | 0.0033 | 0.0034 |
| V41 CG1 | 0.8051 | 0.0112 | 0.7967 | 0.0111 | 0.0084 |
| V41 CG2 | 0.8642 | 0.0090 | 0.8852 | 0.0092 | 0.0210 |
| M44 CE | 0.3639 | 0.0065 | 0.3612 | 0.0064 | 0.0027 |
| V47 CG1 | 0.7501 | 0.0190 | 0.7421 | 0.0188 | 0.0080 |
| V47 CG2 | 0.7385 | 0.0136 | 0.7274 | 0.0135 | 0.0112 |
| I56 CD1 | 0.2548 | 0.0015 | 0.2525 | 0.0015 | 0.0023 |
| V58 CG1 | 0.9060 | 0.0144 | 0.9037 | 0.0144 | 0.0023 |
| V58 CG2 | 0.8474 | 0.0103 | 0.8397 | 0.0102 | 0.0077 |
| L60 CD1 | 0.5256 | 0.0066 | 0.5188 | 0.0066 | 0.0068 |
| L60 CD2 | 0.5724 | 0.0041 | 0.5889 | 0.0043 | 0.0166 |
| L62 CD1 | 0.2412 | 0.0022 | 0.2418 | 0.0022 | 0.0006 |
| L62 CD2 | 0.2401 | 0.0018 | 0.2369 | 0.0018 | 0.0032 |
| L67 CD1 | 0.5367 | 0.0073 | 0.5430 | 0.0074 | 0.0063 |
| L67 CD2 | 0.4607 | 0.0056 | 0.4733 | 0.0057 | 0.0126 |
| L69 CD1 | 0.2651 | 0.0040 | 0.2619 | 0.0039 | 0.0032 |
| L69 CD2 | 0.2454 | 0.0026 | 0.2436 | 0.0025 | 0.0018 |
| V72 CG1 | 0.4999 | 0.0063 | 0.4977 | 0.0063 | 0.0022 |
| V72 CG2 | 0.4984 | 0.0048 | 0.5001 | 0.0048 | 0.0017 |
| L73 CD1 | 0.3013 | 0.0040 | 0.3106 | 0.0042 | 0.0093 |
| L73 CD2 | 0.3277 | 0.0033 | 0.3221 | 0.0032 | 0.0056 |
| L80 CD1 | 0.5903 | 0.0107 | 0.5852 | 0.0106 | 0.0052 |
| L80 CD2 | 0.6516 | 0.0073 | 0.6658 | 0.0075 | 0.0142 |
| L82 CD1 | 0.2168 | 0.0026 | 0.2148 | 0.0025 | 0.0020 |
| L82 CD2 | 0.1919 | 0.0019 | 0.1884 | 0.0019 | 0.0036 |
| V85 CG1 | 0.8339 | 0.0101 | 0.8202 | 0.0099 | 0.0137 |
| V85 CG2 | 0.8025 | 0.0078 | 0.8285 | 0.0081 | 0.0260 |
| M95 CE | 0.4327 | 0.0086 | 0.4312 | 0.0086 | 0.0015 |
| V100 CG1 | 0.9406 | 0.0155 | 0.9251 | 0.0153 | 0.0156 |
| V100 CG2 | 0.8997 | 0.0202 | 0.8845 | 0.0199 | 0.0151 |
| I104 CD1 | 0.4043 | 0.0057 | 0.4099 | 0.0058 | 0.0057 |
| I106 CD1 | 0.2596 | 0.0019 | 0.2579 | 0.0019 | 0.0017 |
| L110 CD1 | 0.2259 | 0.0029 | 0.2211 | 0.0028 | 0.0048 |
| L110 CD2 | 0.2284 | 0.0023 | 0.2265 | 0.0023 | 0.0019 |
| I122 CD1 | 0.7573 | 0.0099 | 0.7643 | 0.0100 | 0.0070 |
| M130 CE | 0.1350 | 0.0013 | 0.1389 | 0.0013 | 0.0039 |
| V132 CG1 | 0.8799 | 0.0139 | 0.8674 | 0.0137 | 0.0125 |
| V132 CG2 | 0.8282 | 0.0164 | 0.8557 | 0.0170 | 0.0276 |
| L134 CD1 | 0.6088 | 0.0123 | 0.6210 | 0.0125 | 0.0121 |
| L134 CD2 | 0.6907 | 0.0109 | 0.7025 | 0.0111 | 0.0118 |
| I143 CD1 | 0.5532 | 0.0103 | 0.5642 | 0.0106 | 0.0109 |
| M148 CE | 0.2775 | 0.0071 | 0.2698 | 0.0069 | 0.0077 |
| V151 CG1 | 0.3607 | 0.0033 | 0.3559 | 0.0033 | 0.0048 |
| V151 CG2 | 0.3725 | 0.0053 | 0.3817 | 0.0054 | 0.0093 |

Summary of replicates from the two independently prepared samples IL-1 $\beta$ . Data gathered from SI Tables S8-S9 (63 methyls over 37 residues). Samples are combined by inverse-variance weighting on the raw analytic $\sigma$ (See Methods). $\Delta_{\text{basin}} = |\langle O^2 \rangle_A - \langle O^2 \rangle_B|$ is the effect of the undetermined anisotropy symmetry, reported with the weighed standard error (WSE) but not combined with it. The WSE averages 0.0075 and exceeds the WSE at 32 of 63 methyls (51%) — the two preparations agree so closely (mean WSE 0.0068) that the basin term, not the fitting error, dominates at individual sites. The $\langle O_{axis}^2 \rangle$ is 0.4840 under basin A (oblate) and 0.4848 under basin B (prolate), a shift of 0.0008; the mean WSE is 0.0068 and 0.0068, respectively i.e., the basins are equally well determined.

**Table S11:**
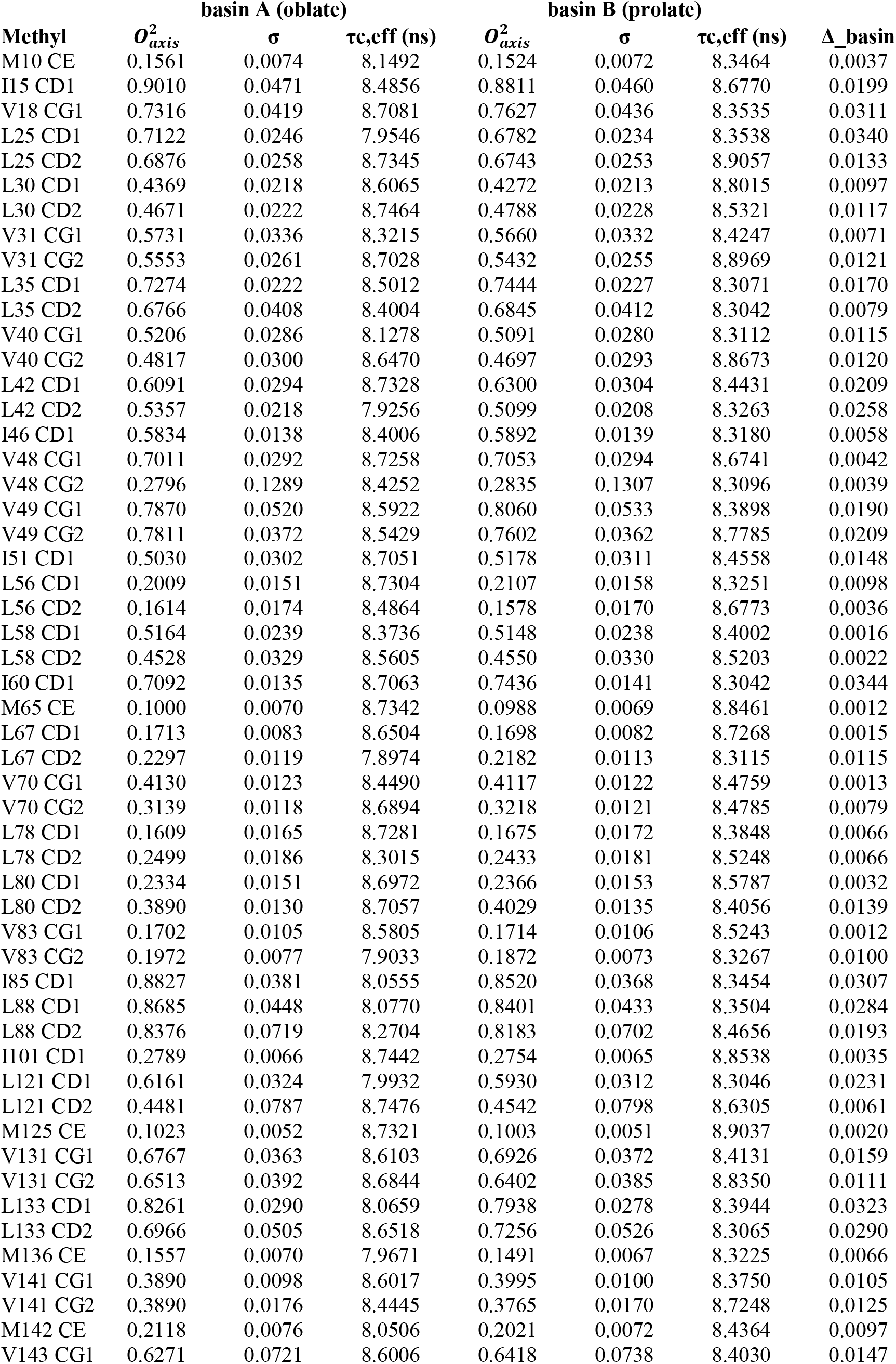
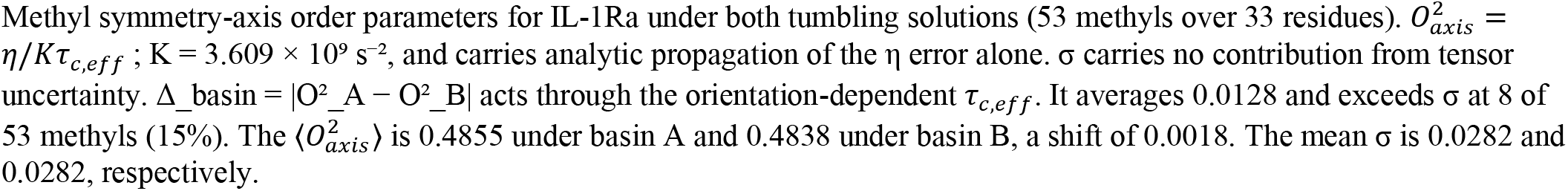
IL-1Ra 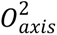 parameters.

| Methyl | basin A (oblate) | | | basin B (prolate) | | | $\Delta_{\text{basin}}$ |
| --- | --- | --- | --- | --- | --- | --- | --- |
| | $O_{axis}^2$ | $\sigma$ | $\tau c, \text{eff (ns)}$ | $O_{axis}^2$ | $\sigma$ | $\tau c, \text{eff (ns)}$ | |
| M10 CE | 0.1561 | 0.0074 | 8.1492 | 0.1524 | 0.0072 | 8.3464 | 0.0037 |
| I15 CD1 | 0.9010 | 0.0471 | 8.4856 | 0.8811 | 0.0460 | 8.6770 | 0.0199 |
| V18 CG1 | 0.7316 | 0.0419 | 8.7081 | 0.7627 | 0.0436 | 8.3535 | 0.0311 |
| L25 CD1 | 0.7122 | 0.0246 | 7.9546 | 0.6782 | 0.0234 | 8.3538 | 0.0340 |
| L25 CD2 | 0.6876 | 0.0258 | 8.7345 | 0.6743 | 0.0253 | 8.9057 | 0.0133 |
| L30 CD1 | 0.4369 | 0.0218 | 8.6065 | 0.4272 | 0.0213 | 8.8015 | 0.0097 |
| L30 CD2 | 0.4671 | 0.0222 | 8.7464 | 0.4788 | 0.0228 | 8.5321 | 0.0117 |
| V31 CG1 | 0.5731 | 0.0336 | 8.3215 | 0.5660 | 0.0332 | 8.4247 | 0.0071 |
| V31 CG2 | 0.5553 | 0.0261 | 8.7028 | 0.5432 | 0.0255 | 8.8969 | 0.0121 |
| L35 CD1 | 0.7274 | 0.0222 | 8.5012 | 0.7444 | 0.0227 | 8.3071 | 0.0170 |
| L35 CD2 | 0.6766 | 0.0408 | 8.4004 | 0.6845 | 0.0412 | 8.3042 | 0.0079 |
| V40 CG1 | 0.5206 | 0.0286 | 8.1278 | 0.5091 | 0.0280 | 8.3112 | 0.0115 |
| V40 CG2 | 0.4817 | 0.0300 | 8.6470 | 0.4697 | 0.0293 | 8.8673 | 0.0120 |
| L42 CD1 | 0.6091 | 0.0294 | 8.7328 | 0.6300 | 0.0304 | 8.4431 | 0.0209 |
| L42 CD2 | 0.5357 | 0.0218 | 7.9256 | 0.5099 | 0.0208 | 8.3263 | 0.0258 |
| I46 CD1 | 0.5834 | 0.0138 | 8.4006 | 0.5892 | 0.0139 | 8.3180 | 0.0058 |
| V48 CG1 | 0.7011 | 0.0292 | 8.7258 | 0.7053 | 0.0294 | 8.6741 | 0.0042 |
| V48 CG2 | 0.2796 | 0.1289 | 8.4252 | 0.2835 | 0.1307 | 8.3096 | 0.0039 |
| V49 CG1 | 0.7870 | 0.0520 | 8.5922 | 0.8060 | 0.0533 | 8.3898 | 0.0190 |
| V49 CG2 | 0.7811 | 0.0372 | 8.5429 | 0.7602 | 0.0362 | 8.7785 | 0.0209 |
| I51 CD1 | 0.5030 | 0.0302 | 8.7051 | 0.5178 | 0.0311 | 8.4558 | 0.0148 |
| L56 CD1 | 0.2009 | 0.0151 | 8.7304 | 0.2107 | 0.0158 | 8.3251 | 0.0098 |
| L56 CD2 | 0.1614 | 0.0174 | 8.4864 | 0.1578 | 0.0170 | 8.6773 | 0.0036 |
| L58 CD1 | 0.5164 | 0.0239 | 8.3736 | 0.5148 | 0.0238 | 8.4002 | 0.0016 |
| L58 CD2 | 0.4528 | 0.0329 | 8.5605 | 0.4550 | 0.0330 | 8.5203 | 0.0022 |
| I60 CD1 | 0.7092 | 0.0135 | 8.7063 | 0.7436 | 0.0141 | 8.3042 | 0.0344 |
| M65 CE | 0.1000 | 0.0070 | 8.7342 | 0.0988 | 0.0069 | 8.8461 | 0.0012 |
| L67 CD1 | 0.1713 | 0.0083 | 8.6504 | 0.1698 | 0.0082 | 8.7268 | 0.0015 |
| L67 CD2 | 0.2297 | 0.0119 | 7.8974 | 0.2182 | 0.0113 | 8.3115 | 0.0115 |
| V70 CG1 | 0.4130 | 0.0123 | 8.4490 | 0.4117 | 0.0122 | 8.4759 | 0.0013 |
| V70 CG2 | 0.3139 | 0.0118 | 8.6894 | 0.3218 | 0.0121 | 8.4785 | 0.0079 |
| L78 CD1 | 0.1609 | 0.0165 | 8.7281 | 0.1675 | 0.0172 | 8.3848 | 0.0066 |
| L78 CD2 | 0.2499 | 0.0186 | 8.3015 | 0.2433 | 0.0181 | 8.5248 | 0.0066 |
| L80 CD1 | 0.2334 | 0.0151 | 8.6972 | 0.2366 | 0.0153 | 8.5787 | 0.0032 |
| L80 CD2 | 0.3890 | 0.0130 | 8.7057 | 0.4029 | 0.0135 | 8.4056 | 0.0139 |
| V83 CG1 | 0.1702 | 0.0105 | 8.5805 | 0.1714 | 0.0106 | 8.5243 | 0.0012 |
| V83 CG2 | 0.1972 | 0.0077 | 7.9033 | 0.1872 | 0.0073 | 8.3267 | 0.0100 |
| I85 CD1 | 0.8827 | 0.0381 | 8.0555 | 0.8520 | 0.0368 | 8.3454 | 0.0307 |
| L88 CD1 | 0.8685 | 0.0448 | 8.0770 | 0.8401 | 0.0433 | 8.3504 | 0.0284 |
| L88 CD2 | 0.8376 | 0.0719 | 8.2704 | 0.8183 | 0.0702 | 8.4656 | 0.0193 |
| I101 CD1 | 0.2789 | 0.0066 | 8.7442 | 0.2754 | 0.0065 | 8.8538 | 0.0035 |
| L121 CD1 | 0.6161 | 0.0324 | 7.9932 | 0.5930 | 0.0312 | 8.3046 | 0.0231 |
| L121 CD2 | 0.4481 | 0.0787 | 8.7476 | 0.4542 | 0.0798 | 8.6305 | 0.0061 |
| M125 CE | 0.1023 | 0.0052 | 8.7321 | 0.1003 | 0.0051 | 8.9037 | 0.0020 |
| V131 CG1 | 0.6767 | 0.0363 | 8.6103 | 0.6926 | 0.0372 | 8.4131 | 0.0159 |
| V131 CG2 | 0.6513 | 0.0392 | 8.6844 | 0.6402 | 0.0385 | 8.8350 | 0.0111 |
| L133 CD1 | 0.8261 | 0.0290 | 8.0659 | 0.7938 | 0.0278 | 8.3944 | 0.0323 |
| L133 CD2 | 0.6966 | 0.0505 | 8.6518 | 0.7256 | 0.0526 | 8.3065 | 0.0290 |
| M136 CE | 0.1557 | 0.0070 | 7.9671 | 0.1491 | 0.0067 | 8.3225 | 0.0066 |
| V141 CG1 | 0.3890 | 0.0098 | 8.6017 | 0.3995 | 0.0100 | 8.3750 | 0.0105 |
| V141 CG2 | 0.3890 | 0.0176 | 8.4445 | 0.3765 | 0.0170 | 8.7248 | 0.0125 |
| M142 CE | 0.2118 | 0.0076 | 8.0506 | 0.2021 | 0.0072 | 8.4364 | 0.0097 |
| V143 CG1 | 0.6271 | 0.0721 | 8.6006 | 0.6418 | 0.0738 | 8.4030 | 0.0147 |

Methyl symmetry-axis order parameters for IL-1Ra under both tumbling solutions (53 methyls over 33 residues). $O_{axis}^2 = \eta / K \tau_{c,eff}$ ; $K = 3.609 \times 10^9 \text{ s}^{-2}$ , and carries analytic propagation of the $\eta$ error alone. $\sigma$ carries no contribution from tensor uncertainty. $\Delta\_basin = |O^2\_A - O^2\_B|$ acts through the orientation-dependent $\tau_{c,eff}$ . It averages 0.0128 and exceeds $\sigma$ at 8 of 53 methyls (15%). The $\langle O_{axis}^2 \rangle$ is 0.4855 under basin A and 0.4838 under basin B, a shift of 0.0018. The mean $\sigma$ is 0.0282 and 0.0282, respectively.

